# An integrated view of the structure and function of the human 4D nucleome

**DOI:** 10.1101/2024.09.17.613111

**Authors:** 4D Nucleome Consortium, Job Dekker, Betul Akgol Oksuz, Yang Zhang, Ye Wang, Miriam K. Minsk, Shuzhen Kuang, Liyan Yang, Johan H. Gibcus, Nils Krietenstein, Oliver J. Rando, Jie Xu, Derek H. Janssens, Steven Henikoff, Alexander Kukalev, Andréa Willemin, Warren Winick-Ng, Rieke Kempfer, Ana Pombo, Miao Yu, Pradeep Kumar, Liguo Zhang, Andrew S Belmont, Takayo Sasaki, Tom van Schaik, Laura Brueckner, Daan Peric-Hupkes, Bas van Steensel, Ping Wang, Haoxi Chai, Minji Kim, Yijun Ruan, Ran Zhang, Sofia A. Quinodoz, Prashant Bhat, Mitchell Guttman, Wenxin Zhao, Shu Chien, Yuan Liu, Sergey V. Venev, Dariusz Plewczynski, Ibai Irastorza Azcarate, Dominik Szabó, Christoph J. Thieme, Teresa Szczepińska, Mateusz Chiliński, Kaustav Sengupta, Mattia Conte, Andrea Esposito, Alex Abraham, Ruochi Zhang, Yuchuan Wang, Xingzhao Wen, Qiuyang Wu, Yang Yang, Jie Liu, Lorenzo Boninsegna, Asli Yildirim, Yuxiang Zhan, Andrea Maria Chiariello, Simona Bianco, Lindsay Lee, Ming Hu, Yun Li, R. Jordan Barnett, Ashley L. Cook, Daniel J. Emerson, Claire Marchal, Peiyao Zhao, Peter Park, Burak H. Alver, Andrew Schroeder, Rahi Navelkar, Clara Bakker, William Ronchetti, Shannon Ehmsen, Alexander Veit, Nils Gehlenborg, Ting Wang, Daofeng Li, Xiaotao Wang, Mario Nicodemi, Bing Ren, Sheng Zhong, Jennifer E. Phillips-Cremins, David M. Gilbert, Katherine S. Pollard, Frank Alber, Jian Ma, William S. Noble, Feng Yue

## Abstract

The dynamic three-dimensional (3D) organization of the human genome (the “4D Nucleome”) is closely linked to genome function. Here, we integrate a wide variety of genomic data generated by the 4D Nucleome Project to provide a detailed view of human 3D genome organization in widely used embryonic stem cells (H1-hESCs) and immortalized fibroblasts (HFFc6). We provide extensive benchmarking of 3D genome mapping assays and integrate these diverse datasets to annotate spatial genomic features across scales. The data reveal a rich complexity of chromatin domains and their sub-nuclear positions, and over one hundred thousand structural loops and promoter-enhancer interactions. We developed 3D models of population-based and individual cell-to-cell variation in genome structure, establishing connections between chromosome folding, nuclear organization, chromatin looping, gene transcription, and DNA replication. We demonstrate the use of computational methods to predict genome folding from DNA sequence, uncovering potential effects of genetic variants on genome structure and function. Together, this comprehensive analysis contributes insights into human genome organization and enhances our understanding of connections between the regulation of genome function and 3D genome organization in general.

## Introduction

Since the publication of the first draft sequence of the human genome over two decades ago, massive efforts have focused on identifying all genes and functional elements encoded in the genome. The resulting encyclopedia of annotations has revealed a vast richness of coding and regulatory information, leading to an increased understanding of gene regulation in a multitude of cell types and conditions across human development and physiology ^1,2^. Integration of functional annotations with genetic variation mapped at an increasing scale throughout populations is starting to link genetically encoded functional elements and genes to complex traits and human diseases.

Genomes are physical objects and not simply abstract linear databases of stored genetic information. In the case of the human genome, 46 chromosomes are intricately organized as physical objects in three dimensions inside the cell nucleus, while retaining the ability to change and adapt to cell state transitions and cell lineage commitment. It is increasingly appreciated that the spatial organization of genomes is tightly linked to how genetic information encoded within them is activated, utilized, and expressed in a cell type and condition–dependent manner. For instance, enhancers functionally interact with specific distal genes, while ignoring others, through a process that can be controlled by genetic sequences such as insulator elements and tethering elements. This may involve biophysical mechanisms including phase separation, direct enhancer-promoter contacts, loop extrusion by cohesin, condensins, and possibly other folding machines, as well as possible “action at a distance” mechanisms involving diffusion and/or DNA tracking factors ^3–7^.

The genome is organized at different scales ^8–12^. At the local scale of the chromatin fiber, nucleosome positioning and histone modifications influence the structure and accessibility of DNA. At the scale of up to hundreds of kilobases, chromatin loops form in a dynamic fashion, sometimes enriched near specific cis-elements, and in many, but not all, cases such loops are generated through active loop extrusion by cohesin and condensin complexes ^13^. The pattern of extrusion along chromosomes is modulated by cis-elements such as enhancers, promoters and insulators ^14–16^. The process of loop extrusion contributes not only to loops between specific cis-elements including CTCF-bound sites, but it also underlies the formation of many topologically associating domains (TADs; ^17–19^). Loci within TADs interact frequently through cohesin-mediated extrusion ^20^. TADs often have CTCF sites at their boundaries that block extrusion ^18,21,22^, thereby lowering the probability of interaction between loci on either side of the boundary, a phenomenon referred to as insulation ^23^. Finally, chromosomal domains that can range in size from several kilobases to megabases cluster together in space to form sub-nuclear “compartments” ^24–26^. Such associations can involve functionally distinct sub-nuclear structures and bodies such as nuclear speckles, nucleoli, and the nuclear periphery. Many studies over the last several years have started to describe these phenomena, exploring the mechanisms of their formation and their potential roles in genome regulation.

To understand how genomes work to process genetic information into biologically meaningful responses, it is critical to quantitatively map and mechanistically understand the physical organization of the genome relative to itself and to nuclear landmarks and bodies, e.g., identifying which distal enhancers contact target genes and how they work together to regulate gene expression.

The goal of the 4D Nucleome project is to gain detailed insights into the three-dimensional folding of the human genome at the resolution of functional elements, in different cell states, over time, and in single cells (i.e., to map the “4D nucleome”) so that links between chromosome folding and genomic function can be derived, mechanisms of folding can be explored, and causal relations between genome structure and function can be deduced ^27–29^. During its first phase, starting in 2015, a major focus of the project has been the development and benchmarking of complementary experimental approaches for measuring the 4D nucleome, the development of computational and modeling approaches to analyze and interpret 4D nucleome data, and the generation of structural and quantitative models of the folded human genome (Figure 1). We have collected data on chromatin state, chromosome folding, and nuclear organization for two defined human cell types – embryonic stem cells (H1-hESC) and immortalized foreskin fibroblasts (HFFc6), as well as multiple secondary cell lines. Datasets were integrated to obtain linear annotations of 4D nucleome features along chromosomes, and to build 3D genome models, including models that reflect cell-to-cell variation in genome organization. Genome models and structural features were used to gain insights into how chromosome structure relates to gene expression and DNA replication patterns, and to build predictive models that can infer effects of sequence variants on chromosome folding, e.g., in disease. Here, we describe the generation and analysis of genomic data types representing different aspects of the 4D nucleome in the two cell types, in cell populations, and in single cells. All data described in this work are publicly available at the 4DN Data Coordination and Integration Center (https://data.4dnucleome.org/).

Here we present a detailed analysis and integration of these data to:

● Benchmark and validate genomic assays for detecting and quantifying distinct features of chromosome folding
● Integrate data obtained with diverse methods to produce genome-scale annotations of spatial features of loci along the linear genome
● Generate ensembles of 3D genome models representing population-based folding states and cell-to-cell variation in spatial organization
● Relate genome folding to functional processes such as transcription and replication
● Train computational methods to predict 3D genome folding from DNA sequence
● Start to apply insights obtained from these 4D nucleome studies to identify roles of specific DNA elements in chromosome folding, and predict impact of disease-related variants on genome folding and function

## Results

### Benchmarking genomic assays for mapping genome folding

A growing number of assays are being developed to probe the three-dimensional folding of genomes. Here we present the generation and analysis of data obtained with sequencing-based assays that were the main focus of the first phase of the 4DN project, while ongoing and future analyses of the consortium place emphasis on imaging-based assays. Sequencing-based assays can be divided in two broad classes (Figure 1). The first relies on chromatin interaction assays that comprehensively detect spatial proximities between loci, i.e., the interaction frequencies between pairs or among sets of associated loci (e.g., 3C-based assays). The second set of approaches report on physical distances of loci (i.e., TSA-seq, multiplexed immunoFISH) or contact frequencies (i.e., DamID, SPRITE) to specific nuclear structures, such as the lamina, nucleoli, nuclear speckles, etc. We started by comparingthe most used chromatin interaction assays and integrating with assays that report sub-nuclear locus positions to gain insights into how folding and nuclear placement relates to genome function.

**Figure 1.**
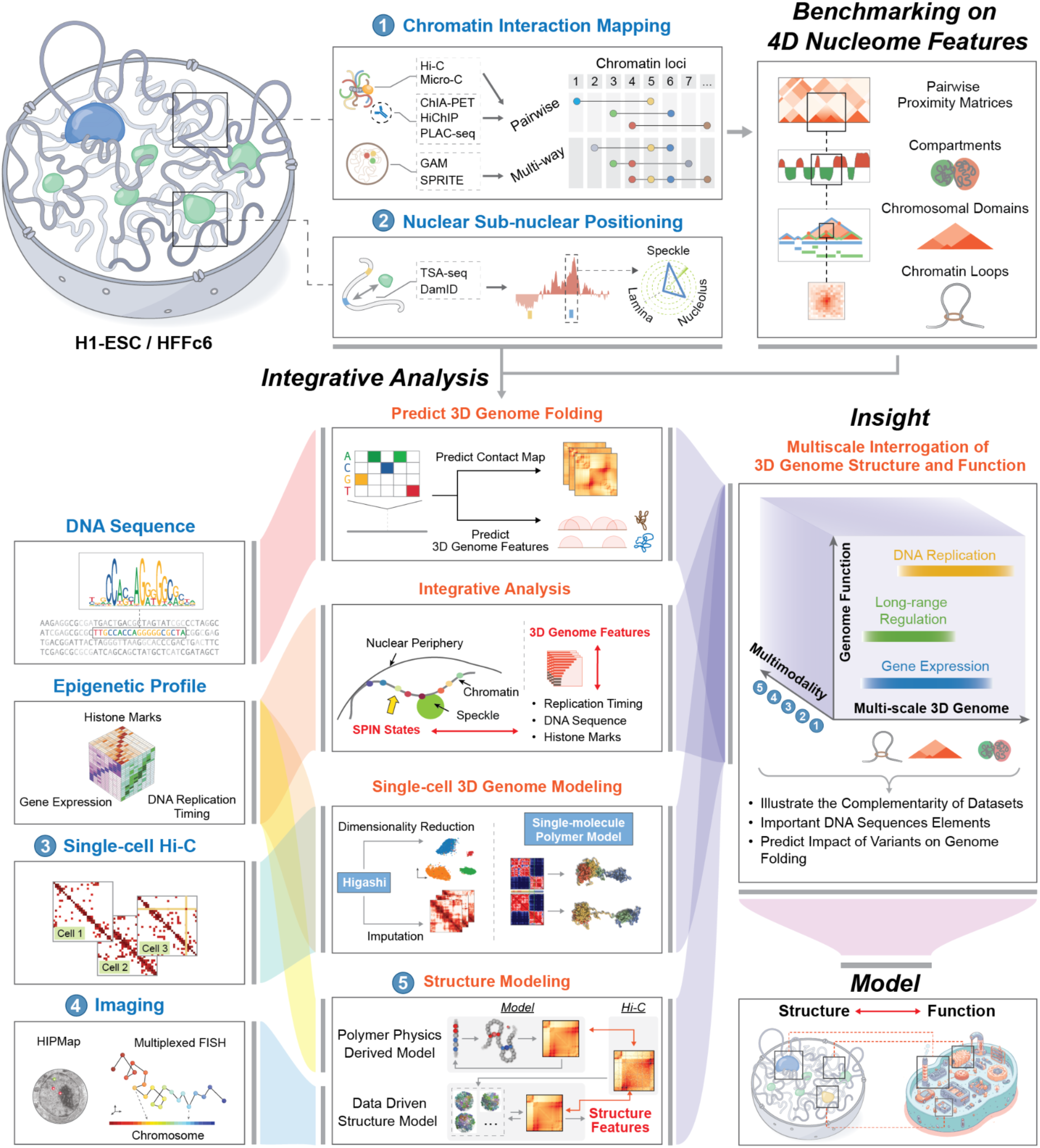
Overview of the key highlights from the first phase of 4D Nucleome project. **(Top-left)** Schematic plots illustrate two types of complementary genomic assays for mapping 3D genome folding and the relative distances of genomic loci to nuclear bodies in H1-hESC and HFFc6 cells. **(Top-right)** Different chromatin interaction mapping methods are compared and benchmarked to assess their ability to identify and quantify 3D genome features at scales ranging from chromatin compartments (Mb) to focally enriched chromatin interactions (kb). **(Bottom-left)** Additional multimodal datasets generated or utilized to facilitate integrative analyses (see below). **(Bottom-center)** Multiple

Sequencing-based chromatin interaction detection methods differ in important ways, e.g., detecting pair-wise contact frequencies as in Hi-C vs. sets of spatially proximal loci as in GAM or SPRITE. These methods can be unbiased in that they identify spatially proximal loci independent of specific factors (e.g., chromosome conformation capture-based assays ^30^ such as Hi-C ^24^ and Micro-C ^31,32^, SPRITE ^33^, or GAM ^34^), or are tailored to identify interactions between loci associated with specific proteins (e.g., ChIA-PET ^35^ and HiChIP/PLAC-seq ^36,37^).

Here we compared the different methods for their ability to determine and quantify 4D nucleome features, including chromatin interactions (i.e., pairwise proximity matrices), chromatin loops, chromosomal domains and compartments, and the sub-nuclear positions of loci in H1-hESC and HFFc6 cells. Data was compared using two concordant biological replicates obtained using unbiased genome-wide approaches (Hi-C, Micro-C, and SPRITE, and GAM) and targeted approaches (ChIA-PET for RNA Polymerase II and CTCF, and PLAC-seq for H3K4me3) (Supplemental Figure 1). We also refer to two recent comprehensive studies benchmarking GAM against Hi-C ^38^, and Micro-C and Hi-C protocol variants ^39^. Those studies showed that GAM and Hi-C, and Micro-C and Hi-C quantitatively differ in compartment detection and loop detection. We also refer to a recent study where polymer models of chromatin 3D architecture were used to show that Hi-C, GAM, and SPRITE bulk data all capture overall reference 3D structures, while single-cell data can reflect the strong variability among single DNA molecules40.

One measure of data quality is the fraction of interactions that are intra-chromosomal (*cis*) vs. inter-chromosomal (*trans*). For all datasets (except GAM, where such metric cannot be directly obtained), the level of cis interactions was 70-90% (Supplemental Figure 1c), indicating high signal-to-noise ratios as random ligation events are thought to enrich inter-chromosomal interactions (*trans*), and an absence of true signal is expected to produce only 2-5% cis interactions. Datasets clustered (based on compartment and insulation profiles) first by cell type and then by method. SPRITE and GAM, being the only multi-way interaction detection methods, each clustered as a separate group (Figure 2b,d).

**Figure 2.**
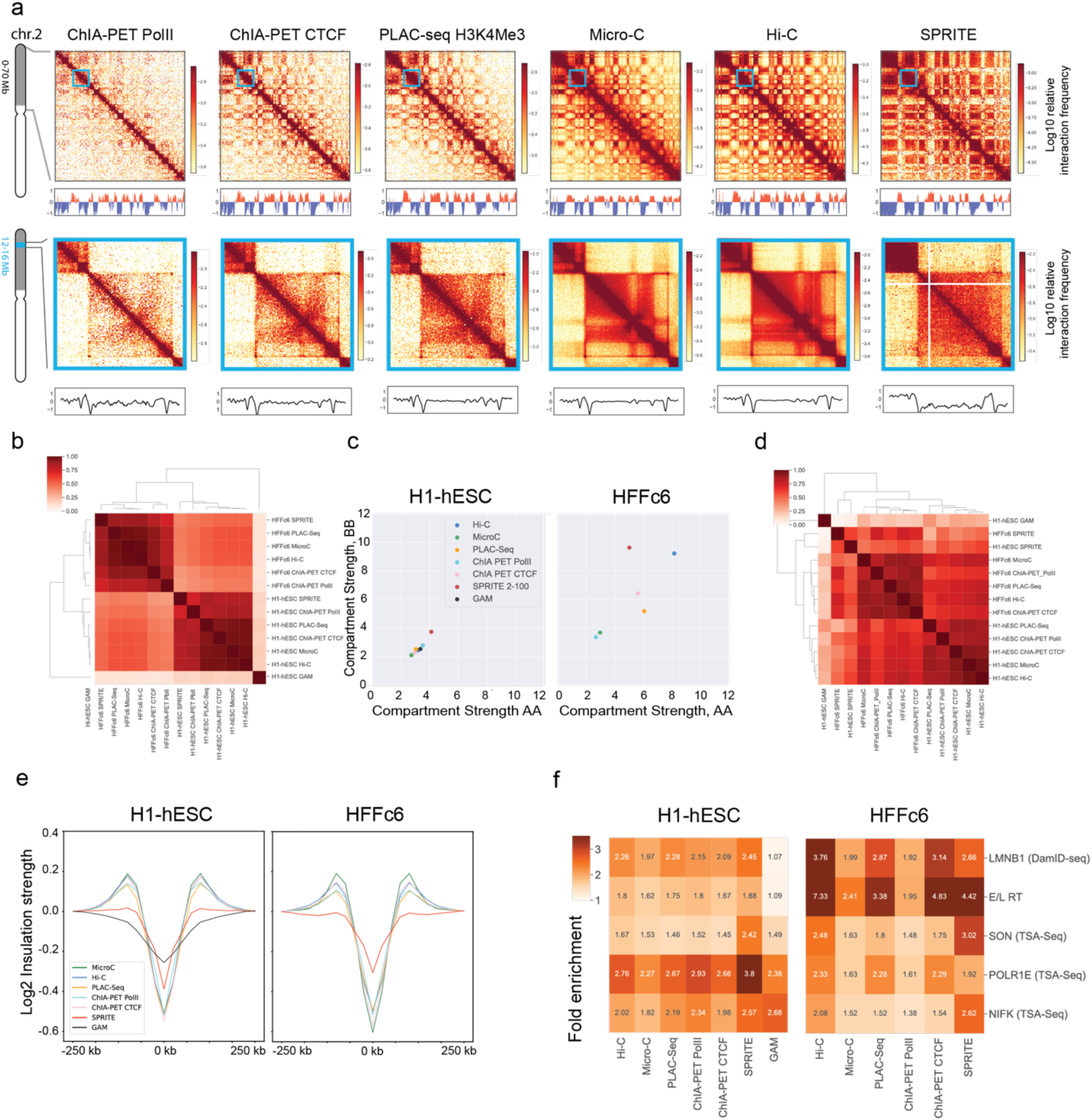
Methods for chromatin interaction detection differ in quantitative detection of compartmentalization. **a.** (Upper panel) Heatmaps of contact maps generated using Hi-C, Micro-C, ChIA-PET, PLAC-Seq, and SPRITE (100 kb bins, chr. 2 0-70 Mb) derived from HFFc6 cells. (Lower Panel) Zoomed heatmaps of contact maps (25 kb bins, chr2 12-16mb) **b.** Spearman correlation of compartment profiles determined by Eigenvector decomposition (see Supplemental Methods). **c.** Compartment strength quantified using eigenvectors detected and quantified from contact data obtained with corresponding 3D methods **d.** Pearson correlation of genome-wide insulation scores for all methods. **e.** Aggregated insulation scores at strong boundaries detected in multiple datasets (see Supplemental Methods), using data obtained from indicated methods. **f.** Preferential interactions quantified in Hi-C, Micro-C, ChIA-PET, PLAC Seq, SPRITE and GAM, using DamID Seq for Lamin B, Early and Late replication timing (E/L RT) using RepliSeq, and TSA Seq for SON to rank loci: The fold enrichment indicates the preference of loci with similar associations with speckles (SON), nucleoli (PolR1E/NFIK), lamina (Lamin B), or that display early or late replication to interact with each other, as detected by the indicated interaction assays.

#### Comparative analysis of chromatin interaction frequencies

To visualize relative interaction frequencies (“contacts”) between loci, contact maps were plotted as two-dimensional heatmaps at different length scales (Figure 2a, data for HFFc6; Supplemental Figure 1a for H1-hESC). Visual inspection of these contact maps at large genomic distances show that ligation-based methods capture similar chromatin organization patterns, independently of whether contacts are mapped with Hi-C, Micro-C, GAM, SPRITE, or based on occupancy of specific proteins (CTCF, RNA Pol II, H3K4Me3). Zooming in on specific genomic regions shows that mapping of chromatin contacts enriched for CTCF, RNA Pol II or the H3K4me3 histone mark captures subsets of contacts seen as a composite in Micro-C and Hi-C. SPRITE detects sets of spatially proximal genomic loci, ranging from clusters of 2 loci up to thousands of loci. To visualize SPRITE data in Figure 2, we converted all clusters into weighted pairwise interactions exactly as described ^33^. For most subsequent analyses described below, SPRITE data was split in subsets of interactions dependent on cluster size.

To quantitatively compare the contacts captured by each method, we computed interaction frequencies *P* as a function of genomic distance (*s*) for HFFc6 cells (Supplemental Figure 1e; similar results were obtained for H1-hESC cells). For all methods, the expected inverse relationship between interaction frequency and genomic distance was observed. The shape of the *P*(*s*) plots is comparable for all datasets, as indicated by the derivative of *P*(*s*). *P*(*s*) of all datasets revealed the presence of loops of around 100 kb as visible by the characteristic “bump” in the *P*(*s*) plot around 100-200 kb ^41^. This characteristic bump has been ascribed to the presence of cohesin-mediated loops. It is noteworthy that such global features of chromosome folding are also detected with ChIA-PET and PLAC-seq that were targeted to enrich for interactions involving sites occupied by CTCF, RNA Pol II, or H3K4me3.

However, the methods differ in dynamic range, with Micro-C having the largest dynamic range and SPRITE (all cluster sizes combined) and GAM (see ^38^) the smallest. We further explored SPRITE data split by cluster size. For small clusters (2-100 fragments), *P*(*s*) is steeper, and the dynamic range approaches that observed with Hi-C. For larger cluster sizes (100-1,000 and 1,000-10,000 fragments), *P*(*s*) became increasingly flat, and *trans* interactions increased greatly (Supplemental Figure 1f-h). Thus, SPRITE clusters of increasing sizes represent increasingly larger chromosome structures, ranging from local pair-wise structures for small clusters to large sub-nuclear structures containing sections from multiple chromosomes for the largest clusters. This is illustrated in more detail below.

#### Methods differ in detection and quantification of chromosomal compartments

Genomes are generally spatially segregated into active A and inactive B compartments that correlate with euchromatin and heterochromatin, respectively ^24,42^. A and B compartments can be further split into subcompartments with distinct chromatin states and interaction profiles (^21,43,44^, and below). Compartmentalization is readily visible in all contact maps shown in Figure 2a (and Supplemental Figure 1a) as a plaid pattern of enriched interactions between domains of the same type, and depleted interactions of domains of different types. We used eigenvector decomposition and found that A and B compartmentalization is typically captured in the first eigenvectors (as shown before ^24^), which were highly correlated for most assays (Spearman coefficient > 0.73; Figure 2b), and clustered according to cell type. GAM eigenvectors correlated with lower Spearman coefficients.

Compartmentalization strength can be calculated by ranking loci based on the value of the first eigenvector along both axes of an interaction map to produce a “saddle plot” ^39,45^. In such plots, B-B interactions cluster in the top left, and A-A interaction in the bottom right.

Compartmentalization strength is then calculated for A and B compartments separately as the ratio of the B-B or A-A interaction scores and the level of A-B interaction scores. We find that in H1-hESCs compartmentalization is relatively weak, in all methods, in comparison to the terminally differentiated HFFc6 fibroblast cells, as reported before ^39,46^. Interestingly, different compartment strengths were found with each method for HFFc6 cells: The strongest compartmentalization was found with SPRITE data obtained from clusters containing 2-100 fragments, and with Hi-C (Figure 2c). Compartmentalization detected with GAM, Micro-C, and the targeted assays was considerably weaker.

We also explored the contribution of larger SPRITE clusters to the detection of compartmentalization. We find that inclusion of larger SPRITE clusters results in decreased compartmentalization strength in both H1-hESC and HFFc6 cells, and loss of the smaller compartment domains due to becoming absorbed into flanking domains (Extended Data Figure 1a,b,c). Comparing the distributions of compartment domain sizes as detected by all methods, we find that GAM and ChIA-PET detect the smallest domains (for GAM: 80% of domains are <1 Mb), while for data obtained with most other assays only 50% of domains are smaller than 1 Mb; Extended Data Figure 1e-f). However, compartmentalization is the strongest when calculated with data obtained with relatively small SPRITE clusters (2-100 fragments) (Figure 2c).

Cytologically, compartmentalization is related to the preferential co-localization of sets of loci at preferred sub-nuclear locations, or around specific sub-nuclear bodies ^47,48^. For instance, B compartment domains are often located near the nuclear and/or nucleolar periphery and are late replicating ^49^. Such domains include Lamin Associated Domains (LADs) and these have been shown to colocalize by Hi-C ^50^. In contrast, A compartments and gene dense chromatin in general are located within the nuclear interior, are earlier replicating ^49^, and enriched for active genes ^51–53^. A subset of A compartment domains contains genomic regions with high gene expression that are preferentially positioned near nuclear speckles and are early replicating ^33,54^. This allowed us to assess, and validate, performance of each of the chromatin interaction assays to detect compartmentalization by using orthogonal data sets representing independent measures of sub-nuclear compartments. For H1-hESCs and HFFc6, we generated genome-wide maps of LADs using Lamin B1 DamID ^55^, of speckle-associated domains using SON TSA-Seq ^56^, of nucleolar associated domains using POLR1E TSA-seq and NFIK TSA-seq ^57^, and determined replication timing using Repli-Seq ^58^. We calculated the extent to which preferential interactions between loci of the same type are detected with each interaction method (Figure 2f). We find compartmentalization calculated in this way is again stronger in HFFc6 cells as compared to H1-hESC cells. SPRITE (2-100 cluster size) and Hi-C generally detect the strongest homotypic associations. Micro-C, ChIA-PET and PLAC-seq also detected such preferentially homotypic interactions, but these preferences appeared weaker. GAM and SPRITE detected relatively strong associations between loci associated with the nucleolus. These assays may be able capture interactions with a larger contact radius, which may contribute to their ability to detect co-association of loci at and around larger subnuclear structures such as speckles and nucleoli. Interestingly, interactions between loci with similar replication timing is observed with all assays, consistent with earlier reports that early and late replication domains correlate strongly with A and B compartments detected by Hi-C ^59^. In HFFc6 cells this correlation between replication timing and interaction frequency is much higher than in H1-hESC cells, consistent with the previous demonstration that consolidation of replication domains occurs during hESC differentiation coincident with a progressive increase in the alignment of replication timing with A / B Hi-C compartments ^46,59,60^.

Together, these observations show that all methods can be used to qualitatively detect compartmentalization and to identify compartment domains (Extended Data Figure 1b). However, quantitative differences between the methods are large, both in terms of the size of compartment domains, the ability to detect smaller compartment domains, and the ability to quantify the strength of compartmentalization.

#### Detection of TAD boundaries

TAD boundaries can reduce the probability of interactions between cis-regulatory elements and genes located on either side of the boundary, and therefore there is a great interest in identifying their genomic locations and characteristics. To measure such boundaries, we performed insulation analysis ^23^. First, we found that insulation score profiles were visually very similar for data obtained with the different methods, although for SPRITE where the sequencing was not high depth, and GAM data the profile has a reduced dynamic range (examples in Figure 2a insets, and Figure 2e). This result was confirmed by calculating and clustering genome-wide Pearson correlation values of insulation scores: insulation profiles for data obtained with all methods were highly correlated (For all assays but SPRITE and GAM: *r* > 0.76 for H1-hESC and *r* > 0.8 for HFFc6); SPRITE and GAM-derived insulation scores had lower correlation values with other data sets (SPRITE *r* = 0.28-0.75; GAM: r = 0.19-0.36; Figure 2d). Insulation profiles are generally clustered by cell type, except SPRITE and GAM data. Finally, insulation scores aggregated at a set of boundaries identified in multiple datasets (see Supplemental Methods) showed that all methods detected boundary strength in comparable ways, except for SPRITE and GAM where boundary strength appeared weaker (Figure 2e). In summary, local domain boundary formation is a robustly detected feature of genome folding that is captured by a variety of chromatin interaction assays.

**Extended Data Figure 1.**
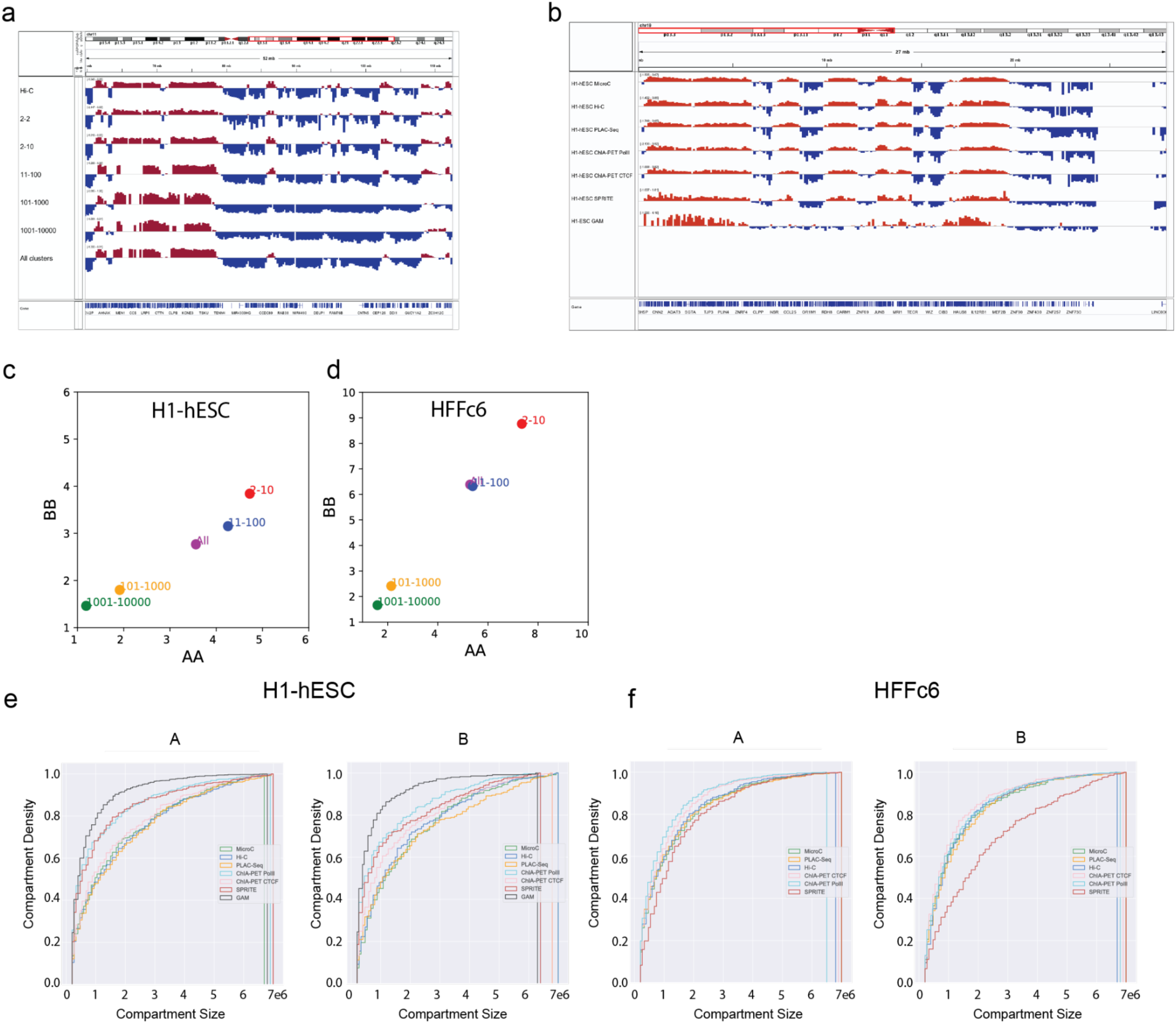
Chromatin interaction assays quantitatively differ in detection of small compartment domains. **a.** Eigenvector 1 obtained from SPRITE data derived from a range of cluster sizes along a typical genomic loci, showing that small compartments are not detected when data from larger SPRITE clusters is used. **b.** Examples of Eigenvector 1 profiles obtained from data generated with the different genomic assays indicated. **c.** Compartmentalization strength calculated with interaction data obtained with different SPRITE clusters, for H1-hESC and HFFc6. **e-f.** Cumulative distributions of compartment sizes, as detected with Hi-C, Micro-C, ChIA PET, PLAC Seq, and SPRITE (cluster size 2-100) for H1-hESC and HFFc6 cells.

#### Detection of chromatin loops

We next evaluated the ability of different chromatin interaction methods to detect chromatin loops, defined as focally enriched long-range interactions between specific pairs of loci (Figure 3a). For this analysis, we employed a hybrid approach, combining our previously developed platform-agnostic tool Peakachu ^61^ with platform-specific methods, which allowed us to effectively identify loops for different assays (Supplemental Methods). We currently do not have tools to detect significant looping interactions with sufficient resolution from GAM data, and the sequencing depth for SPRITE is not sufficient, and therefore did not attempt to call loops using those datasets. However, Aggregate Peak Analysis (APA) revealed that chromatin loops detected with other methods exhibited enriched SPRITE and GAM signals, albeit weaker compared to data obtained from other methods. Notably, smaller SPRITE clusters with 2-10 fragments displayed greater enrichment for such loop signals than larger clusters (Extended Data Figure 2).

Combining all chromatin loops detected, we defined a union set of loops and loop anchors for each cell type (H1-hESC: 91,960 anchors; HFFc6: 80,003 anchors). We first examined the extent to which loop anchors are detected with multiple assays. For H1-hESC cells, a set of 13,569 loop anchors were detected by all assays, representing the largest subset of anchors. The second largest subset comprised 10,542 loop anchors detected by Hi-C, Micro-C, and CTCF ChIA-PET, while the third largest subset was solely detected by Pol II ChIA-PET (Figure 3b). Conversely, in HFFc6 cells, a clearer separation between loop anchors detected through unbiased and targeted methods emerged: the largest subset consisted of loop anchors exclusively detected by Pol II ChIA-PET, the second largest subset being loop anchors solely detected by Micro-C, and the third loop anchors detected by both Hi-C and Micro-C (Extended Data Figure 3a).

We then systematically investigated chromatin features, such as histone modifications and CTCF binding, of loop anchors identified with different assays. For loop anchors detected by each combination of chromatin interaction mapping methods, we calculated fold-enrichment scores for various chromatin states (Figure 3b, and Extended Data Figure 3a). These states were defined by ChromHMM ^62^, a software that uses ChIP-Seq signals of various factors to learn and segment the genome into different states (ChromHMM states, i.e., strong enhancers, poised promoters, active promoters, transcriptional transition states, etc. - see Supplemental Figure 2 and Supplemental Methods). Through clustering analysis, we observed that loop anchors in both H1-hESC and HFFc6 cells could be categorized into two main groups: 1) anchors detected by Pol II ChIA-PET, characterized by high enrichment for active promoters, strong enhancers, and the transcriptional transition state; and 2) anchors detected by Hi-C, Micro-C, or CTCF ChIA-PET, exhibiting enrichment primarily for CTCF-bound sites.

Based on the composition of chromatin states at chromatin loop anchors, we further projected the union set of chromatin loops (H1-hESC: 124,061 loops; HFFc6: 115,850 loops) onto a two-dimensional (2D) space using Uniform Manifold Approximation and Projection (UMAP) (Supplemental Methods) and identified 6 loop clusters (Figure 3c for H1-hESC; Extended Data Figure 3b for HFFc6). Again, we observed distinct chromatin states for different loop clusters (Figure 3d, Extended Data Figure 3c and Supplemental Figure 3): 1) the first cluster predominantly featured loops between poised promoters; 2) the second cluster mainly consisted of loops between insulators; and 3) the remaining 4 clusters exhibited varying degrees of enrichment for transcription-related chromatin states, including active promoters, weak promoters, strong enhancers, transcriptional elongation, and transcriptional transition states.

Interestingly, we found that the enriched chromatin states for some clusters differed greatly between H1-hESC and HFFc6 cells, likely reflecting extensive epigenetic reprogramming and loop dynamics during the developmental process that gives rise to the two cell types. Moreover, insulator-related loops in the second cluster were largely longer-range, while transcription-related loops in clusters 3-6 were largely shorter-range, which agree with previous findings that short-range chromatin loops are more relevant to gene regulation (Figure 3d and Extended Data Figure 3c) ^63–65^.

**Figure 3.**
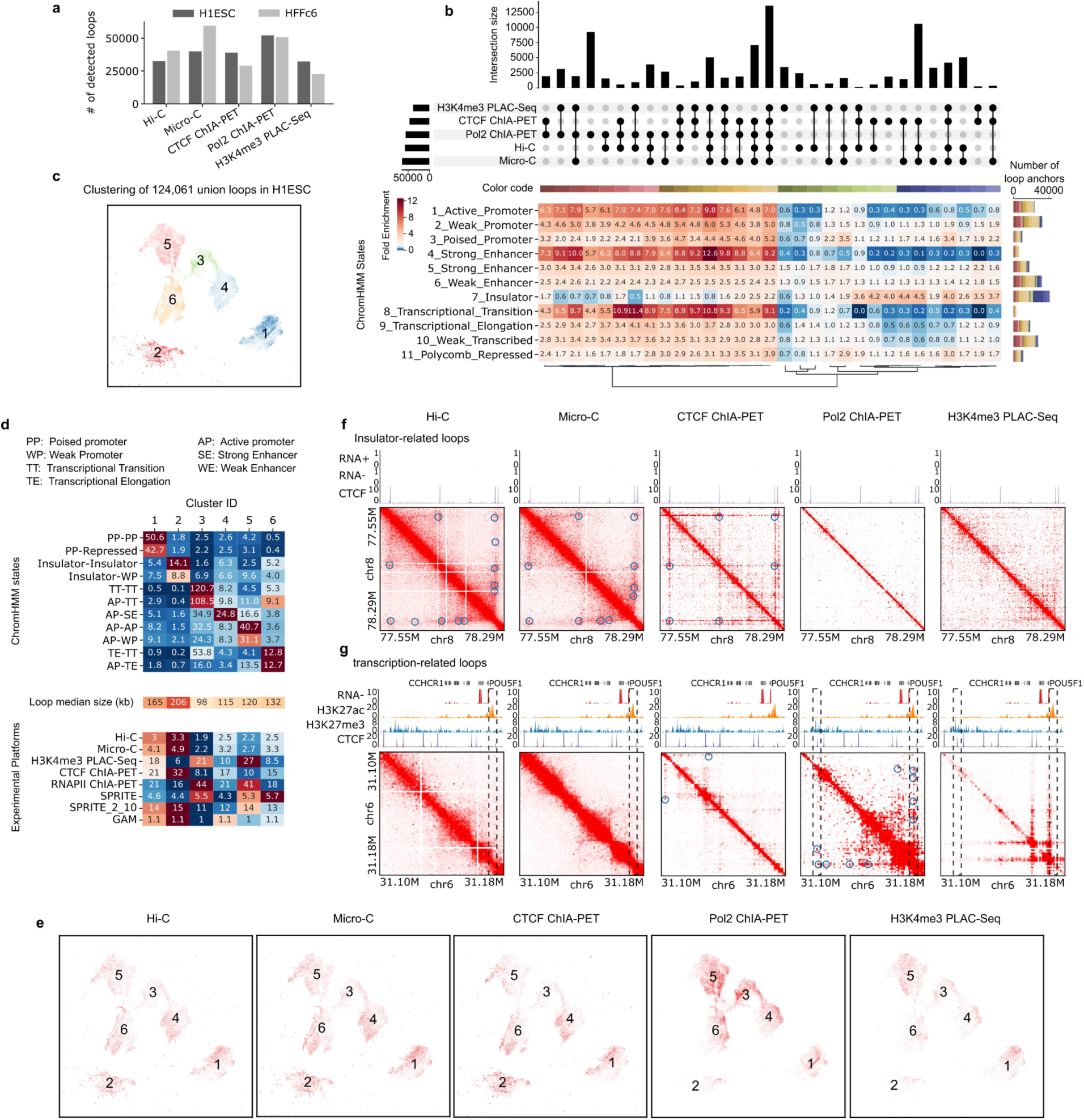
Cross-platform loop comparisons. We identified and compared chromatin loops from 5 experimental methods: Hi-C, Micro-C, CTCF ChIA-PET, RNA Pol II ChIA-PET, and H3K4me3 PLAC-Seq. **a.** The number of detected loops in each platform in two 4DN tier 1 cell lines H1-hESC and HFFc6. In panels b-g, we only included data from H1-ESC. **b.** (top) Upset plot comparing loop anchors from different platforms. (bottom) Fold enrichment scores of ChromHMM states for each loop anchor category in the upset plot. The bar plot on the right represents the number of loop anchors overlapping with different chromatin states, and different colors in a bar represent different categories in the upset plot. **c.** UMAP projection and clustering of the 124,061 union loops in H1-ESC based on the composition of ChromHMM states at interacting loop anchors. **d.** For different loop clusters in panel c, we calculated fold enrichment scores of ChromHMM states, the median genomic distances between the loop anchors, and the average loop strengths in different platforms. SPRITE_2_10 represents a subset of DNA SPRITE clusters with 2-10 fragments. **e.** UMAP projection of chromatin loops from individual platforms. **f.** An example showing the differences of platforms in detecting insulator-related loops. Contact maps are plotted at the 5 kb resolution, and chromatin loops are marked by blue circles. **g.** An example showing the differences of platforms in detecting transcription-related loops. Contact maps are plotted at the 1kb resolution, and chromatin loops are marked by blue circles.

**Extended Data Figure 2.**
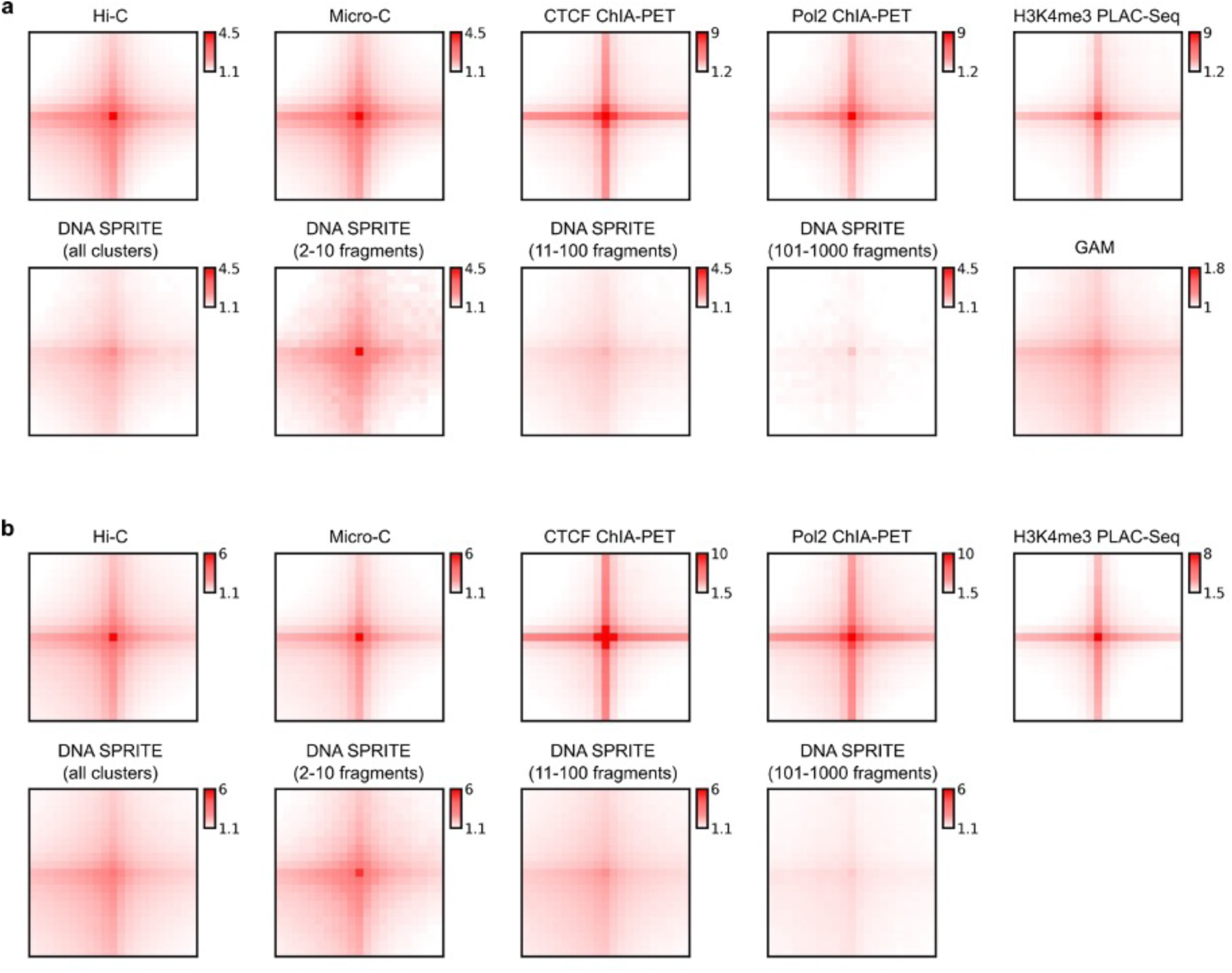
Aggregate Peak Analysis for the same high-confidence chromatin loops in different platforms in H1-hESC (a) and HFFc6 (b). The high-confidence loops for both cell lines were defined as those that can be identified by at least two methods. For each platform, distance-normalized signals at the 25 kb resolution within the 21×21pixel window centered at the coordinate of each loop are extracted and aggregated.

**Extended Data Figure 3.**
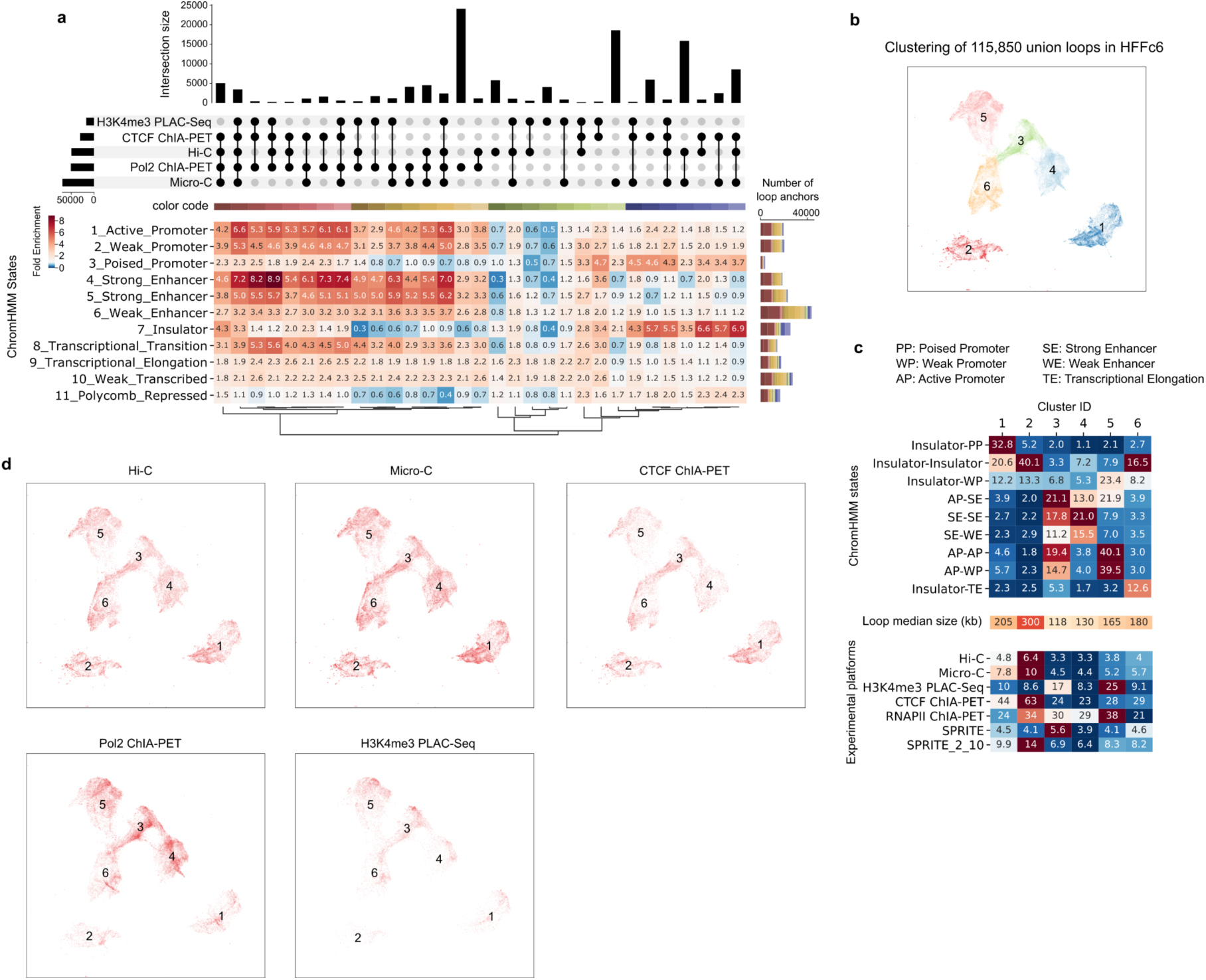
Cross-platform loop comparisons in the HFFc6 cell line. **a.** (top) Upset plot comparing loop anchors from different platforms. (bottom) Fold enrichment scores of ChromHMM states for each loop anchor category in the upset plot. **b.** UMAP projection and clustering of the 115,850 union loops in HFFc6 based on the composition of ChromHMM states at interacting loop anchors. **c.** For different loop clusters in panel b, we calculated fold enrichment scores of ChromHMM states, the median genomic distances between the loop anchors, and the average loop strengths in different platforms. SPRITE_2_10 represents a subset of DNA SPRITE clusters with 2-10 fragments. **d.** UMAP projection of chromatin loops from individual platforms.

We further characterized loops using additional transcription factor binding data and found that loops from different clusters are enriched for distinct transcription factors (Extended Data Figure 4; Supplemental Methods). Notably, Polycomb-group (PcG) proteins, such as EZH2 and RNF2, are specifically enriched at loop anchors in the first cluster. This is reminiscent of recent studies suggesting that a subset of chromatin loops is mediated by Polycomb repressive complex 2 (PRC2) and may play a role in chromatin compaction and gene repression (see ^66,67^ and references therein). Interestingly, the PcG proteins consistently co-localized with KDM4A. Originally known as a demethylase targeting H3K36me3 and H3K9me3, recent studies have shown that knockdown or overexpression of KDM4A can also significantly change the levels of H3K27me3 ^68,69^. This suggests KDM4A might play an important role in mediating a Polycomb- type repressive state for these specific chromatin loops by coordinating with the PcG proteins. As expected, insulator-related loops in the second cluster exhibit the highest enrichment of CTCF and cohesin binding at their anchors but display relative depletion of a wide range of transcription factors. Promoter-promoter loops in the third and fifth clusters are characterized by strong binding of RNA polymerase II, chromatin remodeling proteins, transcription initiation factors, and histone lysine demethylase, consistent with their roles in gene activation. Moreover, cohesin ChIP signal is observed across all loop anchors, suggesting that cohesin is localized at the bases of many or even most types of loops. Loops in the fourth and sixth clusters are of particular interest given that they are less enriched for RNA polymerase II, but are highly enriched for cohesin binding, which suggests that cohesin might be important in mediating enhancer-promoter loops (note that these clusters are enriched with interactions between enhancers and promoters based on Figure 3d).

We proceeded to calculate the average interaction strength for each loop cluster as detected by individual assays (Figure 3d and Extended Data Figure 3c; Supplemental Methods). As expected, in Hi-C, Micro-C, SPRITE, GAM, and CTCF ChIA-PET, the insulator-related loops exhibit relatively higher interaction strength compared to loops in the other 5 clusters.

Conversely, in RNA Pol II ChIA-PET and H3K4me3 PLAC-Seq data, transcription-related loops demonstrate elevated contact strength. Subsequently, we highlighted chromatin loops detected with each of the chromatin interaction assays on the two-dimensional UMAP projection of the union set of loops (Figure 3e and Extended Data Figure 3d). Consistently, we found pulldown-based methods with different enriched factors detect different subsets of chromatin loops: while CTCF ChIA-PET predominantly captures insulator-related loops, both RNA Pol II ChIA-PET and H3K4me3 PLAC-Seq are specifically enriched with transcription-related loops in clusters 3-6.

Interestingly, although both Hi-C and Micro-C are designed to detect all-to-all chromatin interactions without bias, loops detected with these methods are less enriched with transcription-related loops. These assays appear to excel in detecting structural CTCF-anchored loops (Figure 3f-g).

Together, these analyses reveal that different chromatin interaction assays tend to detect different types of chromatin loops with different efficiency. Unbiased genome-wide assays such as Hi-C and Micro-C most efficiently capture CTCF/cohesin based loops, while assays targeting RNA Pol II or H3K4me3 capture more transcription-related loops, which is by design. Further, most loop anchors, of any type, are associated with cohesin, suggesting a general role of this loop-extrusion complex in chromatin looping.

**Extended Data Figure 4.**
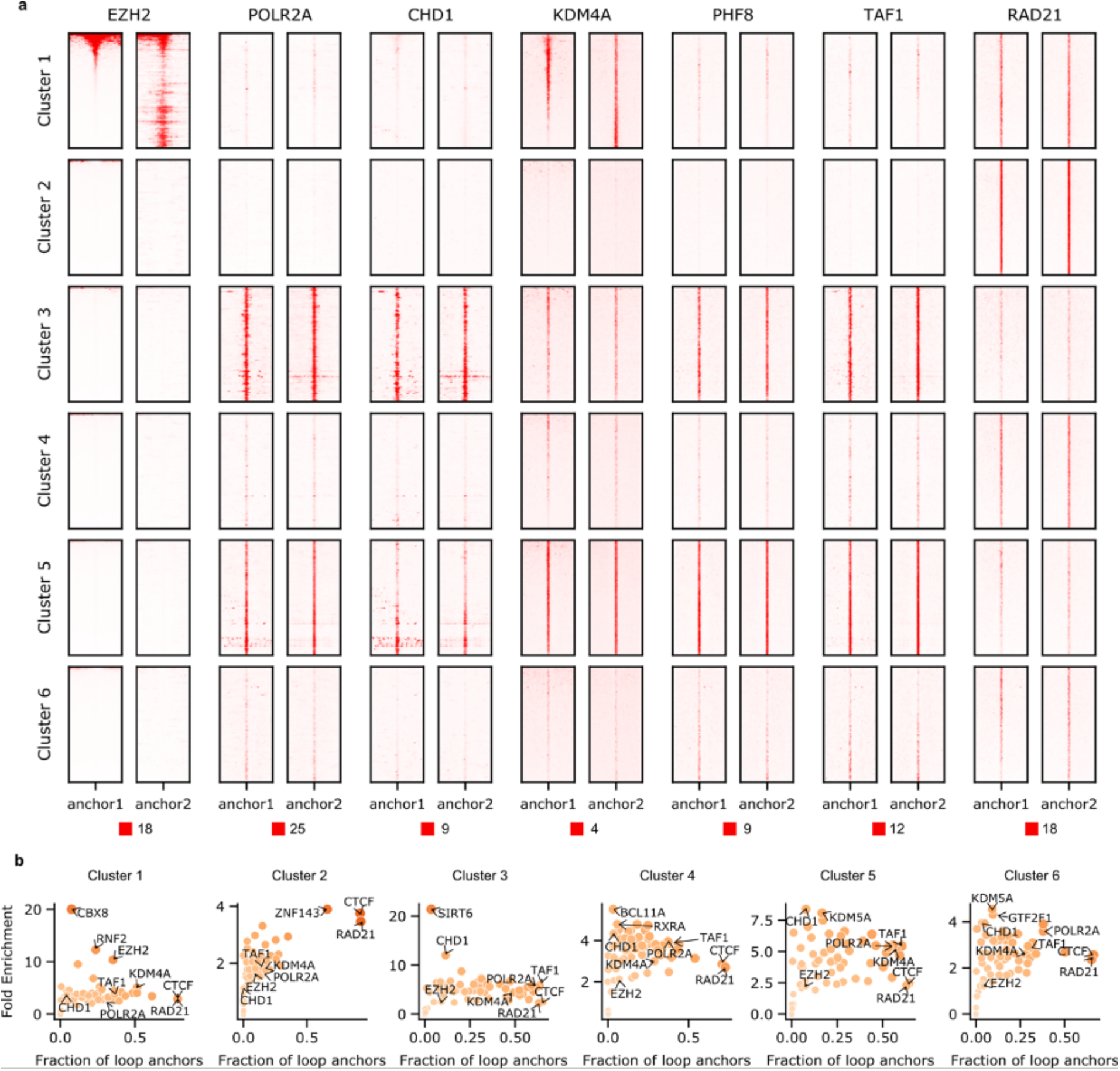
Transcription factor binding signatures of different loop clusters in H1-hESC. **a.** ChIP-Seq binding profiles of selected transcription factors surrounding both anchors from different loop clusters. Each row represents one loop. **b.** Fraction of loop anchors bound versus fold enrichment for 62 transcription factors.

### Annotation of spatial chromatin compartmentalization through integrative modeling

Previously, we have demonstrated that it is possible to derive linear genome-wide annotations of spatial nuclear compartments by integrating complementary 3D genome mapping data, such as subnuclear spatial localization data obtained with TSA-seq, DamID, and Hi-C data into a unified probabilistic model SPIN ^70^. The resulting annotations (i.e., SPIN states) reveal distinct patterns of spatial localization of loci relative to multiple types of nuclear bodies and show strong connections between large-scale chromosome structure and function, including replication timing and gene expression ^59^. Here, we further applied an improved SPIN framework with the support of joint modeling on multiple cell types to identify primary SPIN states in H1-hESC and HFFc6 cells relative to nuclear speckles, nucleolus, and nuclear lamina. We integrated datasets of TSA-seq and DamID to map proximity to nuclear bodies, and Hi-C to map chromatin interactions, (Figure 4, see Supplemental Methods) and identified nine SPIN states with distinct patterns of chromatin compartmentalization. These SPIN states are Speckle, Interior Active 1, 2, 3 (Interior_Act1, Interior_Act2, and Interior_Act3), Interior Repressive 1, 2 (Interior_Repr1 and Interior_Repr2), Near Lamin 1, 2 (Near_Lm1 and Near_Lm2), and Lamina state (Supplemental Figure 4a). The functional annotations of these nine SPIN states were verified by comparisons with various functional genomic data (see below) and exhibit a high correlation with the original SPIN states identified in K562 cells (Supplemental Figure 4b). Overall, different SPIN states have distinct distributions and combinations of TSA-seq and DamID signals, reflecting distinct patterns of spatial compartmentalization (Supplemental Figure 4c).

#### SPIN states stratify by histone modifications

To gain a better understanding of the transcriptional regulatory landscape of these SPIN states, we measured the enrichment of ChIP-seq signals for a range of histone marks on each SPIN state as compared to the genome-wide average for each mark. For H1-hESCs, we used ChIP-seq data, and for HFFc6 cells ChIP-seq data imputed using Avocado ^71,72^. We found that as the SPIN state changes from nuclear periphery to interior (e.g., Lamina state to Speckle state), the enrichment of active histone marks (e.g., H3K27ac, H3K4me1, H3K4me3, and H3K9ac) increases, along with gradual depletion of the repressive heterochromatin mark H3K9me3 (Figure 4), consistent with what we reported earlier ^70^ with additional cross cell-type comparisons. Notably, active histone marks such as H3K4me1, H3K4me2, H3K4me3, and H3K27ac are most prevalent in Speckle states (*P* < 2.2E-16), followed by Interior_Act1/2/3 states. Similarly, CTCF is most enriched in speckle-associated and interior active states, consistent with recent studies ^73^ and shows an overall decrease from interior to peripheral SPIN states (see the HFFc6 analysis in Figure 4). Although the correlations with histone marks are generally consistent between H1-hESC and HFFc6, certain histone marks exhibit more variable association with SPIN states. In particular, H3K27me3 is more enriched in Interior_Repr1 states in HFFc6 but has a stronger association with Interior_Act3 and Speckle in H1-hESC, indicating cell type-dependent and variable nature of spatial localization loci associated with specific histone marks (Figure 4, left panel). The variable distribution of H3K27me3 is also observed in CUT&RUN data (Figure 4, bottom right panel). Previous work also reported a high cell type-specific distribution of H3K27me3 across human cell lines ^74^. In addition, we found that more interior SPIN states are more ubiquitous across cell types than more peripheral SPIN states, consistent with previous observations of conservation of nuclear speckle associated domains ^56^. These results reveal that SPIN states have a generic correlation with active histone marks.

However, at least in certain cell types, SPIN states have a cell type-specific distribution of repressive histone marks such as H3K27me3.

The different SPIN states also show clear separation of the multi-fractions for high resolution Repli-seq ^75^ data in H1-hESC, manifesting the strong connection between chromatin spatial localization and replication timing as a crucial genome function (Figure 4, middle, and see below).

#### SPIN states differentially associate with chromatin-associated RNAs

We further compared SPIN states with different types of chromatin-associated RNAs (caRNAs) detected with iMARGI ^76–78^. RNA facilitates spatial compartmentalization in the nucleus ^79^. caRNAs can promote or suppress chromatin looping depending on their associated genomic sequences. Loop-anchor associated caRNAs often promote looping, including enhancer-promoter loops ^80^, whereas between-loop-anchor-associated caRNA often suppress looping ^78^. We asked whether any SPIN state is enriched with chromatin-associated RNAs (caRNAs) containing specific types of sequence features, especially repetitive elements. We selected the caRNAs if their RNA ends in iMARGI mapped to repetitive elements and further stratified them into different groups according to SPIN states on their DNA ends (Supplemental Methods). To avoid bias due to nascent RNA transcripts interacting with genomic regions where they are transcribed, we only included interchromosomal iMARGI pairs, where the transcription and interacting genomic regions are on different chromosomes. We found that the genomic target sequences of different types of repeat sequences associated with different types of caRNAs are enriched for distinct SPIN states (Figure 4, right panel). The caRNAs that contained Alu, srpRNA, SVA, and snRNA repeat elements are mostly enriched in interior SPIN states (e.g., Speckle, Interior_Act1/2/3), while the caRNAs contained connected to L1, ERVL, ERV1, and LTR repeats are enriched in SPIN states closer to nuclear lamina (Lamina, Near_Lm2) (as shown in Figure 4). Thus, the caRNA’s sequence features correlate with the 3D compartmentalization of the caRNA’s target genomic regions. Similar to the observations of histone marks, SPIN states away from the interior also show more variable and complex patterns with associated caRNAs. This suggests that genomic sequence features are associated with the 3D chromatin localization of the RNA’s target regions.

Together, these results demonstrate that the SPIN framework effectively integrates various nuclear organization mapping data to produce genome-wide large-scale compartmentalization patterns relative to multiple nuclear bodies. These SPIN states stratify orthogonal functional genomic data, including histone modification, replication timing, and RNA association.

**Figure 4.**
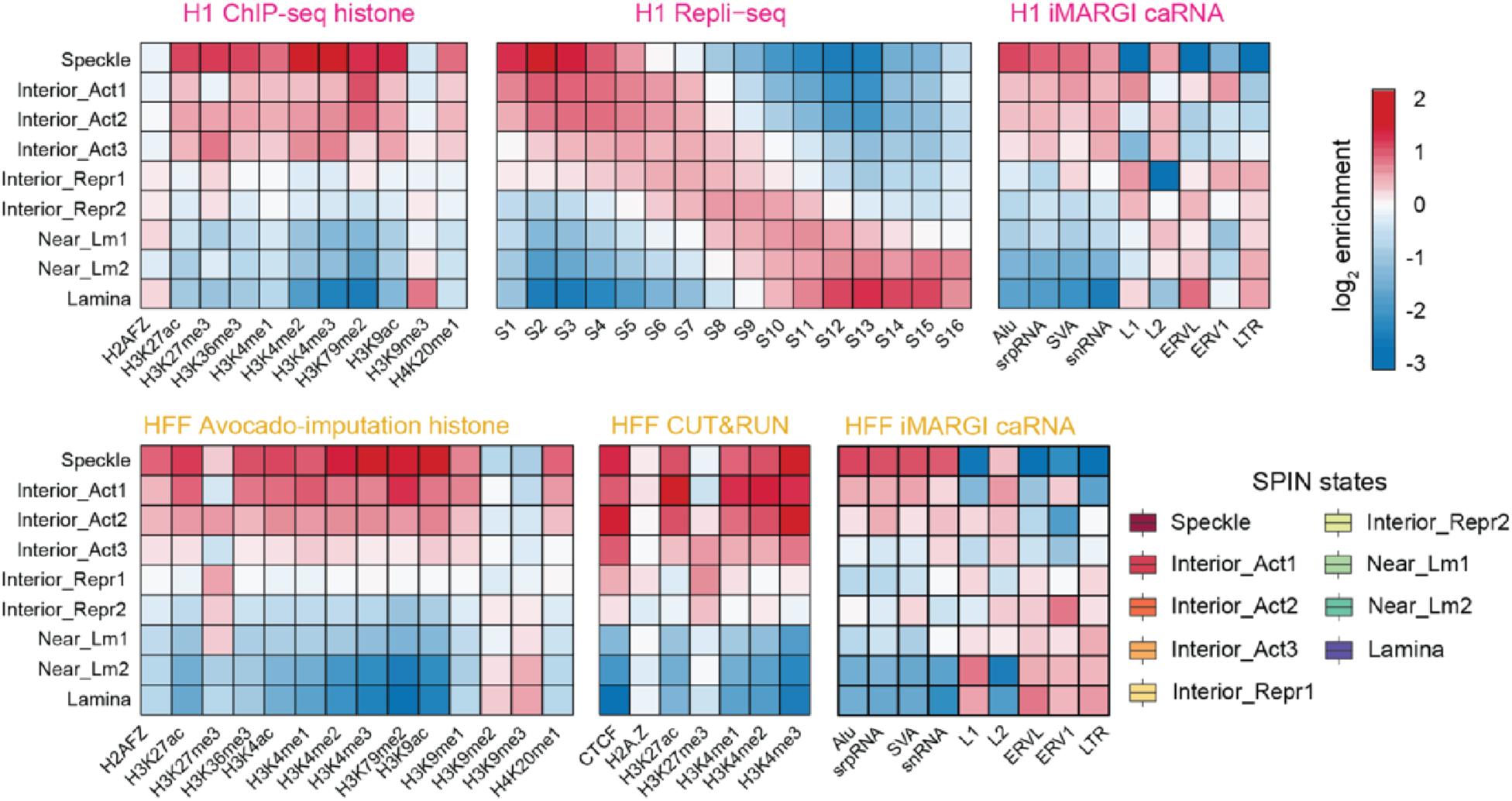
SPIN states stratify the genome into distinct spatial compartments. Heatmaps show the enrichment of histone marks, Repli-seq signals, and caRNAs (columns) on different SPIN states (rows). Colors of the heatmap indicate the log2 fold-change enrichment calculated as the ratio of observed signals over genome-wide expectation.

### A three-dimensional view of the genome

Using ensemble Hi-C, Lamin B1 DamID, and SPRITE data, we employed our integrative genome modeling platform (IGM) ^81^ to construct a population of 1,000 single cell 3D genome structures at 200 kb resolution for H1-hESC and HFFc6 cells. These structures illuminate the folding of chromosomes within the nuclear topography in single cells (Figure 5a,b), as assessed by data from multiplexed FISH imaging ^82^ and TSA-seq data (Supplementary Figure 5). To characterize the nuclear microenvironment of an allele in a cell, we define 14 structural features that collectively specify its 3D position within the nuclear topography relative to the predicted locations of nuclear bodies, compartments and properties of the chromatin fiber, as described in ^83^ (Figure 5c).

#### SPIN states

First, we assess the nuclear microenvironment of chromatin in different SPIN states (Figure 5a,b,c), which exhibit distinct enrichment patterns of 3D structure features (Figure 5c, Extended Data Figure 5a-d). For instance, the radial nuclear positions gradually increase from ‘Speckle’ (in the center) to ‘Lamina’ SPIN states (at the periphery) in HFFc6 cells, confirming expectations from the SPIN state analysis (Figure 5cd, Extended Data Figure 5a,c). Our analysis also reveals associations of specific SPIN states to nuclear bodies, such as nuclear speckles (Extended Data Figure 5a,c) or enriched interactions of ‘Interior Act 2’ and ‘Near Lamina 1’ SPIN state chromatin with the nucleolus (Figure 5c). Moreover, SPIN states show characteristic projections of their 3D structure features confirming their structural distinction (Extended Data Figure 5b).

#### Genome structure differences between cell types

Differences in gene expression between H1- hESC and HFFc6 cells frequently correlated with notable differences in the nuclear microenvironment of genes, illustrated by different distances from nuclear speckles (Figure 5e) and the radial positioning of chromosome 1’s p-arm (Figure 5f). For instance, the transcription factor POU3F1 (OCT-6) plays a pivotal role in cell differentiation and maintenance of the nervous system ^84,85^. Expressed in H1-hESC, the POU3F1 gene shows relatively small speckle distances in a high fraction of cells (high speckle association frequency, or “SAF” in Figure 5g, Extended Data Figure 5g) and is predominantly located in the nuclear interior (RAD (average radial position) in Figure 5f,h), often at chromosome boundaries (high ICP (inter-chromosomal contact probability) and transA/B in Figure 5g, Extended Data Figure 5g), confirming the ‘Speckle’ SPIN state observed in H1-hESC. In HFFc6, POU3F1 is silent and shows a different nuclear microenvironment, located towards the nuclear periphery in a high fraction of cells (RAD in Figure 5eg, Extended Data Figure 5g), with predominantly local intra-chromosomal interactions (low ICP) and a high degree of chromatin fiber compaction (low RG (radius of gyration over 1Mb window)) (Figure 5g,h, Extended Data Figure 5g). Consistently, POU3F1 is in the ‘Speckle’ SPIN state in H1-hESC while it is in the ‘Interior Rep1’ SPIN state in HFFc6 cells. Our observations revealed opposing trends in nuclear locations for genes that are silent in H1- hESC but active in HFFc6. THBS1 (Thrombospondin 1), a gene encoding a protein that promotes cell adhesion in connective tissue cells, exemplifies this observation ^86,87^ (Extended Data Figure 5h left panel). Consistently the SPIN state of THBS1 differs between H1-hESC and HFFc6 cells, with a ‘Near Lamina2’ state in H1-hESC and a ‘Speckle’ state in HFFc6.

Interestingly, genes silent in H1-hESC often show a bimodal distribution of their nuclear locations characterized by smaller subpopulation of alleles presented in a nuclear microenvironment with features of an active state (i.e., THBS1, CAV1 dotted line in Extended Data Figure 5h), which could either indicate increased structural heterogeneity between individual cells or the presence of a subpopulation of H1-hESCs in a different state. Genes highly expressed in both cell types typically show similar microenvironments, exemplified by overlapping 2D distributions of their joint speckle and lamina distances in the models of both cell types (Extended Data Figure 5i).

#### Gene expression

To further study the links between nuclear topography and gene expression, we analyzed the nuclear microenvironment of the 25% most highly and lowly expressed genes in our HFFc6 genome structure models. Overall, most of these genes separate in projections of their 3D structure features according to differences in expression level, highlighting a distinct correlation of the nuclear microenvironment with gene expression (Figure 5i) (Extended Data Figure 5e,f). 90% of the most highly expressed genes (including 73% of all housekeeping genes) have a high to medium speckle association frequency, confirming previous similar observations ^33,54,88^. Among those, the 2,275 genes with the highest speckle associations (top 25% SAF) have predominant interior radial locations (RAD), relatively low cell-to-cell variability in their microenvironment (low **δ**RAD, **δ**SpD) and a high level of inter-chromosomal exposure (high ICP) (37% of these genes are housekeeping genes) (Figure 5j, right panel, Figure 5k, Supplemental Figure 6a,b). We categorized these genes to be part of the ‘class I’ microenvironment (Figure 5j, Supplemental Figure 6ab) (Methods). These genes exhibit relatively short lengths and are located in regions of high gene density (Supplemental Figure 6ab). However, 10% (659 genes) of the most highly expressed genes belong to a very different nuclear microenvironment, designated as ‘class II’, typically associated with lowly expressed or silenced genes (Figure 5j, right panel). These genes feature minimal speckle associations (25% lowest SAF) and high cell-to-cell variability in speckle distances and radial positions (high SpD, **δ**SpD, **δ**RAD, low SAF) (Figure 5k,l, Supplemental Figure 6ab). Situated more peripherally, these genes are frequently located within chromosome territories characterized by low inter-chromosomal proximities (low ICP) and exhibit enhanced associations with B compartment chromatin (low transAB) (Figure 5k, Supplemental Figure 6ab). They typically are long genes and reside in regions of low gene density (Figure 5l). Compared to lower transcribed class II genes, they exhibit notably higher CpG density and H3K4me3 and H3K27ac levels at their promoter sites (p-value < 10^-5^ for both H3K4me3 and H3K27ac) (Supplemental Figure 6ab). 19% of these genes are identified as housekeeping genes, highlighting that housekeeping genes can be found in at least two contrasting nuclear microenvironments. Overall class II gene promoters generally exhibit fewer enhancers within close sequence distances (<100 kb) compared to class I genes (E in Figure 5l). However, highly expressed class II genes demonstrate higher enhancer densities when normalized by the number of nearby genes (<100 kb) than any other genes (E/G in Figure 5l, Supplemental Figure 6ab). A distinction between class I and II gene promoters is evident when we quantify the number of accessible enhancers within a spatial distance of 350 nm in the 3D folded genome structures (E^N^/G, i.e. spatial enhancer density) ^89,90^. Highly expressed class II genes exhibit a greater spatial enhancer density from long-range intra-chromosomal proximities compared to class I genes (p-value < 10^-5^) (E^N^_Intra/G and E^N^_Inter/G in Figure 5l, Supplemental Figure 6ab), which in turn display notably higher contributions from inter-chromosomal proximities. This distinction likely occurs because class I genes more frequently protrude beyond their chromosome territory towards speckles, fostering heightened inter-chromosomal proximities at speckle sites (high ICP) (Extended Data Figure 5j). Highly expressed class II genes therefore have a larger number of enhancers accessible from long-range interactions (up to 2 Mb), while having a lower number of enhancers accessible at relatively close sequence distances (<100 kb) (Figure 5l,m). This observation could explain the lower number of detected high frequency enhancer loops for highly expressed class II housekeeping genes (see section below), as very long-range enhancer interactions are likely to be more variable between individual cells. Overall housekeeping genes in class I and II do not show substantial differences in their structural properties from other highly expressed genes within the same class (Supplemental Figure 6c,d).

In summary, our data-driven genome structure modeling revealed relationships between the nuclear environment and gene expression.

**Figure 5.**
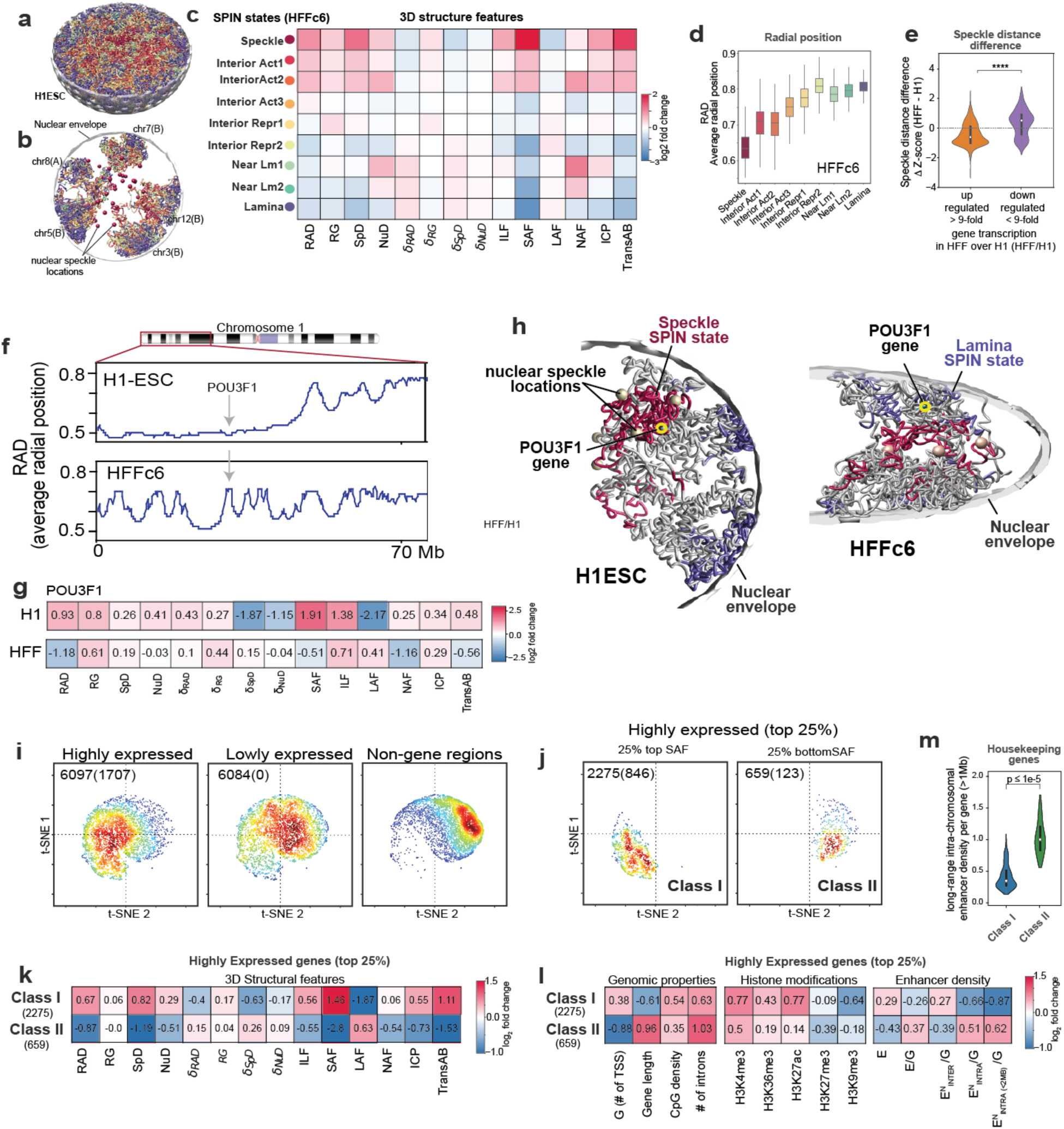
3D Structural features and their cell-to-cell variabilities in relation to gene function. **a.** Single cell genome structure model of the H1-hESC with genomic regions color coded by their SPIN states. **b.** Slice through genome structure in A with only a few chromosomes shown together with predicted speckle locations by red spheres, **c.** Enrichment of different structural features for chromatin in different SPIN states calculated from the population of models. RAD: 1-norm. average radial position, RG: chromatin fiber de-compactness (radius of gyration of chromatin fiber over +/-500kb), SpD: average speckle distance, NuD: average distance to nucleolus, ILF: interior localization probability (fraction of alleles within 50% percentile interior volume), SAF: speckle association frequency, LAF: lamina association frequency, ICP: inter chromosomal interaction probability, TransAB: trans A/B ratio, δ features (RAD, RG, Spd, Nud) show cell-to-cell variability of the respective feature (Methods). **d.** Box plots for the distributions of average radial positions of chromatin regions in each SPIN state **e.** Violin plots for distributions of speckle distance z-score differences for genes in H1-hESC and HFFc6 that are significantly up-regulated (up to 9-fold) and significantly down-regulated (by more than 9-fold) in HFF over H1. Shown are Z-score differences in speckle distances of genomic regions between both cell types (HFF - H1). **f.** Average radial positions for a part of chromosome 1 in H1-hESC cells (upper panel) and HFFc6 cells (lower panel). **g.** Log-fold enrichment of 14 structural features calculated for the POU3F1 gene in H1-hESC (upper panel) and HFFc6 cells (lower panel), calculated from the 3D structure population. **h.** (Left panel) Chromosome 1 in single cell 3D genome structure of H1-hESC (left panel) and HFFc6 (right panel). The nuclear location of POU3F1 gene is shown by a yellow circle, red color shows chromosomal regions annotated in the SPIN speckle state, blue regions show chromatin in the lamin SPIN state. Locations of predicted nuclear speckle locations closest to POU3F1 are shown. **i.** t-SNE projections of 3D structure feature vectors for chromatin regions containing the transcription start sites of genes with the highest and lowest expression quartiles in HFFc6, as well as chromatin regions without known genes. Shown are also the number of all genes in each group, while the number of housekeeping genes within each group are shown in parenthesis. **j.** t-SNE projections of the 25% most highly expressed genes with the top (left panel) and lowest SAF quartile among all genes (bottom panel). **k.** log-fold enrichment of 14 3D structure features for highly expressed genes (top quartile), lowly expressed genes (bottom quartile) genes in class I and class II microenvironments. **l.** log-fold enrichment for genomic properties (within a 200kb region), histone modifications (within +/- 10kb of TSS), and 3D spatial enhancer densities at each TSS (Methods) for highly expressed genes (top quartile), lowly expressed (bottom quartile) genes in class I and class II microenvironments. **m.** Comparison of the intrachromosomal 3D spatial enhancer density at TSS of housekeeping genes in class I (high SAF) and class II (low SAF) microenvironments. Only enhancers at a sequence distance >1Mb are considered.

**Extended Data Figure 5.**
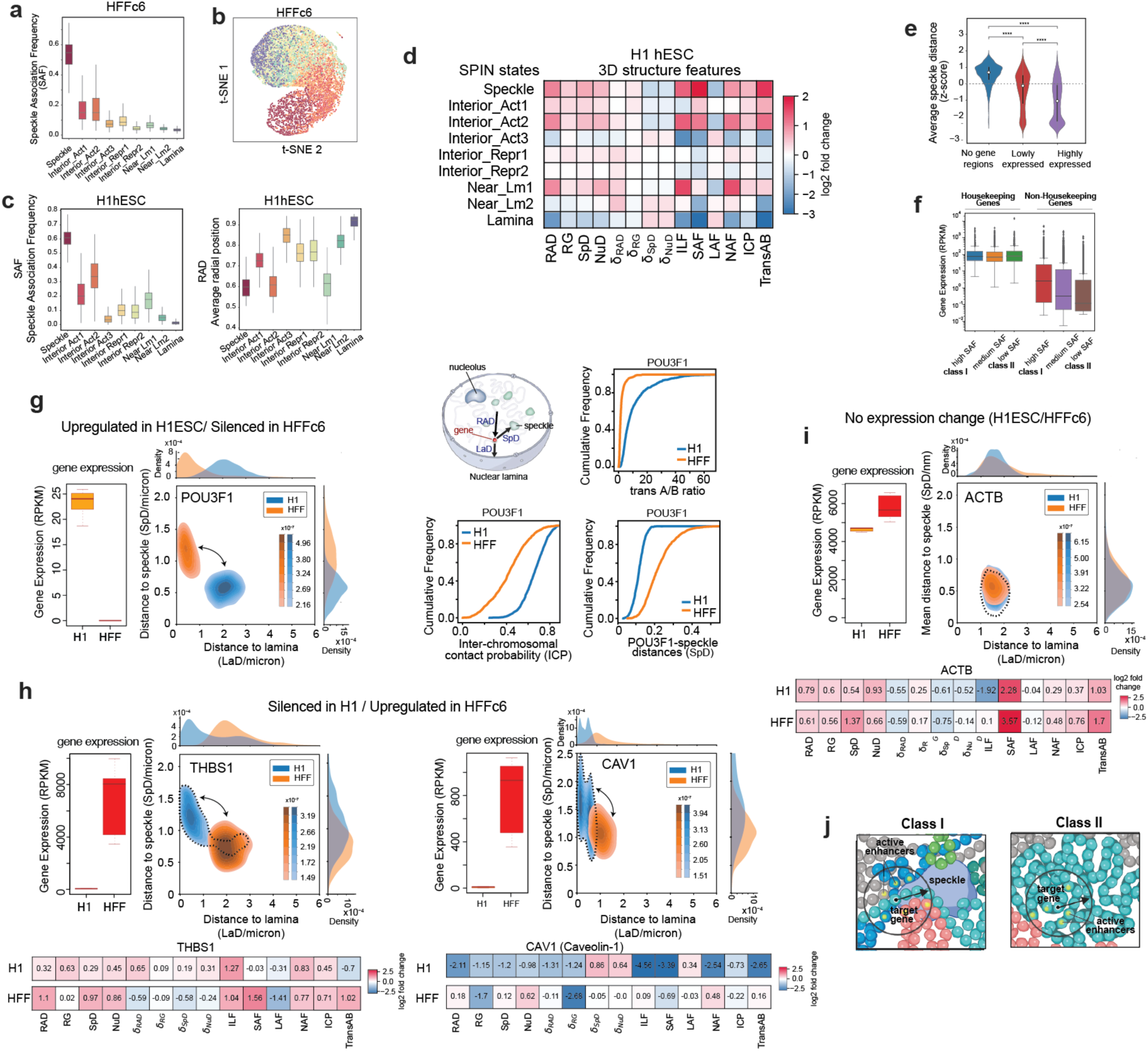
3D Structural features and their cell-to-cell variabilities in relation to gene function. **a.** Distributions of speckle association frequencies (SAF) for chromatin in different SPIN states calculated from a population of 3D structures of HFFc6 genomes. **b.** tSNE projections of structure feature vectors for chromatin in different SPIN states in HFFc6. **c.** Distributions of SAF and normalized radial positions for chromatin in different SPIN states calculated from a population of 3D structures of H1-hESC genomes. **d.** Log-fold enrichment of 14 3D structural features for chromatin in different SPIN states calculated from the population of models. **e.** Distribution of speckle association frequencies for chromatin in different gene expression categories in HFFc6. **f.** Distribution of gene expression levels for housekeeping and non-housekeeping genes stratified by their speckle association frequencies. **g.** (left top panel) Distribution of gene expression levels taken from 6 gene expression experiments for POUF31 gene in H1-hESC and HFFc6. (second panel from right). 2D distribution of joint speckle distance and lamina distance for POU3F1 genes in 1000 single cell models. (right top panel) Schematic view of structural features in single cell models. RAD: radial position, LaD, Distance to the nuclear envelope, SpD: Distance to the closest speckle. (remaining panels) Cumulative distributions from single cell structure features of POU3F1 gene in the simulated cell population. (speckle distance, inter-chromosomal contact probability (ICP), transAB ratio). **h.** (left panel) Gene expression levels of THBS1 in H1-hESC and HFFc6 cells. (middle top panel) distribution of joint speckle distance and lamina distance for THBS1 genes in 1000 single cell models. (lower panels) log-fold enrichment of 14 structure features for THBS1 gene in H1ESC and HFFc6 cells. (Most Right panel), the same panels are shown also for the gene CAV1. **i.** (left panel) Gene expression levels of ACTB in H1ESC and HFFc6 cells. (Middle top panel) distribution of joint speckle distance and lamina distance for ACTB genes in 1,000 single cell models. (lower panels) Structure feature enrichment for ACTB gene in H1-hESC and HFFc6 cells. **j.** Scheme illustrating decreased contributions from long-range sequence distances for class I and increased contributions for class II genes.

### Integrative single-cell 3D genome analysis unveils cell-to-cell variability of 3D genome structures

To further investigate the mechanisms orchestrating chromosome 3D architecture, we sought to further investigate the variability in single-cell 3D genome structure using different integrative modeling approaches. Previous comparisons between model-predicted single-molecule 3D structures and multiplex microscopy data showed that both loop-extrusion and polymer phase separation (Strings & Binders Switch model ^91–93^ recapitulate not only the average contact probabilities and patterns, but also the entire ensemble of microscopically observed single-molecule conformations in single cells, supporting the view that those are fundamental mechanisms shaping chromatin folding ^92,94,95^. Imputation of single-cell Hi-C contact maps using Higashi has also enhanced the single-cell 3D genome folding analysis with graph representation learning ^96,97^, which complements polymer modeling.

We first verified that both SBS polymer models and Higashi imputed scHi-C contact maps of the *DPPA* locus (chr3: 108.3 - 110.3Mb) in WTC-11 pluripotent stem cells at 10 kb resolution, which result in ensembles of structures that are consistent with bulk Hi-C at the population level (Figure 6a) and improve correlations between merged scHi-C and bulk Hi-C data (Figure 6a, also see Supplemental Methods).

Next, we calculated single-cell TAD-like domain boundaries using single-cell insulation scores ^96,97^ and observed that both SBS polymer models and Higashi imputed scHi-C data reveal the variability of TAD-like domain boundaries while remaining consistent with the insulation scores calculated from bulk Hi-C (Figure 6a). To view the direct correspondence of Higashi imputed scHi-C contact maps and SBS polymer models, we found nine mutual nearest neighbors by quantifying the pairwise similarities between scHi-C and polymer models (Figure 6b). This analysis supports cell-to-cell variability of TAD-like structures using an integrative approach from different analysis methods, consistent with observations based on multiplexed imaging methods98.

We next aimed to investigate the variability of chromatin loops at the single-cell level, which has not been analyzed extensively due to the sparsity of scHi-C data and the limitation of spatial resolution in multiplexed imaging methods. We utilized polymer models ^92,94,95^, Higashi-imputed scHi-C contact maps ^96,97^ and the recent SnapHiC contact maps ^99^ derived from WTC-11 scHi-C datasets, to identify chromatin loops at high sensitivity (Supplemental Figure 7). This integrative analysis of chromatin loops, A/B compartments, and TAD-like domains identified from single cells with different analysis approaches revealed that genomic loci within the same compartment or TAD-like domain are more likely to form stronger chromatin loops in the same cell (Figure 6c). For example, in a representative chromatin loop near gene *RABGAP1L*, the normalized loop intensity is much higher for single cells where this loop is located within the same TAD-like domain or the two loop anchors located in the same compartment, specifically in both A compartments (Figure 6d).

Together, our findings illustrate the cell-to-cell variation in chromatin folding in individual cells. Observations from integrative single-cell modeling and analysis suggest that the formation of loops, TAD-like domains, and compartments, although at different scales, are not merely correlated properties of chromatin folding observed in the bulk (averaged) contact maps.

Instead, these structures can be coherently observed at the single-cell and single-molecule level, indicating that they are likely driven by a hierarchical mechanism. The variability of chromatin folding in individual cells, at the level of loops, TADs and compartments, likely contributes to the dynamic regulation of gene expression and other nuclear processes, providing further insight into the complex nature and dynamics of genome organization and function.

**Figure 6.**
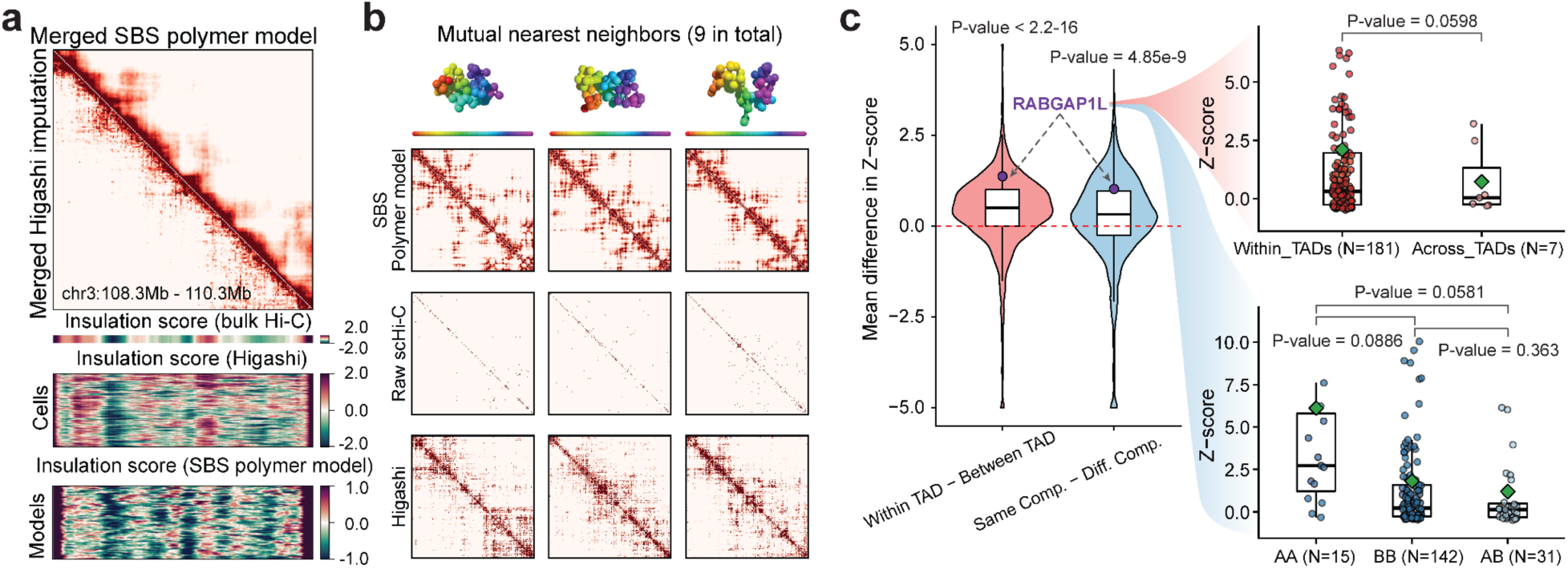
Cell-to-cell variabilities of 3D genome features. **a.** Heatmap on the top shows the merged scHi-C contact maps at a 2Mb region from chr3 imputed by Higashi (bottom-left) or predicted by the SBS polymer model (top-right). Insulation scores from bulk Hi-C, calculated insulation scores after Higashi imputation and SBS polymer modeling are shown at the bottom. **b.** 3D genome structure models, raw scHi-C contact cap, and the imputed contact map from three mutually similar cells between Higashi imputation and SBS model are shown. **c.** The average normalized intensity of chromatin loop across 188 WTC-11 cells is calculated and compared by dividing loops based on their relative position within TADs and A/B compartments. The pink boxplot (left) represents the difference between loops in the same TAD and loops spanning multiple TADs. The blue boxplot (right) shows the difference between loops in the same A/B compartment and loops spanning different compartments. A representative chromatin loop near gene RABGAP1L is highlighted in the box plot on the right. The original distribution of the normalized intensity of this specific loop in each cell is shown in the box plots on the right. Loops are stratified into different groups depending on whether this loop locates within one TAD or spans TADs (top) or the A/B compartments state of two loop anchors in each single cell.

### Connections between the 3D genome and genome function

#### Number of interacting enhancers is correlated with gene transcription

By using the union set of chromatin loops defined above, we examined the relationship between the number of distal enhancers linked to promoters and the transcription levels of corresponding protein-coding genes. Of 19,618 protein-coding genes, 14,730 in H1-hESC and 13,501 in HFFc6 display interactions with at least one distal enhancer. The median distance between interacting enhancers and promoters is 138 kb, notably shorter than that observed for insulator-mediated loops detected using the same datasets (Figure 3d, Extended Data Figure 3c, Extended Data Figure 6a). Importantly, genes with an increasing number of interacting enhancers tend to exhibit higher transcription levels (Figure 7a; Extended Data Figure 6b), and differences in the number of interacting enhancers are closely associated with gene transcription differences between cell lines (Extended Data Figure 6c).

**Figure 7.**
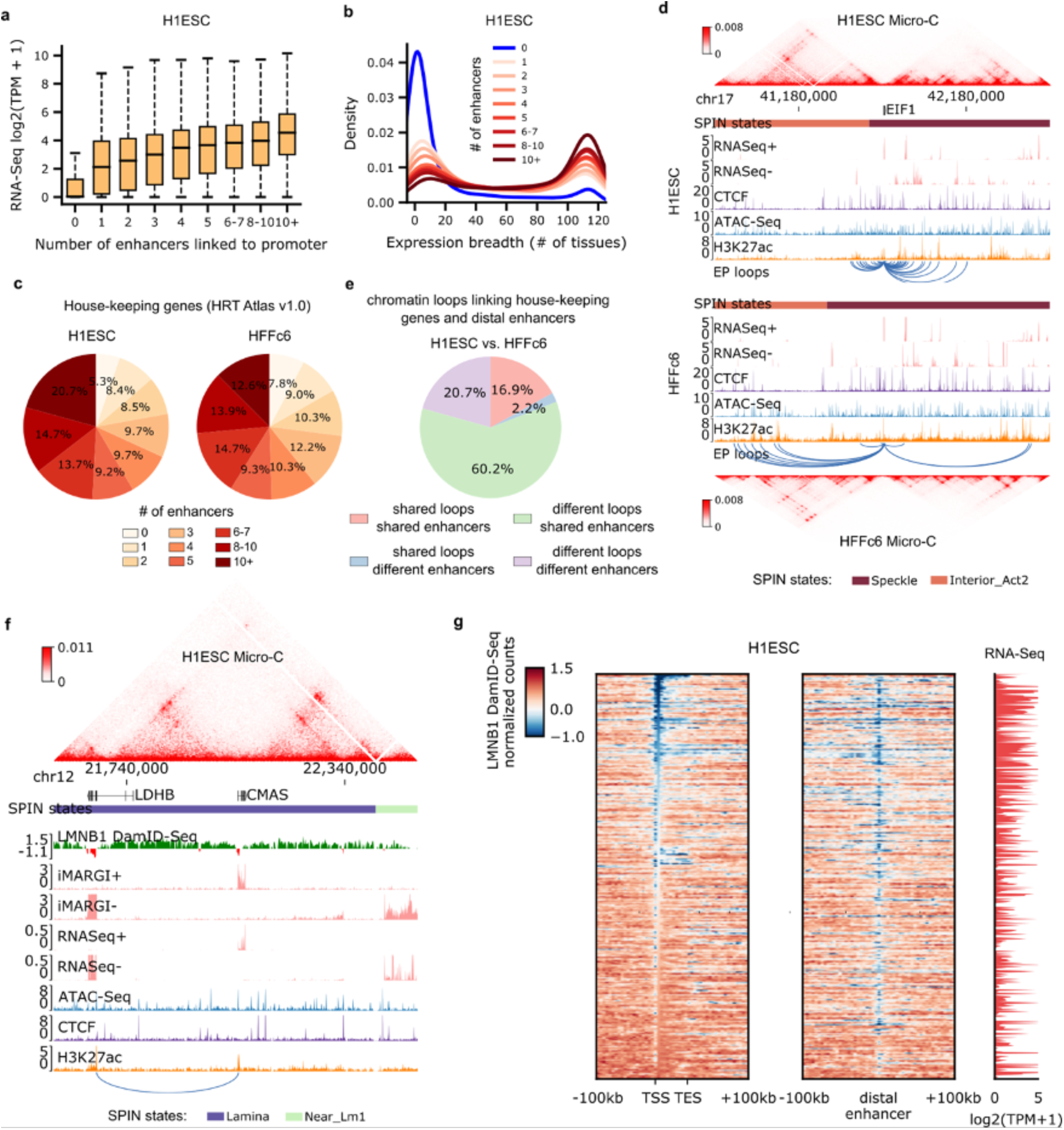
Associations of enhancer-promoter loops with gene regulation. **a.** Gene transcription levels versus the number of interacting enhancers in H1-hESC. For each boxplot, the center line indicates the median, the box limits represent the upper and lower quartiles, and the box whiskers extend to 1.5 times the interquartile range above and below the upper and lower quartiles, respectively. TPM, transcripts per kilobase million. **b.** Expression breadth (number of tissues a gene is expressed in) of genes with different number of interacting enhancers in H1-hESC. **c.** Percentages of house-keeping genes with different number of interacting enhancers. **d.** Genome browser view of a region surrounding the house-keeping gene EIF1. The blue arcs represent chromatin loops linking the EIF1 gene promoter with distal enhancers. **e.** The dynamics of chromatin loops linking house-keeping gene promoters and distal enhancers between H1-hESC and HFFc6. **f.** Genome browser view of the CMAS loci in H1-hESC. **g.** Lamin-B1 DamID-seq signals surrounding lamina-associated genes and their interacting enhancers in H1-hESC. Only genes with interacting enhancers in the Lamina SPIN state are included in this plot. TSS, transcription start sites. TES, transcription end sites. TPM, transcripts per kilobase million.

**Extended Data Figure 6.**
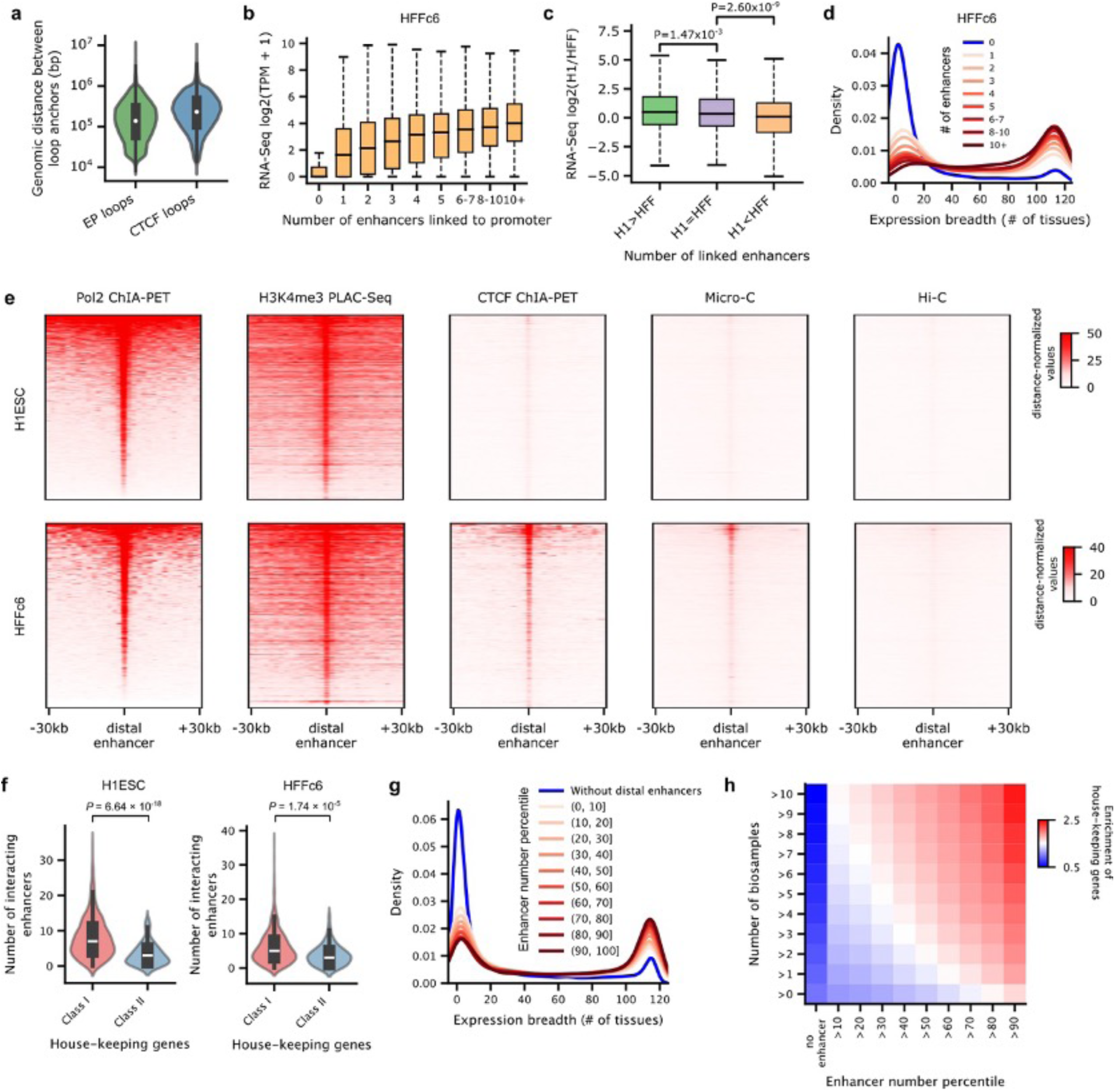
House-keeping genes are engaged in extensive enhancer-promoter loops. For each boxplot, the center line indicates the median, the box limits represent the upper and lower quartiles, and the box whiskers extend to 1.5 times the interquartile range above and below the upper and lower quartiles, respectively. **a**. Size distributions of enhancer-promoter (EP) loops and CTCF-mediated loops. Data were merged from the H1-hESC and HFFc6 cells. **b.** Gene transcription levels versus the number of interacting enhancers in HFFc6. TPM, transcripts per kilobase million. **c.** Fold-change of gene transcription levels (TPM) is depicted for three gene groups: genes with a higher number of interacting enhancers in H1-hESC, genes with the same number of interacting enhancers in both cell lines, and genes with a higher number of interacting enhancers in HFFc6. The P values were computed using the two-sided Mann–Whitney U-test. **d.** Expression breadth of genes with different numbers of interacting enhancers in HFFc6. **e**, Distance-normalized contact signals between house-keeping gene promoters and regions (+/-30 kb) surrounding the interacting enhancers. **f.** Comparison of the number of interacting enhancers between the two house-keeping gene classes defined in the 3D modeling section. The P values were computed using the two-sided Mann–Whitney U-test. **g.** Expression breadth of genes with different number of interacting enhancers. Data were merged from 32 cell lines or primary cells. In each sample, genes are categorized into 11 groups based on the percentile of the number of interacting enhancers. **h.** Enrichment of house-keeping genes across gene sets characterized by the number of interacting enhancers and supported samples. Each bin represents a specific combination of these factors. For instance, the top-right corner bin represents the enrichment score for genes with the number of interacting enhancers greater than the 90th percentile across over 10 samples.

**Extended Data Figure 7.**
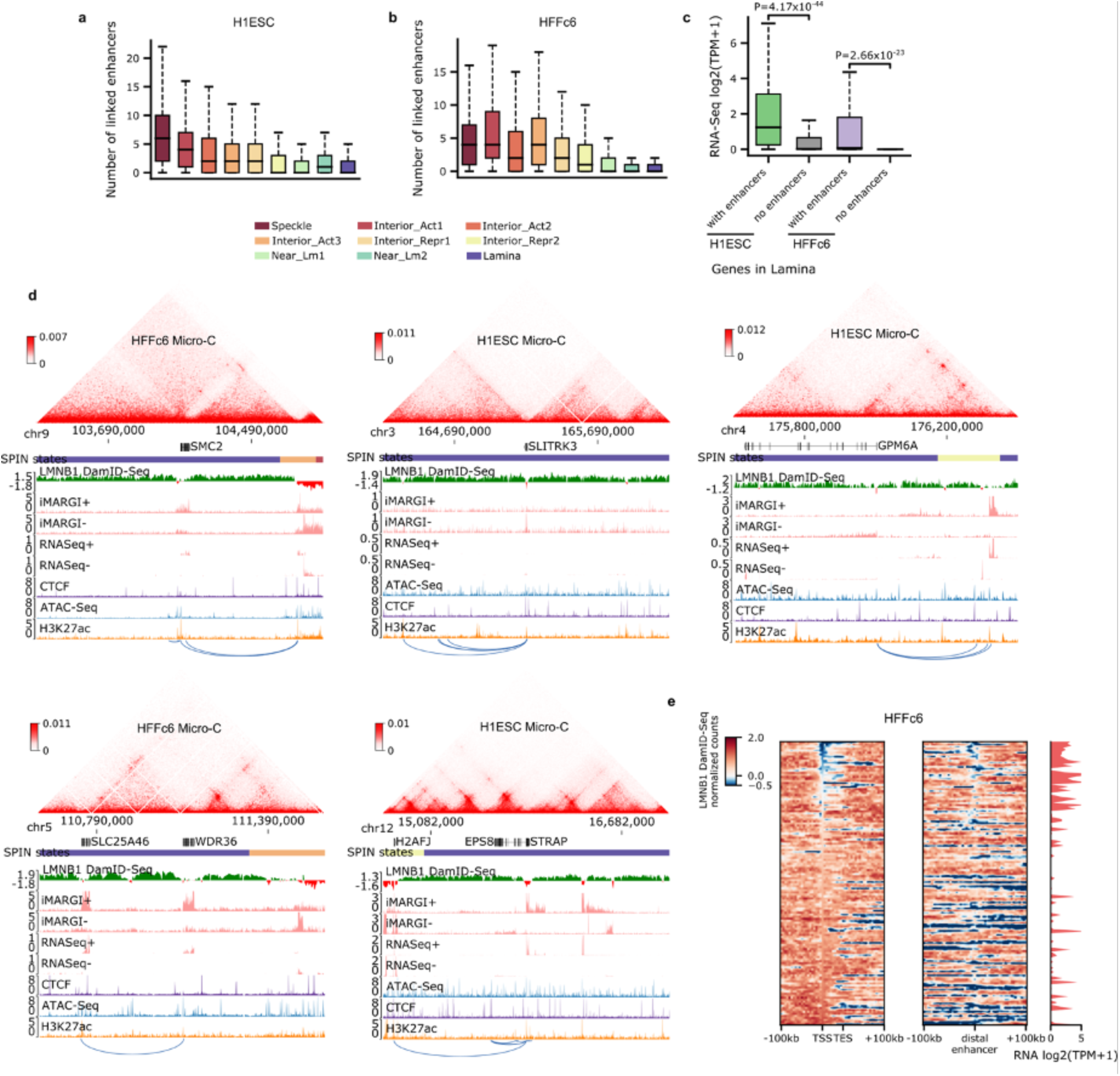
Enhancer-promoter loops within nuclear lamina and their relationships with gene regulation. For each boxplot, the center line indicates the median, the box limits represent the upper and lower quartiles, and the box whiskers extend to 1.5 times the interquartile range above and below the upper and lower quartiles, respectively. **a-b.** The distributions of the number of interacting enhancers for genes in different SPIN states in H1-hESC (a) and HFFc6 (b). **c.** Comparisons of transcription levels for genes with or without interacting enhancers in the Lamina SPIN state. The *P* values were calculated using the two-sided Mann-Whitney U test. **d.** Examples showing expressed genes, and their interacting enhancers are usually synergistically looped out of nuclear lamina to facilitate gene regulation in lamina. The blue arcs represent chromatin loops linking the gene in the center of each region with distal enhancers. **e.** Lamin-B1 DamID-seq signals surrounding lamina-associated genes and their interacting enhancers in HFFc6. Only genes with interacting enhancers in the Lamina SPIN state are included in this plot. TSS, transcription start sites. TES, transcription end sites. TPM, transcripts per kilobase million.

#### House-keeping genes are engaged in extensive and dynamic enhancer-promoter loops

Using RNA-Seq data derived from 116 human tissues or cell lines (See Supplemental Methods for data sources), we observed a strong correlation between the number of interacting enhancers and the number of tissues expressing a gene. While most genes lacking interacting enhancers are tissue-specific, those interacting with more than 10 enhancers in either H1-hESC (Figure 7b) or HFFc6 (Extended Data Figure 6d) are notably enriched with house-keeping genes (Supplemental Methods). Indeed, among the 2,175 house-keeping genes annotated in the HRT Atlas v1.0 database ^100^, we found that over 90% of them have interactions with at least one distal enhancer in both H1-hESC and HFFc6 (Figure 7c). Most of these enhancer-promoter loops are detectable in both Pol II ChIA-PET and H3K4me3 PLAC-Seq data, indicating their association with active transcription (Extended Data Figure 6e). In the 3D modeling section above, we categorized the house-keeping genes into two classes: “class I” genes exhibit the highest overall speckle associations, whereas “class II” genes exhibit minimal speckle associations (Figure 5j-m). Here, we observed that “class II” house-keeping genes have significantly fewer interacting enhancers in both cell types (Extended Data Figure 6f), consistent with our previous observations that the speckle local environment typically harbors more enhancers (Figure 5k-l). Interestingly, we observed extensive enhancer-promoter looping differences for house-keeping genes between H1-hESC and HFFc6 cells (Figure 7d; Supplemental Figure 8). For example, the EIF1 gene promoter interacts with dramatically different enhancer regions between H1-ESC and HFFc6 cells. We further interrogated each enhancer-promoter pair for whether the loop interaction and the distal enhancer are specific to one cell line or can be observed in both cell lines. As shown in Figure 7e, 80.9% of the pairs were unique to each cell line, suggesting the dynamic nature of chromatin looping between house-keeping genes and distal enhancers.

It has been shown that house-keeping genes usually contain strong promoters that are less responsive to distal enhancers ^101^. Based on CRISPR perturbation experiments and machine learning models, a recent study by the ENCODE consortium reported that house-keeping genes exhibit fewer regulatory interactions with enhancers compared to cell-specific genes, which appears to challenge our conclusions ^7^. To consolidate our observations, we extended our analysis to include 32 additional cell lines or primary cells with all the necessary data available from the ENCODE data portal. Given the variation in chromatin interaction data across samples in terms of sequencing depths and the number of detected enhancer-promoter loops, we categorized genes into 11 groups based on the percentile of the number of interacting enhancers in each sample. Consistent with our observations in H1-hESC and HFFc6, the extended analysis confirms that genes interacting with more enhancers have a higher level of enrichment for house-keeping genes (Extended Data Figures 6g). Moreover, we found that a gene is more likely to be a house-keeping gene if it exhibits extensive interactions with distal enhancers across a larger number of samples (Extended Data Figure 6h). It is important to note that the enhancer-promoter loops detected in our analysis represent physical interactions with high intensity, which may not all have functional roles in gene regulation. Interactions with multiple enhancers can lead to redundancy, which may complicate detecting functional enhancer-promoter pairs using experimental approaches where individual enhancers are tested one at the time. Further experiments are needed to investigate how multiple enhancer-promoter loops are coordinated to promote gene transcription, and whether the extensive number of enhancer-promoter loops are the cause or consequence of strong transcriptional activity of promoters, and whether these interactions can contribute to robustness of house-keeping gene transcription in different cell types.

#### Enhancer-promoter loops near the nuclear lamina

Lamina-associated domains (LADs) provide an overall repressive environment where genes are generally not expressed, although some genes can escape this repression, with their promoters usually exhibiting weaker interactions with the nuclear lamina ^102^. Consistent with a recent study ^103^, we observed that, albeit less frequently than in other nuclear environments, genes located within LADs can also interact with distal enhancers (Extended Data Figure 7a-b), and these genes are more likely to be expressed (Extended Data Figure 7c). Upon inspection of the Lamin B1 DamID-Seq signals surrounding several of these genes and their interacting enhancers (Figure 7f and Extended Data Figure 7d), we noted that both the promoters and interacting enhancers are located in small regions locally enriched for active marks and depleted for Lamin B1-DamID signals. This pattern suggests the hypothesis that these genes need to be locally looped out of the nuclear lamina to establish functional chromatin interactions (Figure 7g and Extended Data Figure 7e).

#### Relationship among A/B compartments, SPIN states, TADs/subTADs, loops, and replication timing

We next set out to further understand the genome’s structure-function relationship beyond transcription and enhancer-promoter loops. We integrated A/B compartments, SPIN states, TADs, subTADs, and loops to understand the links among specific structural categories of TADs/subTADs with replication timing during S-phase (Figure 8a).

We first classified the SPIN states in H1-hESCs as either co-registered, nested within, or encompassing A and B compartments (Figure 8b). Nearly all SPINs representing speckles and interior active 1/2/3 regions are embedded within A compartments, whereas the lamina SPIN state is embedded within or co-registered with B compartments (Figure 8b). We examined the size differences among distinct genome folding features (Supplemental Figure 9a-c). Although the size of A/B compartments span a larger dynamic range compared to other folding features, we observed that the A/B compartments that encompass or co-register with SPINs are larger (range: 300 kb - 2 Mb) and span more than 80% of the human genome (Supplemental Figure 9c). Together, these data indicate that SPINs more interior to the nucleus are more likely to be embedded within A compartments with active chromatin modifications, whereas SPINs localized to the periphery are most likely to be embedded within B compartments and inactive chromatin (Figure 8b).

We hypothesized that SPIN states may further subdivide compartments by discrete genomic domains. Noteworthy, the near lamina 1/2 and interior repressive 1/2 SPIN states are embedded within both A and B compartments, suggesting that subsets may have different regulatory functionality depending on their compartment status. To test our hypothesis, we examined the link among SPINs, replication timing, and transcription after accounting for their A/B compartment distribution using 16-fraction Repli-seq, RNA-seq, and iMargi genomics data sets (Figure 8c-d, Extended Data Figure 8a-b). As previously reported, A compartments were largely early replicating and enriched for highly expressed genes, whereas B compartments were late replicating and depleted of actively transcribed genes (Figure 8c-d, Extended Data Figure 8a-b). We observed that Speckle and Interior Active 1 SPINs exhibited further enrichment for highly transcribed genes and earlier replication timing, suggesting they represent local functional units within larger A compartments (Figure 8c-d). We observed a similar pattern for the lamina SPIN state falling largely in or co-registering with compartment B, as it showed largely depleted transcription and even later replication timing compared to the B compartment expectation (Figure 8c,e; Extended Data Figure 8b). Interior active 2/3 SPINs within A compartments did not show clear differences from the larger A compartments. It is noteworthy that certain specific SPIN states which are embedded in either A or B compartments tend to show both transcription and replication timing that is opposite of expectation. Interior repressive 1/2 and near lamina 1/2 SPINs show later replication and depleted gene expression compared to their A compartment background. Moreover, interior repressive 1/2 show enriched gene expression as well as early replication timing when nested in B compartments. These may be hESCs domains that are poised to switch either RT or compartment to re-establish canonical correlations following differentiation ^46^. Altogether, these data reveal that SPIN states represent local neighborhoods of transcription and replication timing that can depart from the global A/B compartment patterns and indicate that the compartment distribution of SPINs should be accounted for when assessing their functional impact.

**Figure 8.**
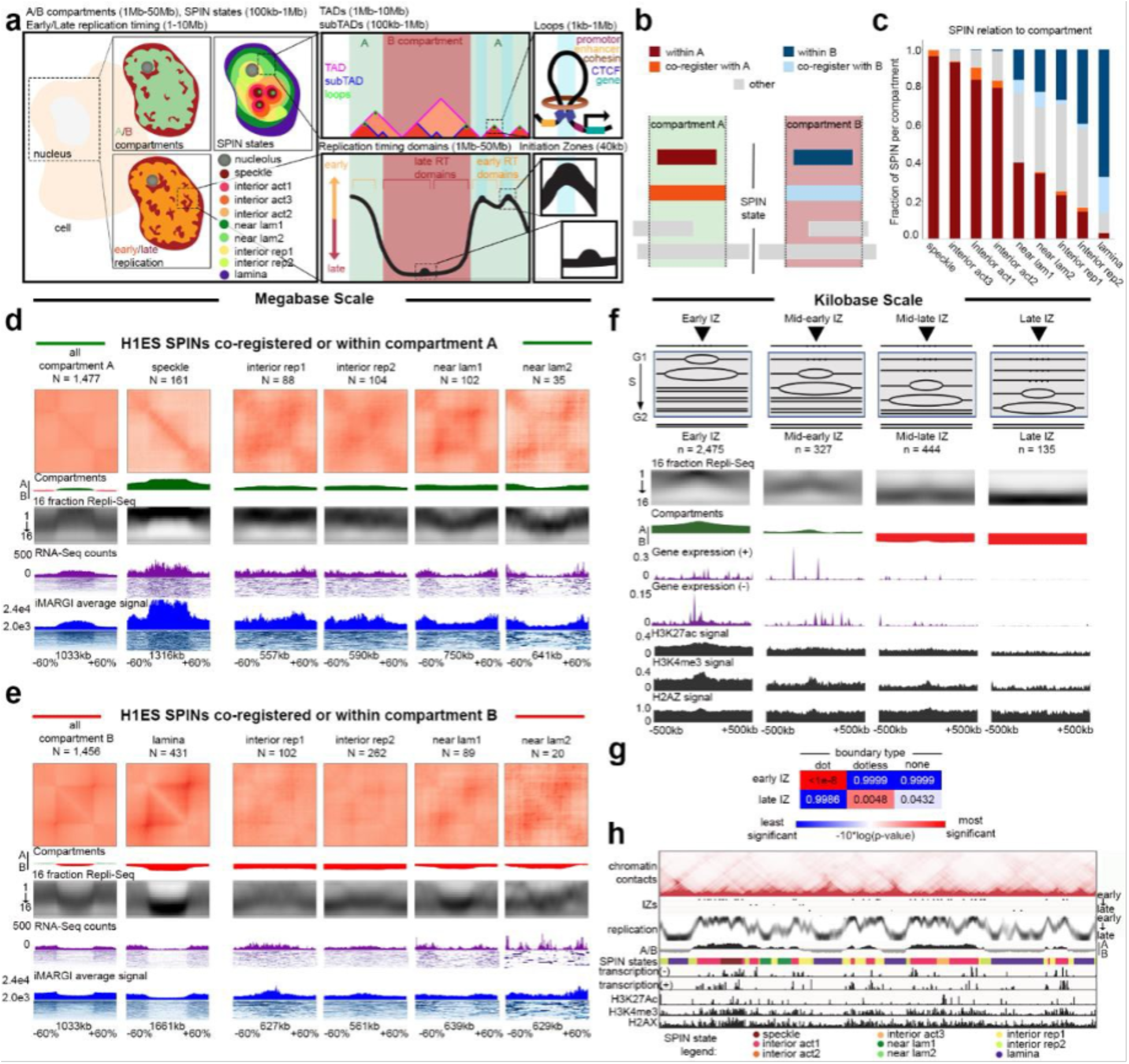
A/B compartments and SPIN states represent subnuclear regions of distinct replication timing and gene expression. **a.** Schematic of human genome folding into A/B compartments, SPIN states, TADs, subTADs, and loops integrated with early/late replication timing and initiation zones. **b.** Intersection of SPIN states with compartments. SPIN states were classified as either fully embedded within A/B compartments (within), co-registering A/B compartments (co-register), or partially-overlapping (other) **c.** Fraction of each SPIN state co-registered or nested within A/B compartments in H1-hESCs. **d,e.** Averaged Hi-C, replication timing (16 fraction Repli-seq), nascent transcription (iMargi), and total mRNA (RNA-seq) signal is plotted for h1ESCs at all A/B compartments (column 1) or co-registered/nested within selected SPIN states in A/B compartments (columns 2-6). Data is plotted as the average signal across SPIN states. The genomic intervals representing SPINs +/- flanks of 60% of the SPIN size are stretched laterally to scale by size. **d.** All A compartments or selected SPINS co-registered/within compartment A. **e.** All B compartments or selected SPINS co-registered/within compartment B. **f.** Average chromatin landscape at IZs in H1ESC. IZs have been grouped depending on their replication timing (RT). Tracks represent the high-resolution replication timing, chromatin compartments, expression and histone marks. **g.** We computed right-tailed, one-tailed empirical p-values using a resampling test with size and A/B compartment-matched null IZs for the intersection of Early and Late S phase IZs with dot boundaries, dotless boundaries, and no boundaries. **h.** Example of chromatin profiles around IZs (portion of chr2 from 20Mb to 58Mb). Tracks represents the chromatin contacts, 4 groups of IZs depending on their RT, the high-resolution replication timing, chromatin compartments, the SPIN states, Expression (minus and plus strands), H3K27Ac, H3K4me3 and H2AX.

We next explored the functional patterns across TADs/loops after accounting for their larger compartment and SPIN state environment. We identified TADs and nested subTADs in H1- hESCs using 3DNetMod and stratified the domains by looping structural features (so-called ‘dot and dotless TADs/subTADs’) as previously reported ^104^ (Extended Data Figure 9a). We found that more than half of Dot and Dotless TADs/subTADs are nested within a single SPIN state (Extended Data Figure 9b,c). To evaluate the interplay of domains and SPINs, we focused on TADs and subTADs that are nested within or co-registered with SPINs embedded within A or B compartments. We observed that nearly all Dot and Dotless TADs/subTADs resemble the replication timing of the larger SPIN state or A/B compartment and do not exhibit clear local replication timing neighborhoods (Extended Data Figure 9d-g). Dot and Dotless TADs/subTADs within active SPINs (Speckles, Interior Active 1,2,3) show enrichment of local co-regulated gene expression domains and peaks of active genes localized at both boundaries (Extended Data Figure 9d-g). Dot domains have stronger enrichment of gene expression at boundaries than Dotless domains, and this pattern is more apparent when the Dot domains are in A compartments. By contrast, dot and dotless TADs/subTADs within repressive SPINs largely do not show enrichment of expressed genes at boundaries. These data together indicate that the Mb scale folding patterns of SPIN states and compartments much more closely resemble replication timing domains compared to TADs/subTADs. TAD/subTAD boundaries, but not SPIN/compartment boundaries, are enriched for actively transcribed genes.

**Extended Data Figure 8.**
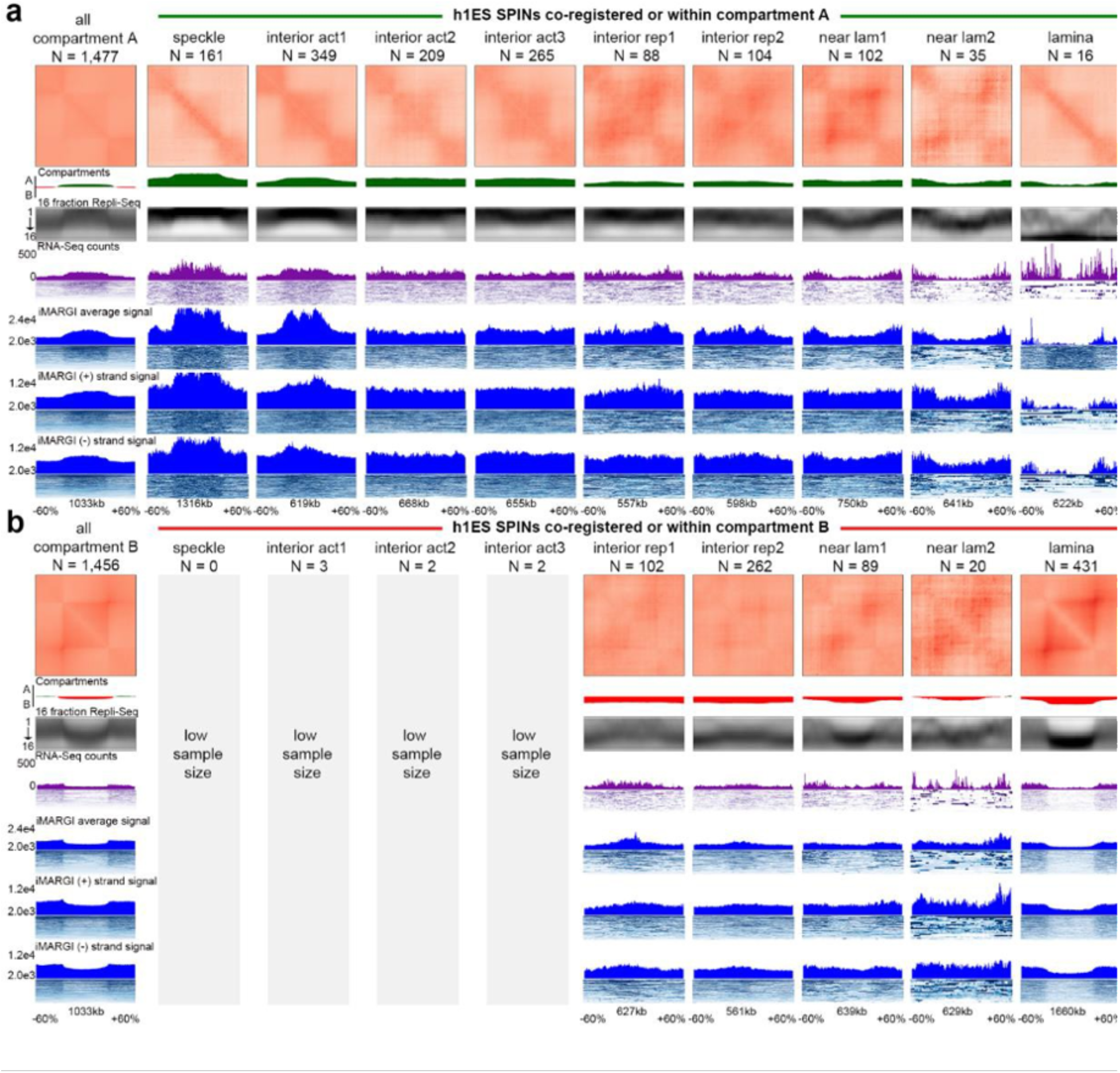
Compartment and SPIN integration with replication timing, RNA-seq, and nascent transcripts from iMargi. a, **b.** Averaged Hi-C, replication timing (16 fraction Repli-seq), nascent transcription (iMargi), and mRNA levels (RNA-seq) for h1ESCs at all A/B compartments (column 1) and SPIN states either co-registered or co-localized within A/B compartments (columns 2-10). All genomics data is plotted as the average signal across all genomic intervals representing SPINs in a particular column. SPIN genomic intervals of (SPIN genomic interval +/- flanks of 60% of the size of the genomic interval) are stretched laterally to scale by size before average signal is computed. **a.** All A compartments or select SPINS co-registered or within compartment A and **b.** All B compartments or select SPINS co-registered or within compartment B. Tracks show pileups in h1ESC for Hi-C Aggregate-Peak-Analysis (APA), A/B compartment, 16 fraction Repli-seq, median RNA-seq signal, condensed RNA-seq reads, median averaged iMARGI (+) and iMARGI (-) signal, condensed iMARGI (+) and iMARGI (-) reads, median iMARGI (+) signal, condensed iMARGI (+) reads, median iMARGI (-) signal, and condensed iMARGI (-) reads.

A distinct essential feature of genome function is the replication initiation zone (IZ). These are ∼50 kb regions of the genome within which replication initiates at one or more of many potential sites. Initiation of replication within each IZ occurs once in each 5-20% of cell cycles ^105^, with the probability of an IZ firing early in S phase regulated by *cis*-acting elements termed early replication control elements (ERCEs) ^106^. Interestingly, a specific subset of TAD boundaries, active gene-enriched dot TAD boundaries, are enriched for early firing IZs in a cohesin-dependent manner ^104^, revealing a biologically significant structure-function relationship unique to this set of TAD boundaries. Extended Data Figure 10 shows that, when IZs are stratified by their timing of firing in early, early-mid, mid-late, and late S phase in H1-hESC, HCT116 and F121-9 mouse ESCs (as in ^75^) and aligned to insulation score of replication timing-matched random sequences, a relationship to TAD boundaries is difficult to identify (Extended Data Figure 10a-c). By contrast, the canonical correlations of replication timing to chromatin accessibility, gene expression and active histone marks are evident (Extended Data Figure 10d-l). Thus, IZs are enriched at only a specialized subset of TAD boundaries suggesting that, when assigning function to TADs, we need to appreciate their potential functional diversity.

Overall, our data indicate that the relationship between higher-order chromatin structure and DNA replication is dependent on the length scale of the folding feature. Mb scale folding patterns of SPIN states and compartments best correlate with replication timing domains. By contrast, TADs/subTADs appear to reflect replication timing of the larger compartment/spin in which they reside, and do not show clear local replication timing neighborhoods but rather enrichment of gene expression at boundaries. While IZs are not globally enriched at TAD boundaries, they are highly enriched at dot TAD boundaries that harbor actively transcribed genes.

**Extended Data Figure 9.**
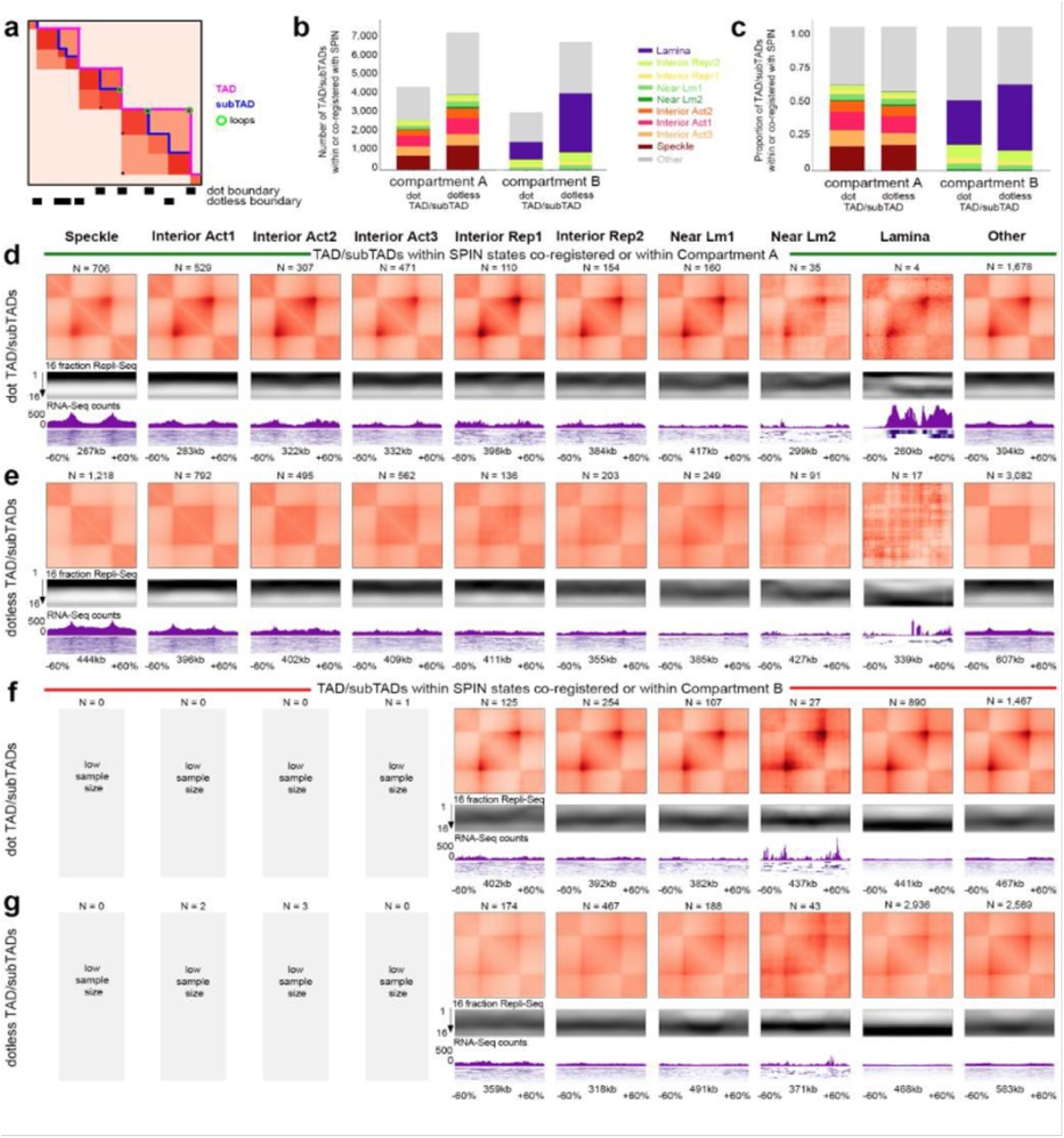
Compartment, SPIN, and TAD integration with replication timing, RNA-seq, and nascent transcripts from iMargi. **a.** Schematic depicting TAD (pink) and subTADs (blue) domains and loops (green circle). Dot domains contain a loop at the domain apex and dot boundaries. Dotless domains do not contain a loop at the apex and thus only dotless boundaries. **b., c.** Dot and dotless TAD/subTAD domains that are within or co-register with a SPIN state contained within or co-registering with A/B compartments. **b.** Number of and **c.** proportion of SPINs stratified by A/B compartment and presence of corner-dot TAD/subTAD domains. **d.-g.** Averaged Hi-C, replication timing (16 fraction Repli-seq), mRNA levels (RNA-seq) for H1-hESCs at dot and dotless TAD/subTAD domains co-registered or co-localized within all SPIN states (columns 1-9) and TAD/subTAD domains with other SPIN alignment (column 10) either co-registered or co-localized within A/B compartments. All genomics data is plotted as the average signal across all genomic intervals representing domains in a particular column. TAD/subTAD genomic intervals of (TAD/subTAD genomic interval +/- flanks of 60% of the size of the genomic interval) are stretched laterally to scale by size before average signal is computed. **d.** Dot and **e.** dotless TAD/subTAD domain co- registered or within a SPIN and co-registered or within compartment A and **f.** Dot and **g.** dotless TAD/subTAD domain co-registered or within a SPIN and co-registered or within compartment B. Tracks show pileups in H1-hESC for Hi-C Aggregate-Peak-Analysis (APA), 16 fraction Repli-seq, median RNA-seq signal, condensed RNA-seq reads.

**Extended Data Figure 10.**
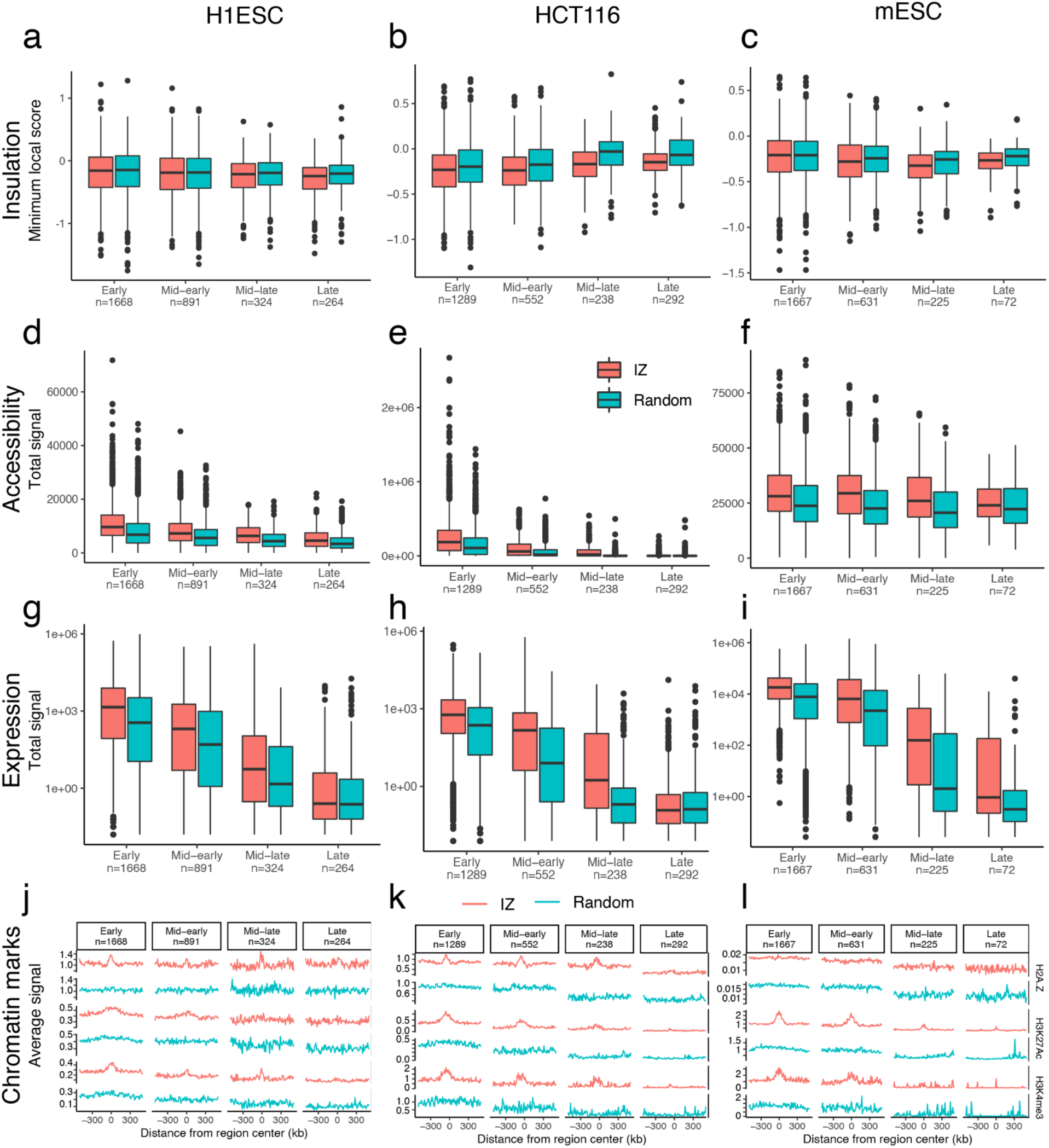
Comparison of IZ properties across cell lines. **a.-c.** Boxplots showing the minimum diamond insulation score at IZs in H1-hESC (**a**), HCT116 (**b**) and mESC (**b**). **d.-f.** Boxplots showing the total chromatin accessibility at IZs in H1-hESC (**d**), HCT116 (**e**), and mESC (**f**). **g.-i.** Boxplots showing the total RNA signal in H1-hESC (**g**), HCT116 (**h**), and mESC (**i**). **j.-l**. Average histone marks signal in 1Mb regions centered on IZs in H1-hESC (**j**), HCT116 (**k**), and mESC (**l**). Boxplots represent the median and interquartile range (IQR); whiskers mark 1.5x the IQR; data beyond 1.5x the IQR are plotted as individual points.

### Predicting chromosome folding from sequence, and relating disease variants to altered chromatin structure

Currently, several deep learning methods use high-resolution chromatin capture data (Hi-C and Micro-C) to predict 3D genome folding from sequence alone ^94,107–110^, epigenetic features alone ^111,112^, or sequence plus epigenetic features ^113^. Additionally, polymer physics-based machine learning methods (PRISMR) trained on only wild-type Hi-C data, were shown to correctly predict the impact of disease associated structural variants on chromosome architecture and specifically on the rewiring of promoter-enhancer contacts, as validated by independent experiments in cells bearing those mutations ^114^. The high accuracy of these models allows the decoding of sequence determinants of genome folding using explainable AI techniques, such as importance scores ^115^ and *in silico* mutagenesis ^116^. These methods can be applied to interpret synthetic manipulations of the reference genome sequence or observed genetic variants with software such as SuPreMo ^117^.

To demonstrate these capabilities, we first trained two cell-type specific models on 4DN H1- hESCs and HFFc6 Micro-C data using the Akita architecture ^107^. Next, we used the HFFc6 model to predict the effect of a deletion at the TAL1 locus (Figure 9a) and confirmed that predicted changes in chromatin interactions mirror those that were observed when the deletion was made in HEK293T cells using CRISPR-Cas9 genome editing ^118^. Finally, we computationally mutated motifs of cell type specific transcription factors at TAD boundaries unique to each cell type and quantified their effects on genome folding (Figure 9b). We found that the H1-hESC model is more sensitive to mutation of motifs for the embryonic stem cell factors POU2F1::SOX2 ^119,120^, while the HFFc6 model is more sensitive to the fibroblast factors FOSL1::JUND ^121^. These findings suggest that deep learning models and explainable AI can be used to screen DNA sequences at scale for the unbiased discovery of genome folding mechanisms and their associations with genome function. The potential of this approach is demonstrated by recent studies of cancer associated variants ^107,117^ and an unbiased genome-wide screen of synthetic variants that revealed the importance of repetitive elements in genome- folding ^116^.

**Figure 9.**
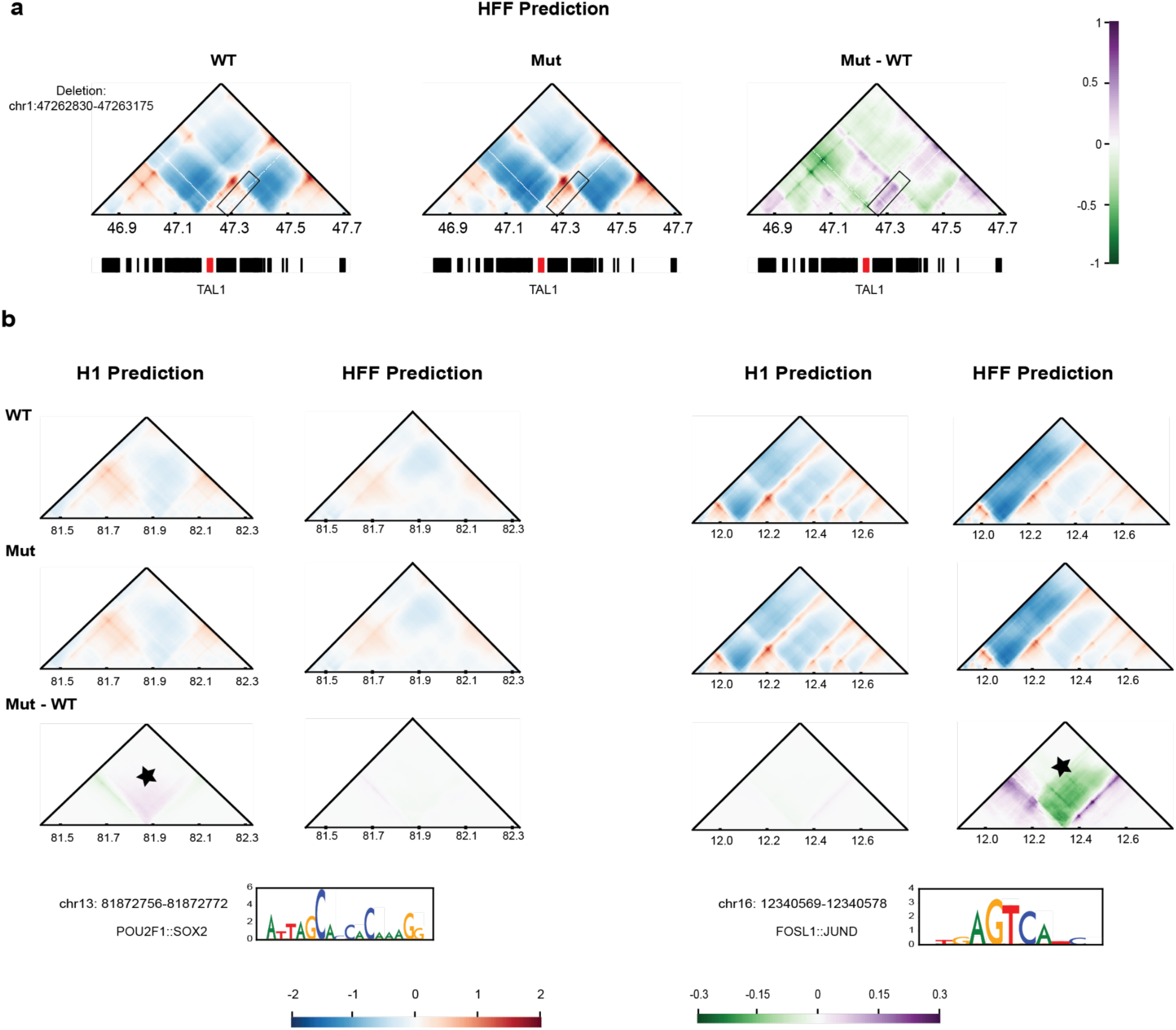
Predicting the effect of genomic variants on 3D genome folding with deep learning. **a.** Example of a 345 bp deletion (chr1: 47262830-47263175) at the TAL1 locus. Contact maps (log(observed/expected)) were predicted for ∼1Mb regions with a model trained on HFFc6 Micro-C data using the Akita architecture ^107^. Maps for the reference human genome sequence (WT) and the in silico mutated sequence (Mut; at center) are plotted on a color scale where red indicates higher than expected interaction frequencies, and blue indicates lower than expected given genomic distance. The effect of the deletion (Mut - WT) is plotted on a color scale where purple indicates increased and green decreased chromatin interactions. Genes in the locus are plotted below the contact maps with TAL1 highlighted in red. The deleted region has a CTCF binding site and is located in a TAD boundary. Mirroring the experimental deletion in HEK293T cells (Hnisz et al., 2016), our model predicted increased contact frequency between TAL1 and adjacent regions (black rectangle). **b.** In silico mutation of transcription factor motifs (replacing motifs with random sequences) affects deep learning predictions of nearby chromatin interactions. An example POU2F1::SOX2 motif (left, chr13: 81872756-81872772) and FOSL1::JUND motif (right, chr16: 12340569-12340578) were generated using models with the Akita architecture trained on H1-hESC or HFFc6 Micro-C data, respectively. Motif logos generated via model importance scores using DeepExplainer ^115^ are shown below the maps. Color scales are the same as in (**a**), and motif sites are centered on the contact maps. Star symbols indicate regions with altered chromatin interaction predictions.

## Discussion

We present an integrated view of the human 4DN nucleome, and describe connections between chromosome folding and looping, nuclear positioning, proximity to nuclear bodies, cell-to-cell variation in organization, and genomic functions such as transcription and replication.

The work provides tangible results. First, the extensive integration of a range of genomic datasets reporting on the spatial organization of the human genome in two cell types allowed us to benchmark these methods and to show which methods are best for specific inquiries. We find that all methods have their own strengths and weaknesses. Compartmentalization is most effectively detected using SPRITE and Hi-C, while looping interactions are best detected using Micro-C (especially structural loops), and enrichment-based assays such as PLAC-seq and ChIA-PET (gene expression related loops). The longer capture radius of SPRITE and GAM allow detection of colocalization of loci around larger sub-nuclear bodies. GAM, and single cell Hi-C can be applied when rare or mixed cell types are studied.

Second, these genomic datasets together provide a large catalog of looping interactions between specific cis-elements, including CTCF-CTCF interactions, and interactions among promoters and enhancers for two widely used cell types. Besides providing a resource that can be mined for future studies, this collection suggests mechanistic connections between the 4D nucleome and genome regulation. For instance, we find that cohesin is enriched at a large proportion of anchors of all types of loops, suggesting that cohesin, and possibly loop extrusion, is involved in their formation. However, this is not directly demonstrated through perturbation experiments, and other mechanisms most likely will also play roles. Looping interactions with distal enhancers is strongly correlated with gene expression. Housekeeping genes are particularly prone to interact physically with distal enhancers, but with different sets of enhancers in different cell types. It is possible that this promiscuity in long-range interactions allows these genes to be expressed in many different cell types. However, we cannot rule out these are non-functional interactions that reflect the active transcriptional state of these genes.

Third, integration of the panel of genomic datasets allowed the generation of a detailed annotation track of spatial information, SPIN states, along the genome. This linear representation of 4DN nucleome information will greatly facilitate integration of spatial genome data with other genomic datasets obtained in the larger community. In one example, our analysis of SPIN states, compartments and TADs, and DNA replication timing shows that one needs to take the heterogeneity of TADs into account when assigning a biological function to their structure. Indeed, we expect many novel functions to be assigned to specific subsets of TADs.

Finally, we generated ensembles of spatial models through integrating and combining these datasets. Detailed analysis of these models starts to place genome functions such as transcription and replication in three-dimensional context, e.g., in relation to nuclear bodies such as the nuclear lamina and nuclear speckles. The models also reflect the extensive cell-to-cell heterogeneity that defines the 4D nucleome, as also detected with single cell assays. These models can provide a powerful resource for future studies, e.g., to benchmark single cell assays and imaging-based assays currently ongoing in the 4D nucleome project and elsewhere. One example is that the models highlight the existence of two types of housekeeping genes that occupy two quite distinct sub-nuclear neighborhoods.

### Future perspective

We present results from an integrated project with many participating groups that was the focus of the first phase of the international 4D Nucleome Project. In the current ongoing second phase of the 4D Nucleome Project, a focus is on integrating genomic datasets with imaging data, development and application of a range of multi-omic single cell datasets, and the analysis of 4D nucleome changes during development and in disease ^28,29^. In addition, one exciting new direction of the project is the development of approaches to use 4D nucleome data to predict cell-type specific chromosome conformation from sequence. One important application of these approaches is the identification of cis-elements, and thereby potentially new mechanisms, that drive chromosome folding. These approaches can predict effects of genetic, and possibly disease-related, variants on chromosome folding and chromatin looping between elements, and thus start to relate alterations in chromosome folding to disease. While existing deep learning models rely on one or a small set of data modalities (e.g., Hi-C and Micro-C), in the future these models can be trained on richer models of the 4D Nucleome, based on integration of multiple datatypes as described here, and by integration of imaging and single cell multi-omics data that are currently being generated by the 4D nucleome project.

Taken together, the rich datasets on genome folding presented here, and their integration, reveal a detailed view of the living physical human genome as it is organized inside cells, and provide a foundation for future deep exploration of the structure and function of the genomes, in humans and across the tree of life, in normal and disease states.

## Acknowledgement

This work has been collectively funded by grants from the National Institutes of Health Common Fund (The 4D Nucleome Project): U01HL157989, U54DK107965, UM1HG011593, U54DK107981, UM1HG011593, U01CA200059, U01CA200059, U54DK107977, UM1HG011585, U01DA052769, U01CA200147, U01CA200060, U01DA040612, U01DK127420, U01HL130007, U54DK107967, U54DK107979, UM1HG011531, U54DK107980, HG011536, U01HL129998, U01DA052715, U01DK127405.

## Author contributions

### Project coordinators

Job Dekker, Feng Yue, William Noble

### Topic leaders

Job Dekker, Feng Yue, William Noble, Xiaotao Wang, Jian Ma, Frank Alber, Mario Nicodemi, Bing Ren, Sheng Zhong, Jennifer Phillips-Cremins, David M. Gilbert, Katherine S. Pollard

### Manuscript writing and editing

Job Dekker, Feng Yue, William Noble, Jian Ma, Bing Ren, Ting Wang, Yun Li, Andrew S. Belmont, Katherine S. Pollard, Frank Alber, Mario Nicodemi, Shuzhen Kuang, Yang Zhang, Ruochi Zhang, Xingzhao Wen, Xiaotao Wang

### Data production (execution of data production experiments, experiment coordination, and supervision)

Liyan Yang, Johan H. Gibcus, Nils Krietenstein, Oliver J. Rando, Jie Xu, Derek H. Janssens, Steven Henikoff, Alexander Kukalev, Andréa Willemin, Warren Winick-Ng, Rieke Kempfer, Bing Ren, Miao Yu, Pradeep Kumar, Liguo Zhang, Andrew S. Belmont, David M. Gilbert, Takayo Sasaki, Tom van Schaik, Laura Brueckner, Daan Peric-Hupkes, Bas van Steensel, Sheng Zhong, Ping Wang, Haoxi Chai, Yijun Ruan, Ran Zhang, William Noble, Sofia Quinodoz, Mitchell Guttman, Ana Pombo

### Methods Benchmarking

Job Dekker, Feng Yue, Betul Akgol Oksuz, Xiaotao Wang, Johan H. Gibcus, Sergey V. Venev, Dariusz Plewczynski, Mario Nicodemi, Alexander Kukalev, Ibai Irastorza Azcarate, Dominik Szabó, Christoph J Thieme, Teresa Szczepińska, Ana Pombo

### Annotation of spatial chromatin compartmentalization through integrative modeling

Yang Zhang, Ran Zhang, Mateusz Chiliński, Kaustav Sengupta, Asli Yildirim, Mattia Conte, Andrea Esposito, Alex Abraham, Ruochi Zhang, Yuchuan Wang, Andrew S. Belmont, William Noble, Xingzhao Wen, Qiuyang Wu, Wenxin Zhao, Shu Chien, Yang Yang, Sheng Zhong, Jie Liu, Dariusz Plewczynski, Mario Nicodemi, Frank Alber, Jian Ma

### 3D modeling

Ye Wang, Asli Yildirim, Lorenzo Boninsegna, Yuxiang Zhan, Mateusz Chiliński, Kaustav Sengupta, Mattia Conte, Andrea Esposito, Alex Abraham, Yang Zhang, Dariusz Plewczynski, Mario Nicodemi, Jian Ma, Frank Alber

### Single cell 3D genome analysis

Ruochi Zhang, Ran Zhang, Mattia Conte, Andrea Esposito, Alex Abraham, Miao Yu, Lindsay Lee, Ming Hu, Bing Ren, Mario Nicodemi, Jian Ma

### 3D genome and genome function

Xiaotao Wang, Ye Wang, Daniel J. Emerson, R. Jordan Barnett, Miriam K. Minsk, Ashley L. Cook, Jennifer E. Phillips-Cremins, Claire Marchal, Peiyao Zhao, David M. Gilbert, Sheng Zhong, Xingzhao Wen, Yuan Liu, Yang Zhang, Jian Ma

### Predicting chromosome folding from sequence

Shuzhen Kuang, Katherine S. Pollard

### Data coordination and visualization

Peter Park, Burak Alver, Andrew Schroeder, Rahi Navelkar, Clara Bakker, William Ronchetti, Shannon Ehmsen, Alex Veit, Nils Gehlenborg, Ting Wang, Daofeng Li, Yang Zhang, Jian Ma

## References

1 ENCODE-Project-Consortium. An integrated encyclopedia of DNA elements in the human genome. Nature 489, 57–74 (2012).

2 Consortium, E. P. et al. Expanded encyclopaedias of DNA elements in the human and mouse genomes. Nature 583, 699–710 (2020). 10.1038/s41586-020-2493-4

3 Furlong, E. E. M. & Levine, M. Developmental enhancers and chromosome topology. Science 361, 1341–1345 (2018). 10.1126/science.aau0320

4 Robson, M. I., Ringel, A. R. & Mundlos, S. Regulatory Landscaping: How Enhancer-Promoter Communication Is Sculpted in 3D. Mol Cell 74, 1110–1122 (2019). 10.1016/j.molcel.2019.05.032

5 Galouzis, C. C. & Furlong, E. E. M. Regulating specificity in enhancer-promoter communication. Curr Opin Cell Biol 75, 102065 (2022). 10.1016/j.ceb.2022.01.010

6 Popay, T. M. & Dixon, J. R. Coming full circle: On the origin and evolution of the looping model for enhancer-promoter communication. J Biol Chem 298, 102117 (2022). 10.1016/j.jbc.2022.102117

7 Gschwind, A. R. et al. An encyclopedia of enhancer-gene regulatory interactions in the human genome. bioRxiv (2023). 10.1101/2023.11.09.563812

8 Bonev, B. & Cavalli, G. Organization and function of the 3D genome. Nat Rev Genet 17, 772 (2016). 10.1038/nrg.2016.147

9 Mirny, L. & Dekker, J. Mechanisms of Chromosome Folding and Nuclear Organization: Their Interplay and Open Questions. Cold Spring Harb Perspect Biol 14 (2022). 10.1101/cshperspect.a040147

10 Belmont, A. S. Nuclear Compartments: An Incomplete Primer to Nuclear Compartments, Bodies, and Genome Organization Relative to Nuclear Architecture. Cold Spring Harb Perspect Biol 14 (2022). 10.1101/cshperspect.a041268

11 Misteli, T. The Self-Organizing Genome: Principles of Genome Architecture and Function. Cell 183, 28–45 (2020). 10.1016/j.cell.2020.09.014

12 Oberbeckmann, E. & Oudelaar, A. M. Genome organization across scales: mechanistic insights from in vitro reconstitution studies. Biochem Soc Trans 52, 793–802 (2024). 10.1042/BST20230883

13 Hoencamp, C. & Rowland, B. D. Genome control by SMC complexes. Nat Rev Mol Cell Biol 24, 633–650 (2023). 10.1038/s41580-023-00609-8

14 Busslinger, G. A. et al. Cohesin is positioned in mammalian genomes by transcription, CTCF and Wapl. Nature 544, 503–507 (2017). 10.1038/nature22063

15 Valton, A. L. et al. A cohesin traffic pattern genetically linked to gene regulation. Nat Struct Mol Biol 29, 1239–1251 (2022). 10.1038/s41594-022-00890-9

16 Rinzema, N. J. et al. Building regulatory landscapes reveals that an enhancer can recruit cohesin to create contact domains, engage CTCF sites and activate distant genes. Nat Struct Mol Biol 29, 563–574 (2022). 10.1038/s41594-022-00787-7

17 Nora, E. P. et al. Spatial partitioning of the regulatory landscape of the X-inactivation centre. Nature 485, 381–385 (2012).

18 Dixon, J. R. et al. Topological domains in mammalian genomes identified by analysis of chromatin interactions. Nature 485, 376–380 (2012).

19 Fudenberg, G. et al. Formation of chromosomal domains by loop extrusion. Cell Rep. 15, 2038–2049 (2016).

20 Rao, S. S. P. et al. Cohesin Loss Eliminates All Loop Domains. Cell 171, 305–320 e324 (2017). 10.1016/j.cell.2017.09.026

21 Rao, S. S. P. et al. A 3D map of the human genome at kilobase resolution reveals principles of chromatin looping. Cell 159, 1665–1680 (2014).

22 Sanborn, A. L. et al. Chromatin extrusion explains key features of loop and domain formation in wild-type and engineered genomes. Proc Natl Acad Sci U S A. 112, E6456–6465 (2015).

23 Crane, E. et al. Condensin-driven remodeling of X-chromosome topology during dosage compensation. Nature 523, 240–244 (2015).

24 Lieberman-Aiden, E. et al. Comprehensive mapping of long-range interactions reveals folding principles of the human genome. Science 326, 289–293 (2009).

25 Harris, H. L. et al. Chromatin alternates between A and B compartments at kilobase scale for subgenic organization. Nat Commun 14, 3303 (2023). 10.1038/s41467-023-38429-1

26 Goel, V. Y., Huseyin, M. K. & Hansen, A. S. Region Capture Micro-C reveals coalescence of enhancers and promoters into nested microcompartments. Nat Genet 55, 1048–1056 (2023). 10.1038/s41588-023-01391-1

27 Dekker, J. et al. The 4D nucleome project. Nature 549, 219–226 (2017). 10.1038/nature23884

28 Roy, A. L. et al. Elucidating the structure and function of the nucleus-The NIH Common Fund 4D Nucleome program. Mol Cell 83, 335–342 (2023). 10.1016/j.molcel.2022.12.025

29 Dekker, J. et al. Spatial and temporal organization of the genome: Current state and future aims of the 4D nucleome project. Mol Cell 83, 2624–2640 (2023). 10.1016/j.molcel.2023.06.018

30 Dekker, J., Rippe, K., Dekker, M. & Kleckner, N. Capturing Chromosome Conformation. Science 295, 1306-1311 (2002).

31 Hsieh, T. S. et al. Mapping nucleosome resolution chromosome folding in yeast by Micro-C. . Cell 162, 108–119 (2015).

32 Krietenstein, N., et al. Ultrastructural details of mammalian chromosome architecture. Mol. Cell in press (2020).

33 Quinodoz, S. A. et al. Higher-Order Inter-chromosomal Hubs Shape 3D Genome Organization in the Nucleus. Cell 174, 744–757 e724 (2018). 10.1016/j.cell.2018.05.024

34 Beagrie, R. A. et al. Complex multi-enhancer contacts captured by genome architecture mapping. Nature 543, 519–524 (2017).

35 Fullwood, M. J. et al. An oestrogen-receptor-alpha-bound human chromatin interactome. Nature 462, 58–64 (2009).

36 Mumbach, M. R. et al. HiChIP: efficient and sensitive analysis of protein-directed genome architecture. Nat Methods 13, 919–922 (2016). 10.1038/nmeth.3999

37 Fang, R. et al. Mapping of long-range chromatin interactions by proximity ligation-assisted ChIP-seq. Cell Res 26, 1345–1348 (2016). 10.1038/cr.2016.137

38 Beagrie, R. A. et al. Multiplex-GAM: genome-wide identification of chromatin contacts yields insights overlooked by Hi-C. Nat Methods 20, 1037–1047 (2023). 10.1038/s41592-023-01903-1

39 Akgol Oksuz, B., et al. Systematic evaluation of chromosome conformation capture assays. Nat Methods 18, 1046–1055 (2021). 10.1038/s41592-021-01248-7

40 Fiorillo, L. et al. Comparison of the Hi-C, GAM and SPRITE methods using polymer models of chromatin. Nat Methods 18, 482–490 (2021). 10.1038/s41592-021-01135-1

41 Polovnikov, K. E. et al. Crumpled polymer with loops recapitulates key features of chromosome organization. Phys Rev X 13 (2023). 10.1103/physrevx.13.041029

42 Simonis, M. et al. Nuclear organization of active and inactive chromatin domains uncovered by chromosome conformation capture-on-chip (4C). Nat. Genet. 38, 1348–1354 (2006).

43 Xiong, K. & Ma, J. Revealing Hi-C subcompartments by imputing high-resolution inter-chromosomal interactions. BioRxiv doi: 10.1101/505503 (2019).

44 Spracklin, G. et al. Diverse silent chromatin states modulate genome compartmentalization and loop extrusion barriers. Nat Struct Mol Biol 30, 38–51 (2023). 10.1038/s41594-022-00892-7

45 Nora, E. P. et al. Targeted Degradation of CTCF Decouples Local Insulation of Chromosome Domains from Genomic Compartmentalization. Cell 169, 930–944 e922 (2017). 10.1016/j.cell.2017.05.004

46 Dileep, V. et al. Rapid Irreversible Transcriptional Reprogramming in Human Stem Cells Accompanied by Discordance between Replication Timing and Chromatin Compartment. Stem Cell Reports 13, 193–206 (2019). 10.1016/j.stemcr.2019.05.021

47 Solovei, I., Thanisch, K. & Feodorova, Y. How to rule the nucleus: divide et impera. Curr Opin Cell Biol 40, 47–59 (2016). 10.1016/j.ceb.2016.02.014

48 Bickmore, W. A. The spatial organization of the human genome. Annu Rev Genomics Hum Genet. 14 (2013).

49 Ryba, T., Battaglia, D., Pope, B. D., Hiratani, I. & Gilbert, D. M. Genome-scale analysis of replication timing: from bench to bioinformatics. Nat Protoc 6, 870–895 (2011). 10.1038/nprot.2011.328

50 Paulsen, J. et al. Long-range interactions between topologically associating domains shape the four-dimensional genome during differentiation. Nat Genet 51, 835–843 (2019). 10.1038/s41588-019-0392-0

51 Kosak, S. T. et al. Subnuclear compartmentalization of immunoglobulin loci during lymphocyte development. Science. 296, 158–162 (2002).

52 Solovei, I. et al. Nuclear architecture of rod photoreceptor cells adapts to vision in mammalian evolution. Cell. 137, 356–368 (2009).

53 Croft, J. A. et al. Differences in the localization and morphology of chromosomes in the human nucleus. J. Cell Biol. 145, 1119–1131 (1999).

54 Chen, Y. et al. Mapping 3D genome organization relative to nuclear compartments using TSA-Seq as a cytological ruler. J Cell Biol 217, 4025–4048 (2018). 10.1083/jcb.201807108

55 Guelen, L. et al. Domain organization of human chromosomes revealed by mapping of nuclear lamina interactions. Nature 453, 948–951 (2008).

56 Zhang, L. et al. TSA-seq reveals a largely conserved genome organization relative to nuclear speckles with small position changes tightly correlated with gene expression changes. Genome Res 31, 251–264 (2021). 10.1101/gr.266239.120

57 Kumar, P. et al. Nucleolus and centromere TSA-Seq reveals variable localization of heterochromatin in different cell types. bioRxiv (2023). 10.1101/2023.10.29.564613

58 Marchal, C. et al. Genome-wide analysis of replication timing by next-generation sequencing with E/L Repli-seq. Nat Protoc 13, 819–839 (2018). 10.1038/nprot.2017.148

59 Ryba, T. et al. Evolutionarily conserved replication timing profiles predict long-range chromatin interactions and distinguish closely related cell types. Genome Res. 20, 761–770 (2010).

60 Rivera-Mulia, J. C. et al. Dynamic changes in replication timing and gene expression during lineage specification of human pluripotent stem cells. Genome Res 25, 1091–1103 (2015). 10.1101/gr.187989.114

61 Salameh, T. J. et al. A supervised learning framework for chromatin loop detection in genome-wide contact maps. Nat Commun 11, 3428 (2020). 10.1038/s41467-020-17239-9

62 Ernst, J. & Kellis, M. ChromHMM: automating chromatin-state discovery and characterization. Nat Methods 9, 215–216 (2012).

63 Tang, Z. et al. CTCF-Mediated Human 3D Genome Architecture Reveals Chromatin Topology for Transcription. Cell 163, 1611–1627 (2015).

64 Thiecke, M. J. et al. Cohesin-Dependent and -Independent Mechanisms Mediate Chromosomal Contacts between Promoters and Enhancers. Cell Rep 32, 107929 (2020). 10.1016/j.celrep.2020.107929

65 Uyehara, C. M. & Apostolou, E. 3D enhancer-promoter interactions and multi-connected hubs: Organizational principles and functional roles. Cell Rep 42, 112068 (2023). 10.1016/j.celrep.2023.112068

66 Ngan, C. Y. et al. Chromatin interaction analyses elucidate the roles of PRC2-bound silencers in mouse development. Nat Genet 52, 264–272 (2020). 10.1038/s41588-020-0581-x

67 Kraft, K. et al. Polycomb-mediated genome architecture enables long-range spreading of H3K27 methylation. Proc Natl Acad Sci U S A 119, e2201883119 (2022). 10.1073/pnas.2201883119

68 Massett, M. E. et al. The Histone Demethylase KDM4A Is Required to Sustain H3K9me3/H3K27me3 Epigenetic States and Oncogenesis in MLL-AF9 Acute Myeloid Leukemia. Blood 132, 3879 (2018).

69 Van Rechem, C. et al. Collective regulation of chromatin modifications predicts replication timing during cell cycle. Cell Rep 37, 109799 (2021). 10.1016/j.celrep.2021.109799

70 Wang, Y. et al. SPIN reveals genome-wide landscape of nuclear compartmentalization. Genome Biol 22, 36 (2021). 10.1186/s13059-020-02253-3

71 Schreiber, J., Durham, T., Bilmes, J. & Noble, W. S. Avocado: a multi-scale deep tensor factorization method learns a latent representation of the human epigenome. Genome Biol 21, 81 (2020). 10.1186/s13059-020-01977-6

72 Schreiber, J., Durham, T., Bilmes, J. & Noble, W. S. Author Correction: Avocado: a multi-scale deep tensor factorization method learns a latent representation of the human epigenome. Genome Biol 22, 255 (2021). 10.1186/s13059-021-02470-4

73 Yu, R. et al. CTCF/cohesin organize the ground state of chromatin-nuclear speckle association. bioRxiv (2023). 10.1101/2023.07.22.550178

74 Pinello, L., Xu, J., Orkin, S. H. & Yuan, G. C. Analysis of chromatin-state plasticity identifies cell-type-specific regulators of H3K27me3 patterns. Proc Natl Acad Sci U S A 111, E344–353 (2014). 10.1073/pnas.1322570111

75 Zhao, P. A., Sasaki, T. & Gilbert, D. M. High-resolution Repli-Seq defines the temporal choreography of initiation, elongation and termination of replication in mammalian cells. Genome Biol 21, 76 (2020). 10.1186/s13059-020-01983-8

76 Yan, Z. et al. Genome-wide colocalization of RNA-DNA interactions and fusion RNA pairs. Proc Natl Acad Sci U S A 116, 3328–3337 (2019). 10.1073/pnas.1819788116

77 Wu, W. et al. Mapping RNA-chromatin interactions by sequencing with iMARGI. Nat Protoc 14, 3243–3272 (2019). 10.1038/s41596-019-0229-4

78 Calandrelli, R. et al. Genome-wide analysis of the interplay between chromatin-associated RNA and 3D genome organization in human cells. Nat Commun 14, 6519 (2023). 10.1038/s41467-023-42274-7

79 Quinodoz, S. A. et al. RNA promotes the formation of spatial compartments in the nucleus. Cell 184, 5775–5790 e5730 (2021). 10.1016/j.cell.2021.10.014

80 Liang, L. et al. Complementary Alu sequences mediate enhancer-promoter selectivity. Nature 619, 868–875 (2023). 10.1038/s41586-023-06323-x

81 Boninsegna, L. et al. Integrative genome modeling platform reveals essentiality of rare contact events in 3D genome organizations. Nat Methods 19, 938–949 (2022). 10.1038/s41592-022-01527-x

82 Su, J. H., Zheng, P., Kinrot, S. S., Bintu, B. & Zhuang, X. Genome-Scale Imaging of the 3D Organization and Transcriptional Activity of Chromatin. Cell 182, 1641–1659 e1626 (2020). 10.1016/j.cell.2020.07.032

83 Yildirim, A. et al. Evaluating the role of the nuclear microenvironment in gene function by population-based modeling. Nat Struct Mol Biol 30, 1193–1206 (2023). 10.1038/s41594-023-01036-1

84 Zhu, Q. et al. The transcription factor Pou3f1 promotes neural fate commitment via activation of neural lineage genes and inhibition of external signaling pathways. Elife 3 (2014). 10.7554/eLife.02224

85 Suzuki, N., Rohdewohld, H., Neuman, T., Gruss, P. & Scholer, H. R. Oct-6: a POU transcription factor expressed in embryonal stem cells and in the developing brain. EMBO J 9, 3723–3732 (1990). 10.1002/j.1460-2075.1990.tb07585.x

86 Murphy-Ullrich, J. E. Thrombospondin 1 and Its Diverse Roles as a Regulator of Extracellular Matrix in Fibrotic Disease. J Histochem Cytochem 67, 683–699 (2019). 10.1369/0022155419851103

87 Sid, B. et al. Thrombospondin 1: a multifunctional protein implicated in the regulation of tumor growth. Crit Rev Oncol Hematol 49, 245–258 (2004). 10.1016/j.critrevonc.2003.09.009

88 Alexander, K. A. et al. p53 mediates target gene association with nuclear speckles for amplified RNA expression. Mol Cell 81, 1666–1681 e1666 (2021). 10.1016/j.molcel.2021.03.006

89 Li, J. et al. Single-gene imaging links genome topology, promoter-enhancer communication and transcription control. Nat Struct Mol Biol 27, 1032–1040 (2020). 10.1038/s41594-020-0493-6

90 Chen, H. et al. Dynamic interplay between enhancer-promoter topology and gene activity. Nat Genet 50, 1296–1303 (2018). 10.1038/s41588-018-0175-z

91 Barbieri, M. et al. Complexity of chromatin folding is captured by the strings and binders switch model. Proc Natl Acad Sci U S A 109, 16173–16178 (2012). 10.1073/pnas.1204799109

92 Conte, M. et al. Polymer physics indicates chromatin folding variability across single-cells results from state degeneracy in phase separation. Nat Commun 11, 3289 (2020). 10.1038/s41467-020-17141-4

93 Nicodemi, M. & Prisco, A. Thermodynamic pathways to genome spatial organization in the cell nucleus. Biophys J 96, 2168–2177 (2009). 10.1016/j.bpj.2008.12.3919

94 Esposito, A. et al. Polymer physics reveals a combinatorial code linking 3D chromatin architecture to 1D chromatin states. Cell Rep 38, 110601 (2022). 10.1016/j.celrep.2022.110601

95 Conte, M. et al. Loop-extrusion and polymer phase-separation can co-exist at the single-molecule level to shape chromatin folding. Nat Commun 13, 4070 (2022). 10.1038/s41467-022-31856-6

96 Zhang, R., Zhou, T. & Ma, J. Multiscale and integrative single-cell Hi-C analysis with Higashi. Nat Biotechnol 40, 254–261 (2022). 10.1038/s41587-021-01034-y

97 Zhang, R., Zhou, T. & Ma, J. Author Correction: Multiscale and integrative single-cell Hi-C analysis with Higashi. Nat Biotechnol 40, 432 (2022). 10.1038/s41587-022-01263-9

98 Bintu, B. et al. Super-resolution chromatin tracing reveals domains and cooperative interactions in single cells. Science 362 (2018). 10.1126/science.aau1783

99 Yu, M. et al. SnapHiC: a computational pipeline to identify chromatin loops from single-cell Hi-C data. Nat Methods 18, 1056–1059 (2021). 10.1038/s41592-021-01231-2

100 Hounkpe, B. W., Chenou, F., de Lima, F. & De Paula, E. V. HRT Atlas v1.0 database: redefining human and mouse housekeeping genes and candidate reference transcripts by mining massive RNA-seq datasets. Nucleic Acids Res 49, D947–D955 (2021). 10.1093/nar/gkaa609

101 Bergman, D. T. et al. Compatibility rules of human enhancer and promoter sequences. Nature 607, 176–184 (2022). 10.1038/s41586-022-04877-w

102 Leemans, C. et al. Promoter-Intrinsic and Local Chromatin Features Determine Gene Repression in LADs. Cell 177, 852–864 e814 (2019). 10.1016/j.cell.2019.03.009

103 Madsen-Osterbye, J., Abdelhalim, M., Pickering, S. H. & Collas, P. Gene Regulatory Interactions at Lamina-Associated Domains. Genes (Basel) 14 (2023). 10.3390/genes14020334

104 Emerson, D. J. et al. Cohesin-mediated loop anchors confine the locations of human replication origins. Nature 606, 812–819 (2022). 10.1038/s41586-022-04803-0

105 Wang, W. et al. Genome-wide mapping of human DNA replication by optical replication mapping supports a stochastic model of eukaryotic replication. Mol Cell 81, 2975–2988 e2976 (2021). 10.1016/j.molcel.2021.05.024

106 Sima, J. et al. Identifying cis Elements for Spatiotemporal Control of Mammalian DNA Replication. Cell 176, 816–830 e818 (2019). 10.1016/j.cell.2018.11.036

107 Fudenberg, G., Kelley, D. R. & Pollard, K. S. Predicting 3D genome folding from DNA sequence with Akita. Nat Methods 17, 1111–1117 (2020). 10.1038/s41592-020-0958-x

108 Schwessinger, R. et al. DeepC: predicting 3D genome folding using megabase-scale transfer learning. Nat Methods 17, 1118–1124 (2020). 10.1038/s41592-020-0960-3

109 Zhou, J. Sequence-based modeling of three-dimensional genome architecture from kilobase to chromosome scale. Nat Genet 54, 725–734 (2022). 10.1038/s41588-022-01065-4

110 Zhang, R., Wang, Y., Yang, Y., Zhang, Y. & Ma, J. Predicting CTCF-mediated chromatin loops using CTCF-MP. Bioinformatics 34, i133–i141 (2018). 10.1093/bioinformatics/bty248

111 Feng, F., Yao, Y., Wang, X. Q. D., Zhang, X. & Liu, J. Connecting high-resolution 3D chromatin organization with epigenomics. Nat Commun 13, 2054 (2022). 10.1038/s41467-022-29695-6

112 Yang, R. et al. Epiphany: predicting Hi-C contact maps from 1D epigenomic signals. Genome Biol 24, 134 (2023). 10.1186/s13059-023-02934-9

113 Tan, J. et al. Cell-type-specific prediction of 3D chromatin organization enables high-throughput in silico genetic screening. Nat Biotechnol 41, 1140–1150 (2023). 10.1038/s41587-022-01612-8

114 Bianco, S. et al. Polymer physics predicts the effects of structural variants on chromatin architecture. Nat Genet 50, 662–667 (2018). 10.1038/s41588-018-0098-8

115 Avsec, Z. et al. Base-resolution models of transcription-factor binding reveal soft motif syntax. Nat Genet 53, 354–366 (2021). 10.1038/s41588-021-00782-6

116 Gunsalus, L. M., Keiser, M. J. & Pollard, K. S. In silico discovery of repetitive elements as key sequence determinants of 3D genome folding. Cell Genom 3, 100410 (2023). 10.1016/j.xgen.2023.100410

117 Gjoni, K. & Pollard, K. S. SuPreMo: a computational tool for streamlining in silico perturbation using sequence-based predictive models. Bioinformatics 40 (2024). 10.1093/bioinformatics/btae340

118 Hnisz, D. et al. Activation of proto-oncogenes by disruption of chromosome neighborhoods. Science 351, 1454–1458 (2016). 10.1126/science.aad9024

119 Ferraris, L. et al. Combinatorial binding of transcription factors in the pluripotency control regions of the genome. Genome Res 21, 1055–1064 (2011). 10.1101/gr.115824.110

120 Zhang, S. & Cui, W. Sox2, a key factor in the regulation of pluripotency and neural differentiation. World J Stem Cells 6, 305–311 (2014). 10.4252/wjsc.v6.i3.305

121 Vierbuchen, T. et al. AP-1 Transcription Factors and the BAF Complex Mediate Signal-Dependent Enhancer Selection. Mol Cell 68, 1067–1082 e1012 (2017). 10.1016/j.molcel.2017.11.026

