## Supplementary Methods for "An integrated view of the structure and function of the human 4D nucleome"

Supplemental Figure 1

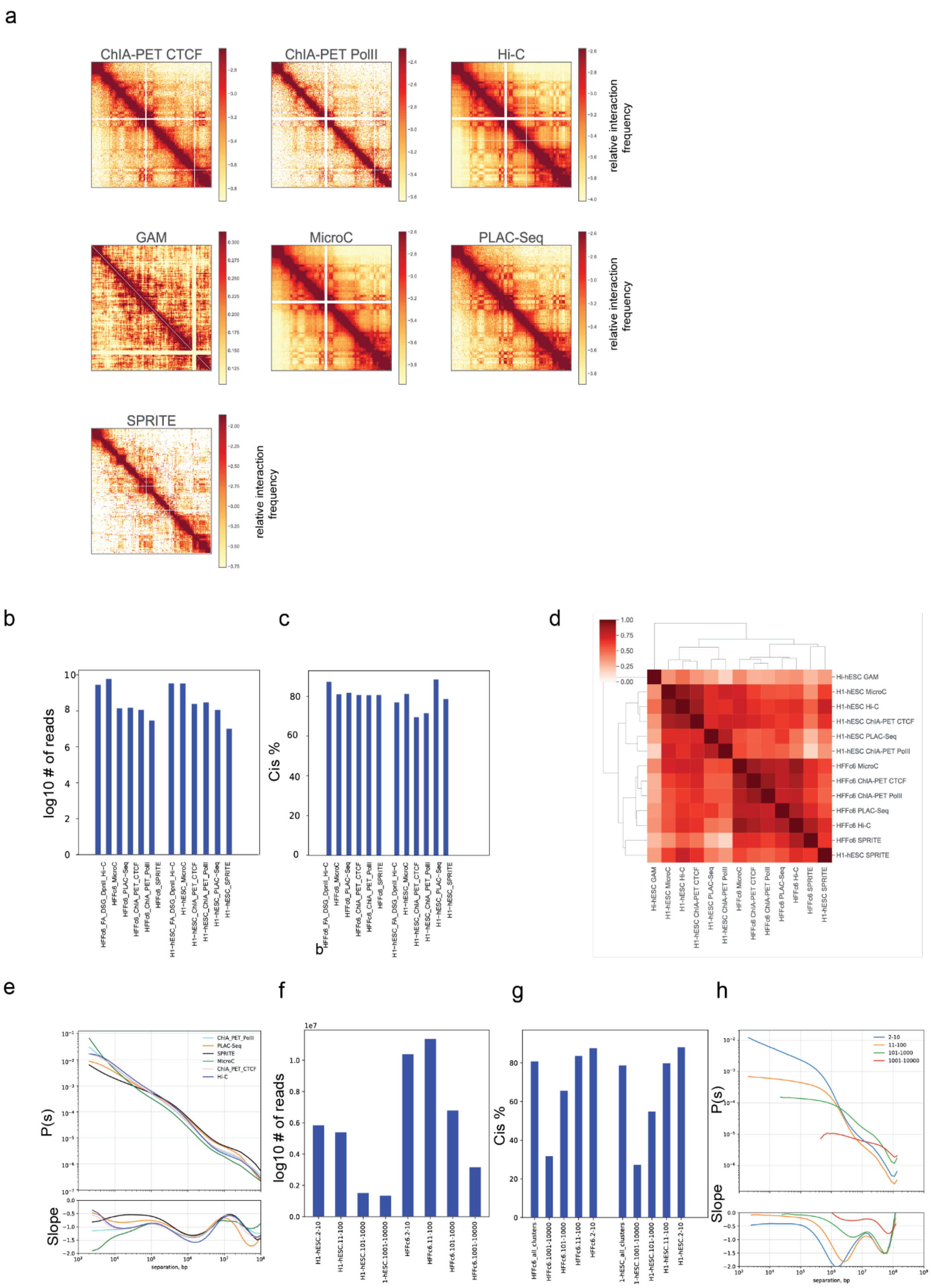

### Legend Supplemental Figure 1

**Quantitative metrics of data obtained with different chromatin interaction mapping methods**

**a**. Chromatin interaction maps obtained with different chromatin interaction mapping methods for H1-hESC cells. The region shown is Chromosome 19, 30,000,000-58,700,000. All data sets are shown at 100 kb bin size.

**b.** The number of reads obtained for Hi-C, Micro-C, ChIA PET, PLAC Seq, SPRITE datasets

**c.** The percentage of *cis* contacts for data obtained with each method indicated.

**d.** HiCRep [61] correlations of insulation profiles obtained with Hi-C, Micro-C, ChIA PET, PLAC Seq, GAM, and SPRITE

**e.** *P*(*s*) plot showing distance dependent contact probability of interactions detected with all protocols applied to HFFc6 cells (top). Derivative of the *P*(*s*) plots shown in panel d (bottom).

**f.** The number of fragments in each SPRITE cluster. Cluster sizes are 2-10, 11-100, 101-1000, 1001-10000 fragments.

**g.** The percentage of *cis* contacts in SPRITE clusters of indicated cluster size, for HFFc6 and H1-hESC cells.

**h.** *P*(*s*) plot showing distance dependent contact probability of interactions detected with different SPRITE cluster sizes for HFFc6 cells (top). Derivative of the *P*(*s*) plots shown in the bottom panel.

Supplemental Figure 2

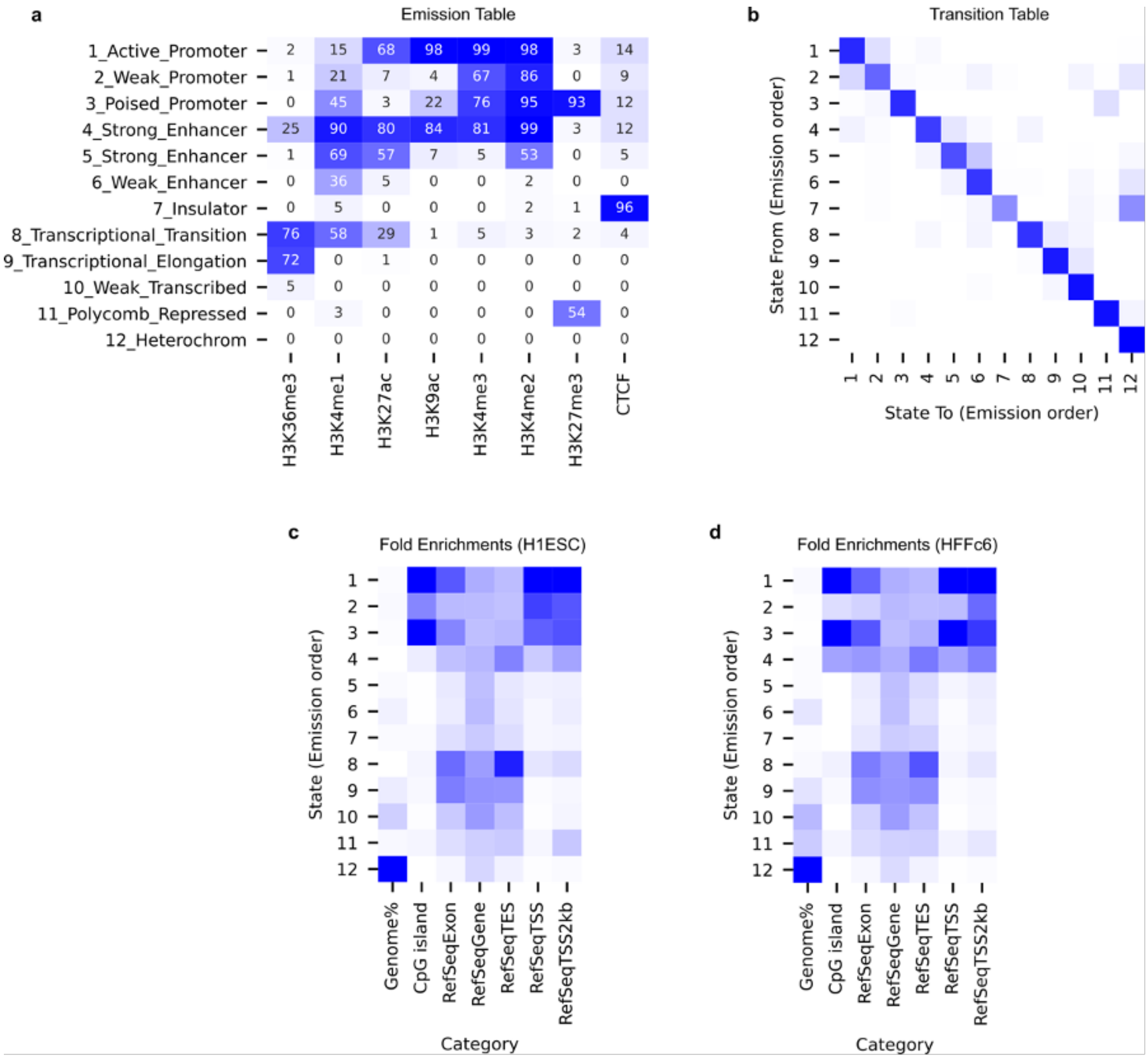

### Legend Supplemental Figure 2

**Consensus ChromHMM segmentation in H1-hESC and HFFc6.**

**a.** Heatmap for the emission parameters of the model.

**b**. Heatmap for the transition parameters of the model.

**c-d**. Relative percentage of the genome represented by each chromatin state and relative fold enrichment for several types of genomic annotations. TSS, transcription start site; TES, transcription end site.

Supplemental Figure 3

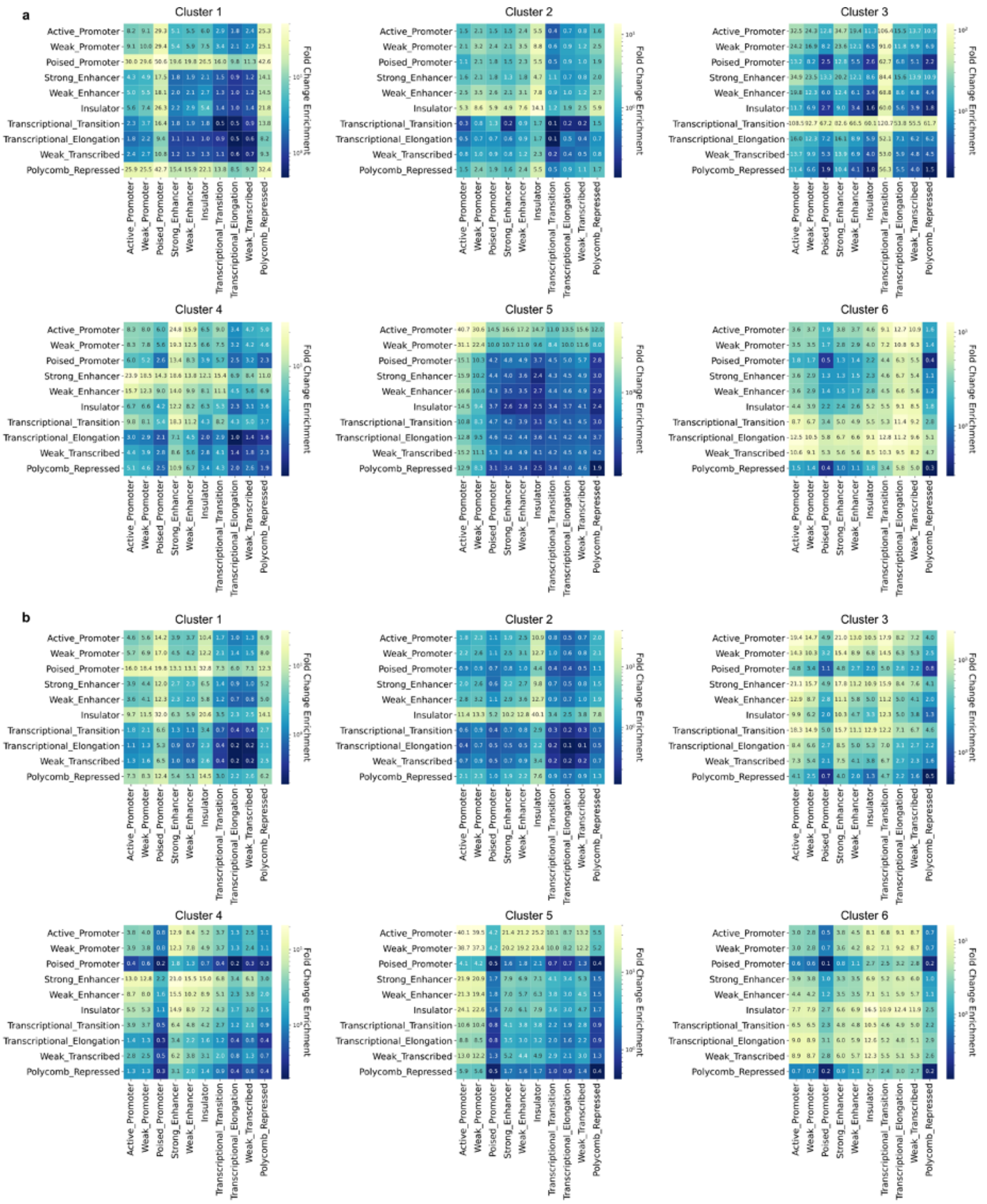

### Legend Supplemental Figure 3

**Fold-enrichment scores of ChromHMM states for different loop clusters revealed by the UMAP in H1-hESC (a) and HFFc6 (b).**

Supplemental Figure 4

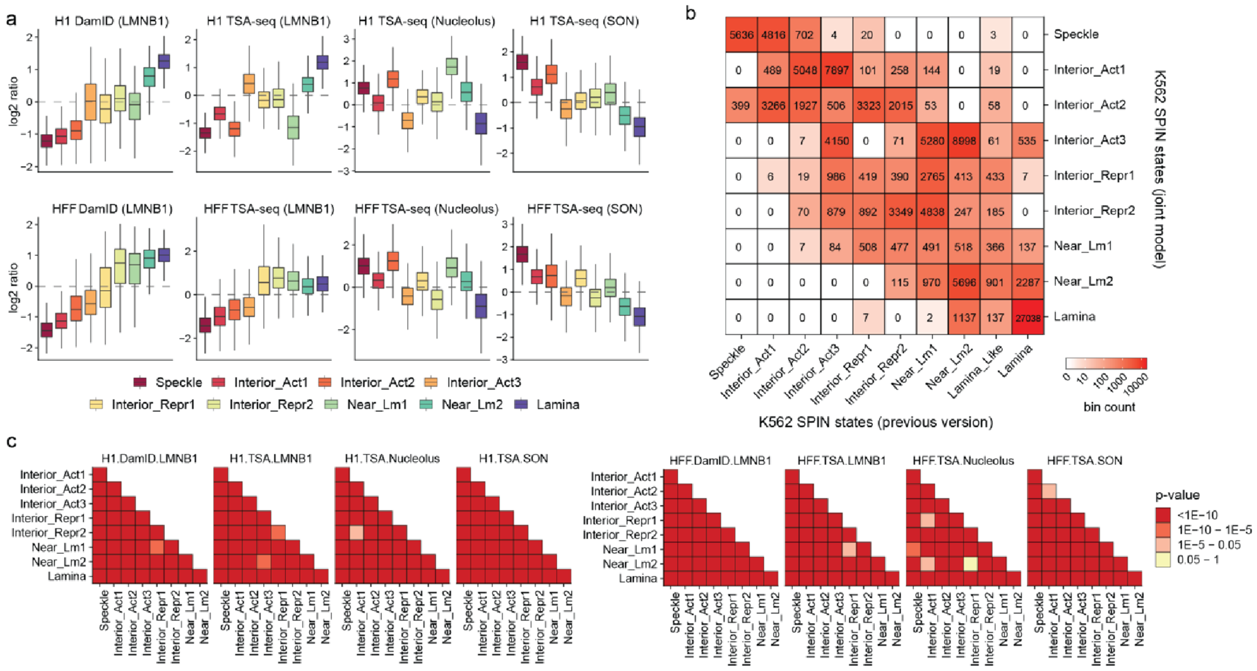

### Legend Supplemental Figure 4

**a.** Box plots show the distribution of normalized TSA-seq and DamID scores on distinct SPIN states in H1 and HFFc6 cell lines.

**b.** A confusion matrix shows the comparison of SPIN states in K562 between the previous version and the new result based on the joint modeling across four cell lines. Note that the new result is based on a new nucleolus TSA-seq data. The numbers in the heatmap indicate the number of 25kb bins.

**c.** The differences of the distributions of normalized TSA-seq and DamID between any two pairs of SPIN states are tested by the Wilcoxon rank sum test. Colors of heatmap indicate the p-value under the null hypothesis that two distributions are derived from the same population.

### Supplemental Figure 5

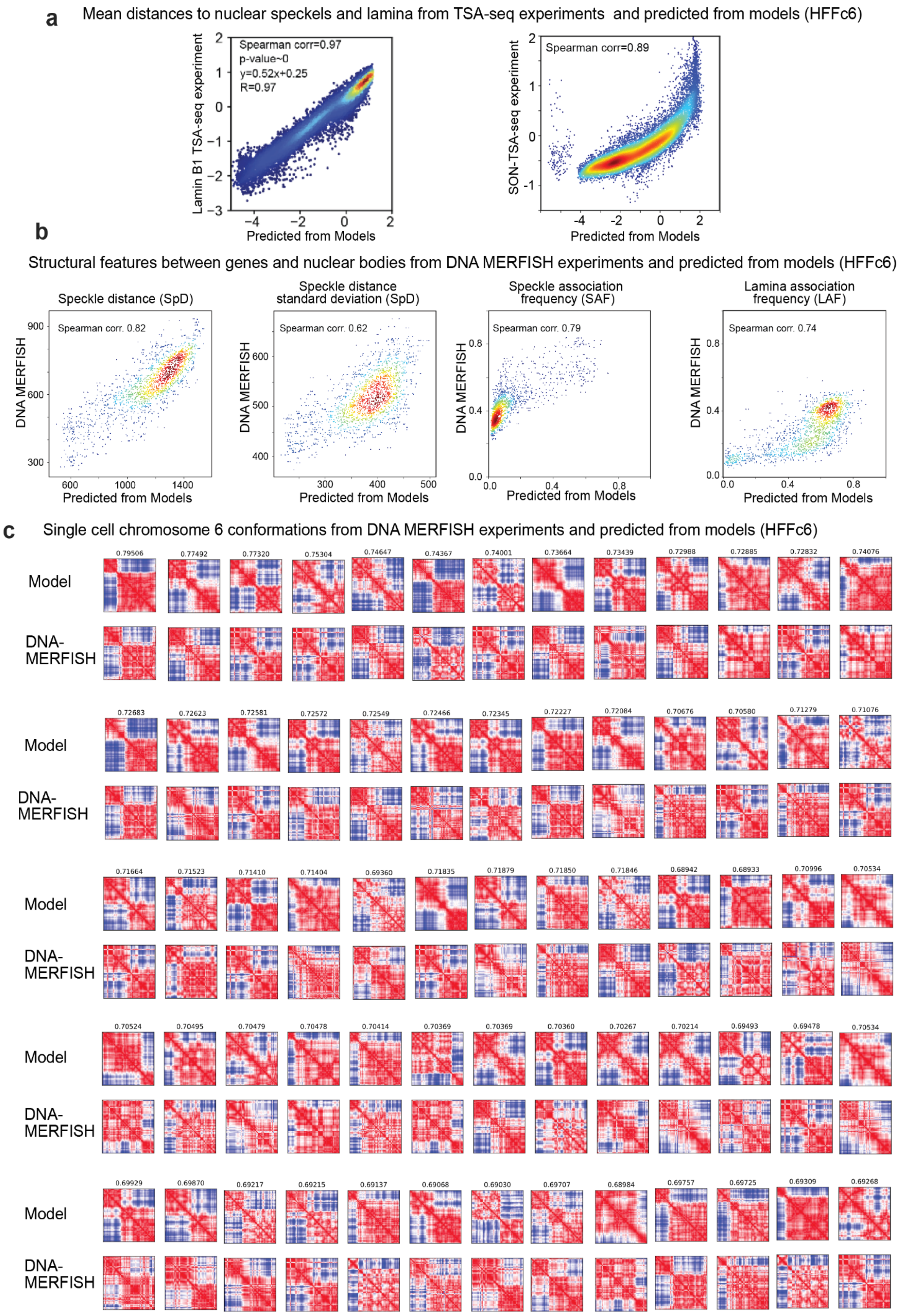

### Legend Supplementary Figure 5

**Assessment of 3D genome structure models against independent experimental data.** Assessment of genome structure models with independent experimental data.

**a.** Genome-wide correlation of TSA-seq data from experiment and predicted from our models (Left, Lamin B1 TSA-seq [91]. Right, SON TSA-seq [56].

**b.** Genome-wide correlation of structure features between genome structures from DNA-MERFISH chromosome tracing experiments [82] and predictions from our models. (From left to right, mean speckle distance (SpD), standard deviation of mean speckle distance in structure population, Speckle association frequency (SAF), Lamina association frequency (LAF)). **c.** Comparison of single cell distance matrices of chromosome 6 from simulated models (Top panel) and DNA-MERFISH imaging data [82] (Bottom panel). Numbers above the distance matrices represent Pearson correlations between distance matrices from models and experiment.

### Supplemental Figure 6

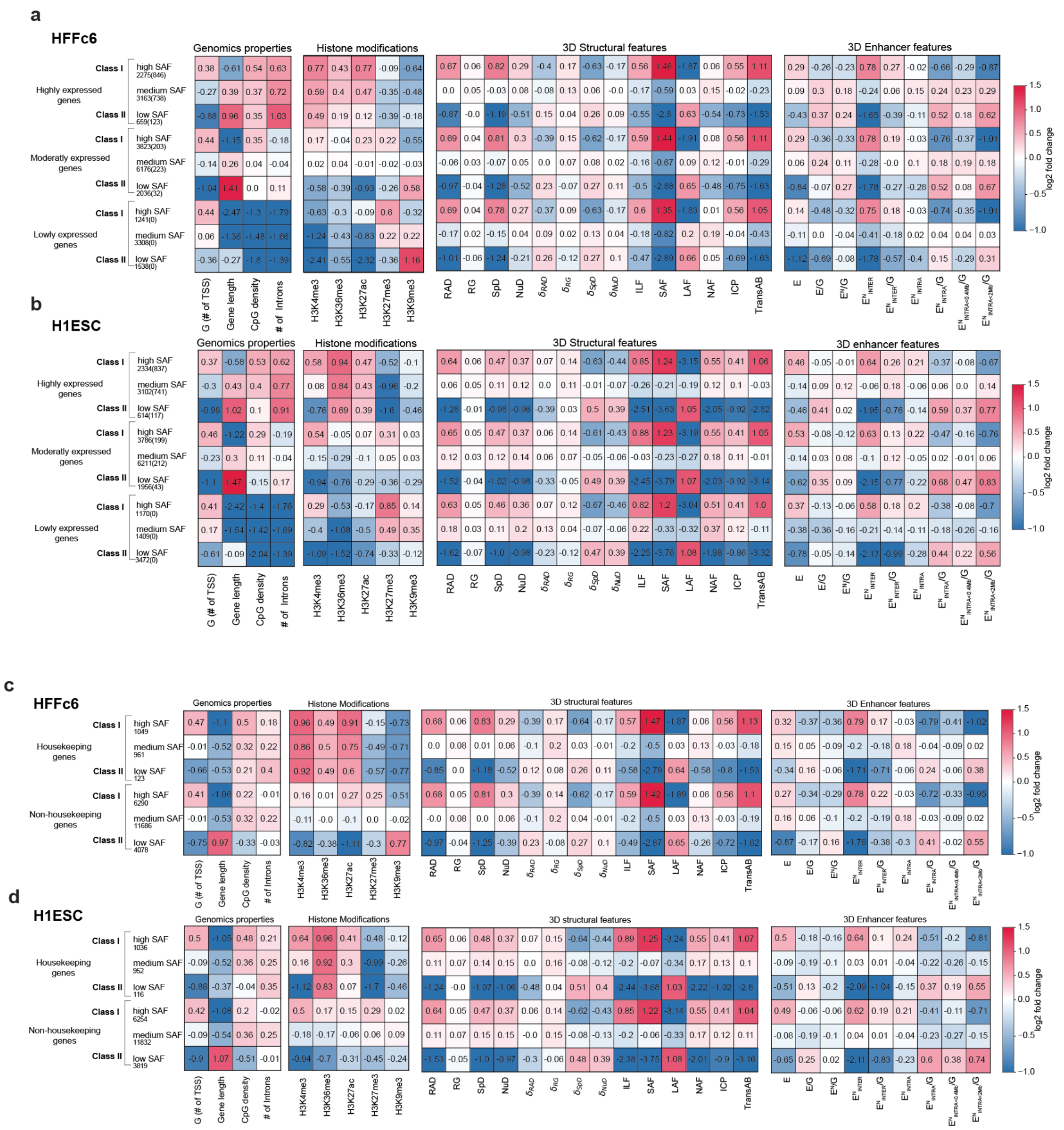

### Legend Supplementary Figure 6

**Enrichment plots for structural and functional features of genes.**

**a.** Log-fold enrichment for genomic properties (calculated within a 200 kb region), histone modifications (within +/- 10 kb of TSS), 3D structure features, and 3D spatial enhancer densities at each TSS (Methods) for genes stratified by their speckle association frequencies in HFFc6.

**b.** Same as in K but for H1-hESC cells.

**c.** Same analysis for HFFc6 cells as in A, but now genes are stratified into housekeeping genes and non-housekeeping genes.

**d.** Same as in panel **c**, but for H1-hESC cells.

Supplemental Figure 7

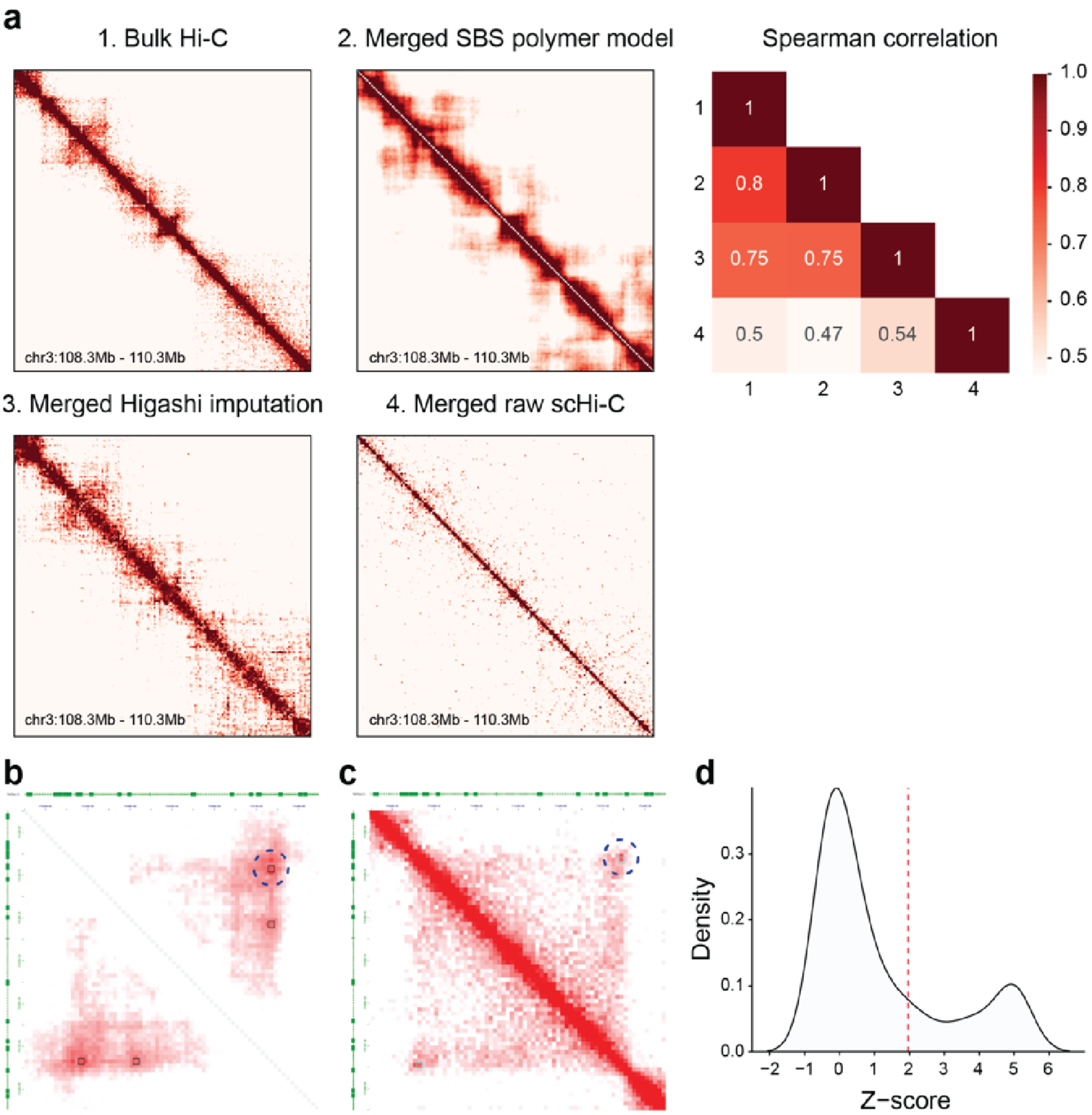

### Legend Supplemental Figure 7

**WTC11 scHi-C analysis compared with bulk Hi-C.**

**a.** Comparison of contact frequency map near the DPPA locus between bulk Hi-C merge SBS polymer model, merged Higashi imputed contact frequency map and raw contact frequency map without imputation. The heatmap on the top-right shows the Spearman correlation coefficients between these contact frequency maps.

**b.** The heatmap shows the aggregated contact map from single-cell Hi-C data at the gene *RABGAP1L* locus (for cells with z-score>1.96). The circle with dashed line indicates the 450 kb loop identified by SnapHi-C.

**c.** KR-normalized Hi-C contact frequency from WTC11 bulk Hi-C data.

**d.** The distribution of Z-score of 188 WTC11 single cells at the chromatin loop at the gene RABGAP1L locus. The red vertical dash line represents Z-score = 1.96.

Supplemental Figure 8

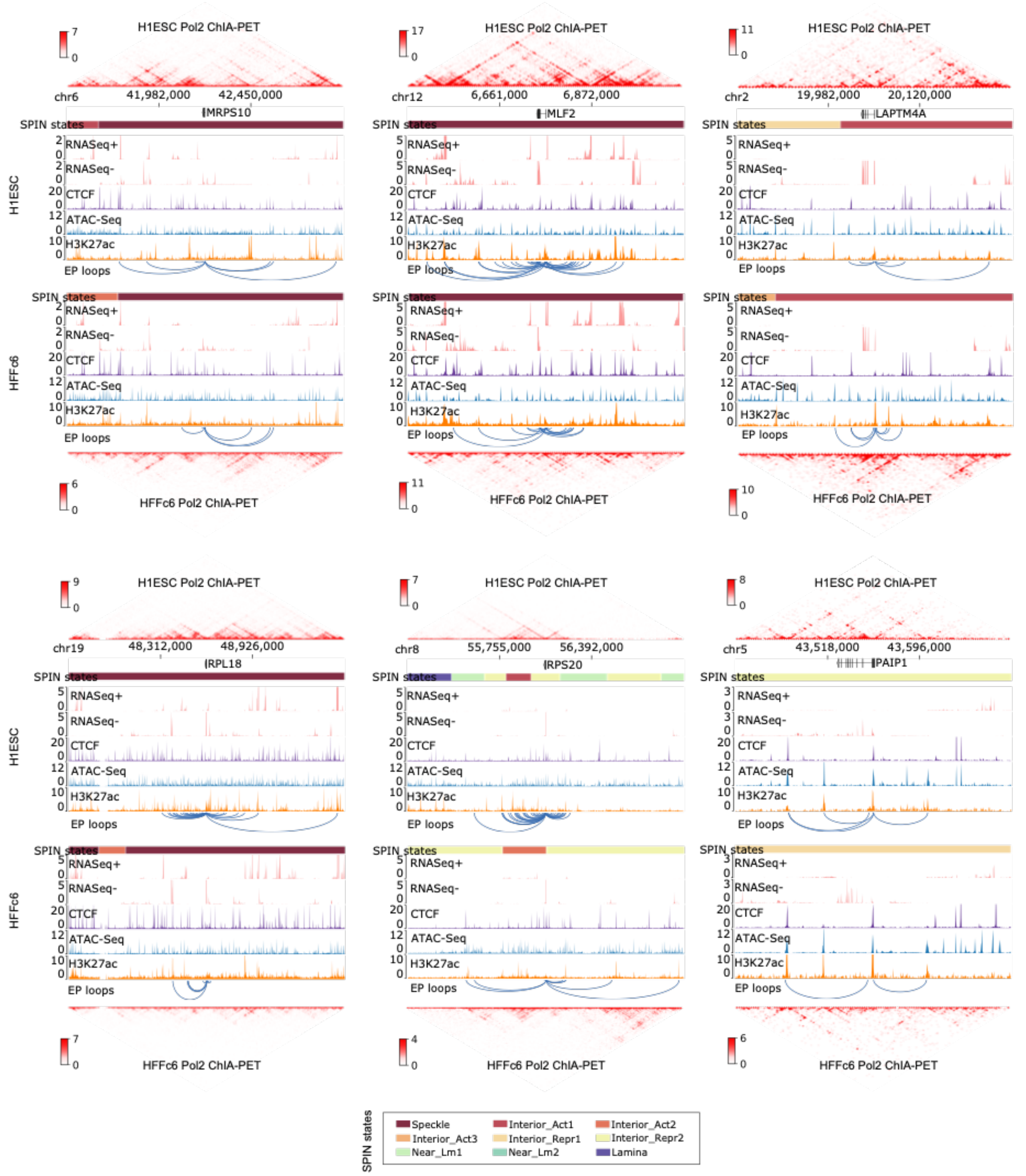

### Legend Supplemental Figure 8

**Example genome-browser views showing house-keeping genes usually interact with different sets of distal enhancers in different cell lines.** Blue arcs represent chromatin loops linking the indicated house-keeping genes with distal enhancers.

Supplemental Figure 9

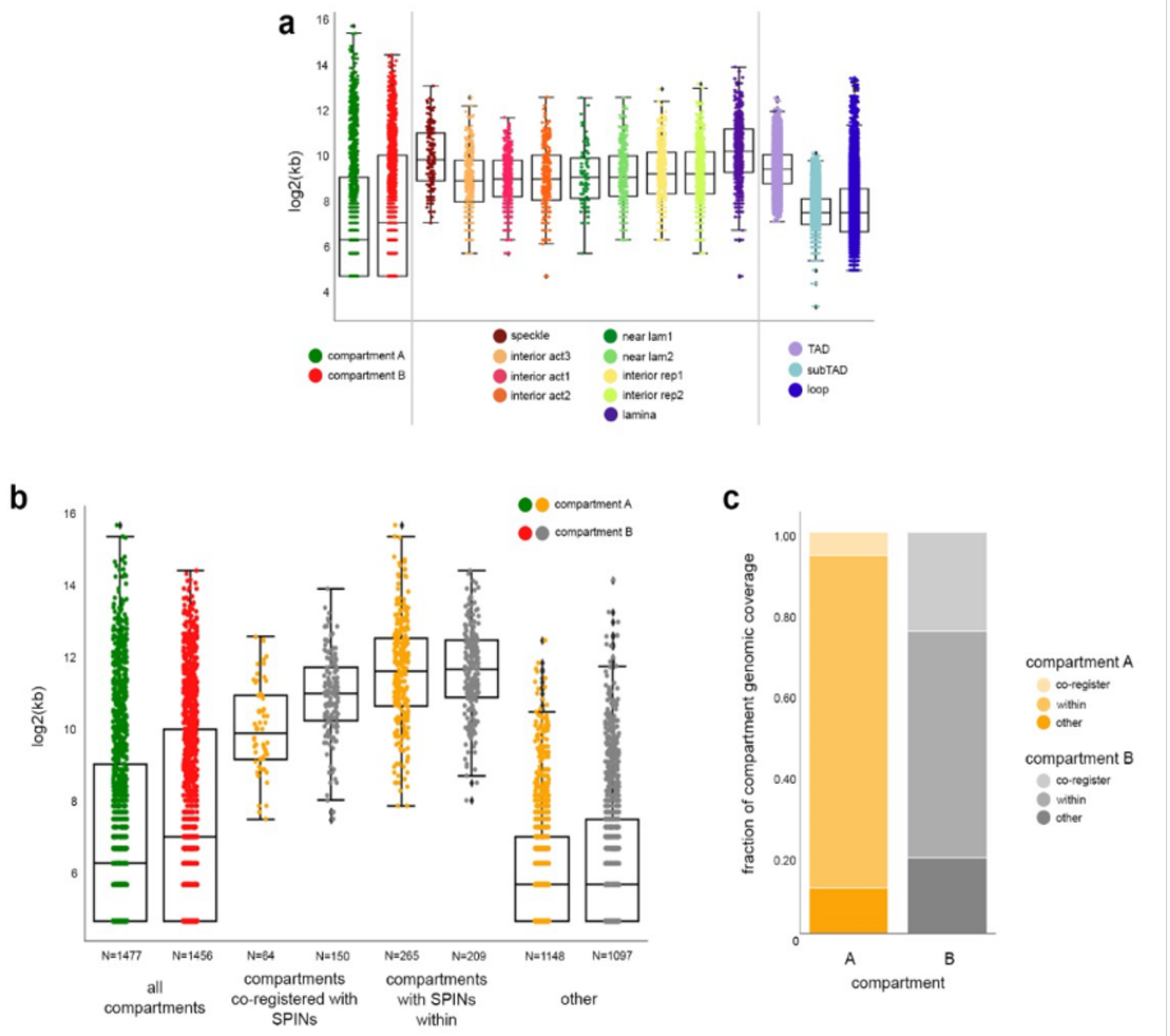

### Legend Supplemental Figure 9

**Compartments and SPIN comparison.**

**a.** Size comparison of genome folding features in H1-hESCs. All regions plotted by size of their genomic coordinates, box plot shows 25th, 50th, and 75th quartiles, whiskers show minimum and maximum values with outliers annotated.

**b.** Genomic size of A/B compartments stratified by SPIN alignment in H1-hESCs. Total compartment A (green; N=1477) and total compartment B (red ; N=1456) were stratified by those that co-register with a SPIN, fully encompass or contain a SPIN within, or other, including nested within a SPIN and no SPIN intersection. **c.** Genomic coverage of A/B compartments stratified by SPIN alignment.

**Links to code**

Methods benchmarking:

https://github.com/dekkerlab/Flagship_paper.git

GAM analysis

https://github.com/pombo-lab/WinickNg_Kukalev_Harabula_Nature_2021

Loop calling, and transcription-loop analysis:

https://github.com/XiaoTaoWang/4DN-joint-analysis)

TADs, SPIN, Compartment, and replication analysis:

https://bitbucket.org/creminslab/cremins_lab_4dn_phase1_jawg_code__2023/

Predicting Hi-C data from sequence:

https://github.com/shuzhenkuang/Contact_map_prediction_visualization.

Replication timing analysis

https://github.com/ClaireMarchal/flagship_paper_scripts

**Links to data**

All data described in this work are publicly available at the 4DN Data Coordination and Integration Center (https://data.4dnucleome.org/), or through other publicly accessible portals listed below for each relevant dataset.

### Methods and data links

#### 1. Methods and data for benchmarking section

### Datasets

| Cell Type | Experimental Type | Data Type | Data Source/Download Link | Note |
|---|---|---|---|---|
| H1-ESC | Hi-C | Contact matrix | 4DN (4DNFI82R42AD) | |
| | Micro-C | Contact matrix | 4DN (4DNFI9GMP2J8) | |
| | CTCF ChIA-PET | Contact matrix | 4DN (4DNFINMHXGVQ) | |
| | RNAPII ChIA-PET | Contact matrix | 4DN (4DNFIO8YJ5JA) | |
| | H3K4me3 PLAC-Seq | Contact matrix | 4DN (4DNFICOGAKW2) | |
| | DNA SPRITE | Contact matrix | 4DN (4DNFITX6WCRT) | |
| | | Clusters | 4DN (4DNFIV3PDS5F, 4DNFIIY1TXUZ) | |
| | GAM | Contact matrix at 25kb | 4DNESVAMUDHA | |
| | CTCF ChIP-Seq | Signal (Bigwig) | https://epigenome.wustl.edu/epimap/data/observed/FINAL_CTCF_BSS00478.sub_VS_FINAL_WCE_BSS00478.pval.signal.bedgraph.gz.bigWig | ChromHMM Input |
| | H3K27ac ChIP-Seq | Signal (Bigwig) | https://epigenome.wustl.edu/epimap/data/observed/FINAL_H3K27ac_BSS00478.sub_VS_FINAL_WCE_BSS00478.pval.signal.bedgraph.gz.bigWig | |
| | H3K27me3 ChIP-Seq | Signal (Bigwig) | https://epigenome.wustl.edu/epimap/data/observed/FINAL_H3K27me3_BSS00478.sub_VS_FINAL_WCE_BSS00478.pval.signal.bedgraph.gz.bigWig | |
| | H3K36me3 ChIP-Seq | Signal (Bigwig) | https://epigenome.wustl.edu/epimap/data/observed/FINAL_H3K36me3_BSS00478.sub_VS_FINAL_WCE_BSS00478.pval.signal.bedgraph.gz.bigWig | |

| | | | |
|---|---|---|---|
| H3K4me1 ChIP-Seq | Signal (Bigwig) | https://epigenome.wustl.edu/epimap/data/observed/FINAL_H3K4me1_BSS00478.sub_VS_FINAL_WCE_BSS00478.pval.signal.bedgraph.gz.bigWig | |
| H3K9ac ChIP-Seq | Signal (Bigwig) | https://epigenome.wustl.edu/epimap/data/observed/FINAL_H3K9ac_BSS00478.sub_VS_FINAL_WCE_BSS00478.pval.signal.bedgraph.gz.bigWig | |
| H3K4me3 ChIP-Seq | Signal (Bigwig) | https://epigenome.wustl.edu/epimap/data/observed/FINAL_H3K4me3_BSS00478.sub_VS_FINAL_WCE_BSS00478.pval.signal.bedgraph.gz.bigWig | |
| H3K4me2 ChIP-Seq | Signal (Bigwig) | https://epigenome.wustl.edu/epimap/data/observed/FINAL_H3K4me2_BSS00478.sub_VS_FINAL_WCE_BSS00478.pval.signal.bedgraph.gz.bigWig | |
| RNA-Seq | Signal (Bigwig) | ENCODE (ENCFF501KFP, ENCFF563OKS) | Visualization |
| CTCF ChIP-Seq | Signal (Bigwig) | ENCODE (ENCFF332TNJ) | |
| H3K27ac ChIP-Seq | Signal (Bigwig) | ENCODE (ENCFF103PND) | |
| H3K27me3 ChIP-Seq | Signal (Bigwig) | ENCODE (ENCFF502GXT) | |
| EZH2 ChIP-Seq | Signal (Bigwig) | ENCODE (ENCFF109KCQ) | |
| POLR2A ChIP-Seq | Signal (Bigwig) | ENCODE (ENCFF942TZX) | |
| CHD1 ChIP-Seq | Signal (Bigwig) | ENCODE (ENCFF597VKW) | |
| KDM4A ChIP-Seq | Signal (Bigwig) | ENCODE (ENCFF269CHA) | |
| PHF8 ChIP-Seq | Signal (Bigwig) | ENCODE (ENCFF059EBB) | |
| TAF1 ChIP-Seq | Signal (Bigwig) | ENCODE (ENCFF837BSZ) | |
| RAD21 ChIP-Seq | Signal (Bigwig) | ENCODE (ENCFF056GWP) | |
| KDM1A ChIP-Seq | Peaks | ENCODE (ENCFF759CSN) | Enrichment analysis |
| CBX8 ChIP-Seq | Peaks | ENCODE (ENCFF483UZG) | |
| EZH2 ChIP-Seq | Peaks | ENCODE (ENCFF414CAB) | |
| KDM4A ChIP-Seq | Peaks | ENCODE (ENCFF021QGZ) | |
| SP4 ChIP-Seq | Peaks | ENCODE (ENCFF257FUV) | |
| MAX ChIP-Seq | Peaks | ENCODE (ENCFF994IHO) | |
| POU5F1 ChIP-Seq | Peaks | ENCODE (ENCFF383EYO) | |
| CHD1 ChIP-Seq | Peaks | ENCODE (ENCFF731EYW) | |
| ZNF143 ChIP-Seq | Peaks | ENCODE (ENCFF235ROG) | |
| TAF1 ChIP-Seq | Peaks | ENCODE (ENCFF886BPR) | |
| TCF12 ChIP-Seq | Peaks | ENCODE (ENCFF959HJP) | |

| | | |
|---|---|---|
| CHD7 ChIP-Seq | Peaks | ENCODE (ENCFF338IDU) |
| CEBPB ChIP-Seq | Peaks | ENCODE (ENCFF823KCM) |
| BRCA1 ChIP-Seq | Peaks | ENCODE (ENCFF721TNS) |
| MXI1 ChIP-Seq | Peaks | ENCODE (ENCFF727RNJ) |
| CTBP2 ChIP-Seq | Peaks | ENCODE (ENCFF501VJW) |
| SIRT6 ChIP-Seq | Peaks | ENCODE (ENCFF219XEX) |
| SUZ12 ChIP-Seq | Peaks | ENCODE (ENCFF225AMM) |
| EP300 ChIP-Seq | Peaks | ENCODE (ENCFF244VKF) |
| ZNF274 ChIP-Seq | Peaks | ENCODE (ENCFF718OGI) |
| HDAC6 ChIP-Seq | Peaks | ENCODE (ENCFF802HUU) |
| GABPA ChIP-Seq | Peaks | ENCODE (ENCFF308NZT) |
| POLR2A ChIP-Seq | Peaks | ENCODE (ENCFF322DAE) |
| GTF2F1 ChIP-Seq | Peaks | ENCODE (ENCFF138MYA) |
| MYC ChIP-Seq | Peaks | ENCODE (ENCFF049SMR) |
| NRF1 ChIP-Seq | Peaks | ENCODE (ENCFF414RES) |
| REST ChIP-Seq | Peaks | ENCODE (ENCFF738LQB) |
| JUN ChIP-Seq | Peaks | ENCODE (ENCFF821GUI) |
| HDAC2 ChIP-Seq | Peaks | ENCODE (ENCFF923TXH) |
| RXRA ChIP-Seq | Peaks | ENCODE (ENCFF745EBL) |
| CTCF ChIP-Seq | Peaks | ENCODE (ENCFF023LAA) |
| YY1 ChIP-Seq | Peaks | ENCODE (ENCFF376FVJ) |
| ATF3 ChIP-Seq | Peaks | ENCODE (ENCFF440FTA) |
| USF2 ChIP-Seq | Peaks | ENCODE (ENCFF346KIW) |
| BCL11A ChIP-Seq | Peaks | ENCODE (ENCFF847HXU) |
| JUND ChIP-Seq | Peaks | ENCODE (ENCFF287KKY) |
| USF1 ChIP-Seq | Peaks | ENCODE (ENCFF978MNS) |
| RNF2 ChIP-Seq | Peaks | ENCODE (ENCFF241UKW) |
| SP2 ChIP-Seq | Peaks | ENCODE (ENCFF309QRC) |
| SIX5 ChIP-Seq | Peaks | ENCODE (ENCFF384KWP) |
| CBX5 ChIP-Seq | Peaks | ENCODE (ENCFF218OXB) |
| ATF2 ChIP-Seq | Peaks | ENCODE (ENCFF352KLD) |
| RFX5 ChIP-Seq | Peaks | ENCODE (ENCFF142NQQ) |
| FOSL1 ChIP-Seq | Peaks | ENCODE (ENCFF428RHR) |
| NANOG ChIP-Seq | Peaks | ENCODE (ENCFF435DTC) |
| POLR2AphosphoS5 ChIP-Seq | Peaks | ENCODE (ENCFF872MKT) |
| BACH1 ChIP-Seq | Peaks | ENCODE (ENCFF749UPP) |
| E2F6 ChIP-Seq | Peaks | ENCODE (ENCFF174AVU) |
| CHD2 ChIP-Seq | Peaks | ENCODE (ENCFF726GBF) |
| RAD21 ChIP-Seq | Peaks | ENCODE (ENCFF883FUW) |
| SRF ChIP-Seq | Peaks | ENCODE (ENCFF648QJE) |
| EGR1 ChIP-Seq | Peaks | ENCODE (ENCFF100KKH) |
| TEAD4 ChIP-Seq | Peaks | ENCODE (ENCFF885PQR) |
| TBP ChIP-Seq | Peaks | ENCODE (ENCFF817TGF) |

| | | | | |
|---|---|---|---|---|
| | CREB1 ChIP-Seq | Peaks | ENCODE (ENCFF978KGB) | |
| | MAFK ChIP-Seq | Peaks | ENCODE (ENCFF710YJK) | |
| | RBBP5 ChIP-Seq | Peaks | ENCODE (ENCFF610PCN) | |
| | SIN3A ChIP-Seq | Peaks | ENCODE (ENCFF432EYM) | |
| | SP1 ChIP-Seq | Peaks | ENCODE (ENCFF284JVS) | |
| | SAP30 ChIP-Seq | Peaks | ENCODE (ENCFF043HXQ) | |
| | KDM5A ChIP-Seq | Peaks | ENCODE (ENCFF471OCM) | |
| | ASH2L ChIP-Seq | Peaks | ENCODE (ENCFF693FGQ) | |
| HFFc6 | Hi-C | Contact matrix | 4DN (4DNFIAVXXO55) | |
| | Micro-C | Contact matrix | 4DN (4DNFI9FVHJZQ) | |
| | CTCF ChIA-PET | Contact matrix | 4DN (4DNFIG2ILS39) | |
| | RNAPII ChIA-PET | Contact matrix | 4DN (4DNFIIASUSSX) | |
| | H3K4me3 PLAC-Seq | Contact matrix | 4DN (4DNFI9REIU8H) | |
| | DNA SPRITE | Contact matrix | 4DN (4DNFIP9HGF9M) | |
| | | Clusters | 4DN (4DNFIRXON7Z2) | |
| | CTCF ChIP-Seq | Signal (Bigwig) | https://epigenome.wustl.edu/epimap/data/observed/FINAL_CTCF_BSS00353.sub_VS_FINAL_WCE_BSS00353.pval.signal.bedgraph.gz.bigWig | ChromHMM Input |
| | H3K27ac ChIP-Seq | Signal (Bigwig) | https://epigenome.wustl.edu/epimap/data/observed/FINAL_H3K27ac_BSS00353.sub_VS_FINAL_WCE_BSS00353.pval.signal.bedgraph.gz.bigWig | |
| | H3K27me3 ChIP-Seq | Signal (Bigwig) | https://epigenome.wustl.edu/epimap/data/observed/FINAL_H3K27me3_BSS00353.sub_VS_FINAL_WCE_BSS00353.pval.signal.bedgraph.gz.bigWig | |
| | H3K36me3 ChIP-Seq | Signal (Bigwig) | https://epigenome.wustl.edu/epimap/data/observed/FINAL_H3K36me3_BSS00353.sub_VS_FINAL_WCE_BSS00353.pval.signal.bedgraph.gz.bigWig | |
| | H3K4me1 ChIP-Seq | Signal (Bigwig) | https://epigenome.wustl.edu/epimap/data/observed/FINAL_H3K4me1_BSS00353.sub_VS_FINAL_WCE_BSS00353.pval.signal.bedgraph.gz.bigWig | |
| | H3K9ac ChIP-Seq | Signal (Bigwig) | https://epigenome.wustl.edu/epimap/data/imputed/impute_BSS00353_H3K9ac.bigWig | |

| | | |
|---|---|---|
| H3K4me3 ChIP-Seq | Signal (Bigwig) | https://epigenome.wustl.edu/epimap/data/observed/FINAL_H3K4me3_BSS00353.sub_VS_FINAL_WCE_BSS00353.pval.signal.bedgraph.gz.bigWig |
| H3K4me2 ChIP-Seq | Signal (Bigwig) | https://epigenome.wustl.edu/epimap/data/imputed/impute_BSS00353_H3K4me2.bigWig |

**Genome Architecture Mapping methods and data processing**

**Preparation of cryosections.** H1-hESCs were fixed and processed for cryosectioning as described previously {Branco, 2006 #572}. Briefly, H1-hESCs were grown to 70% confluency, media was removed, and cells were fixed in 4% and 8% paraformaldehyde in 250 mM HEPES-NaOH (pH 7.6; 10 min and 2 h, respectively), gently scrapped, and softly pelleted, before embedding (>2h) in saturated 2.1 M sucrose in PBS and frozen in liquid nitrogen on copper sample holders. Frozen samples were stored in liquid nitrogen. Ultrathin cryosections were cut using a Leica ultracryomicrotome (UltraCut EM UC7, Leica Microsystems) at approximately 220 nm thickness, captured on sucrose-PBS drops and transferred to 4 μm PEN steel frame slide for laser microdissection (Leica Microsystems, Cat# 11600289). Sucrose embedding medium was removed by washing with 0.2 μm filtered molecular biology grade PBS (3 × 5 min each) and filtered ultra-pure water (5 min). For laser microdissection, cryosections on PEN membranes were washed, permeabilized and incubated (2 h, room temperature) in blocking solution (1% BSA (w/v), 5% FBS (w/v, GibcoTM Cat#10270), 0.05% Triton X-100 (v/v) in PBS). After incubation (overnight, 4°C) with primary anti-pan-histone (1:50) antibody (Merck, Cat#MAB3422) in blocking solution, the cryosections were washed (3-5x; 30 min) in 0.025% Triton X-100 in PBS (v/v) and immunolabeled (1 h, room temperature) with secondary antibodies in blocking solution, followed by 3 (15 min) washes in PBS.

**Isolation of nuclear profiles.** Nuclear staining was visualized using a Leica laser microdissection microscope (Leica Microsystems, LMD7000) using a 63x dry objective. Individual nuclear profiles (NPs) were laser micro-dissected from the PEN membrane, and collected into PCR adhesive caps (AdhesiveStrip 8C opaque; Carl Zeiss Microscopy #415190-9161-000). GAM data was collected in multiplexGAM mode, where three NPs are collected into each adhesive cap. The presence of NPs in each lid was confirmed with a 5x objective using a 420-480 nm emission filter. Control lids not containing nuclear profiles (water controls) were included for each dataset collection to keep track of contamination and noise amplification of whole genome amplification and library reactions. Collected nuclear profiles were kept at -20oC until whole-genome amplification.

**Whole-genome amplification (WGA).** Whole Genome Amplification (WGA) was performed as described previously {Winick-Ng, 2021 #1952} with minor modifications. Briefly, DNA was extracted from NPs at $60^0$C in the lysis buffer (20 mM Tris-HCl pH 8.0, 1.4 mM EDTA, 560 mM guanidinium-HCl, 3.5% Tween-20, 0.35% Triton X-100) containing 0.75 units/ml Qiagen protease (Qiagen, 19155). After 24h of DNA extraction, the protease was heat inactivated at $75^0$C for 30 min and the extracted DNA was amplified via two rounds of PCR. At-first quasi-linear amplification was performed with random hexamer GAT-7N primers with an adaptor sequence. The lysis buffer containing the extracted genomic DNA was mixed with 2x DeepVent mix buffer (2x Thermo polymerase buffer (10x), 400 µm dNTPs, 4 mM $MgSO_4$ in ultrapure DNase free water), 0.5 µM GAT-7N primers (5′- GTG AGT GAT GGT TGA GGT AGT GTG GAG NNN NNN N) and 2 units/µl DeepVent® (exo-) DNA polymerase (New England Biolabs, M0259L) and incubated for 11 cycles in the BioRad thermocycler. The second exponential PCR amplification was performed in presence of 1x DeepVent mix, 10 mM dNTPs, 0.4 µM GAM-COM primers (5′-GTG AGT GAT GGT TGA GGT AGT GTG GAG) and 2 units/µl DeepVent (exo-) DNA polymerase in the programmable thermal cycler for 26 cycles. WGA was performed in 96-well plates using Microlab STARLine liquid handling workstation (Hamilton).

**Preparation of GAM libraries for high-throughput sequencing**. Following WGA, the samples were purified using the SPRI magnetic beads (1.7x ratio of beads per sample volume). The DNA concentration of each purified sample was measured using the Quant-iT PicoGreen dsDNA assay kit (Invitrogen Cat#P7589). Sequencing libraries were then made using the in-house tagmentation based protocol. Following library preparation, DNA concentration for each sample was measured using the Quant-iT PicoGreen dsDNA assay, and equal amounts of DNA from each sample was pooled together. The final pool of libraries was analyzed using DNA High Sensitivity on-chip electrophoresis on an Agilent 2100 Bioanalyzer and sequenced on Illumina NextSeq 500 machine.

**GAM data sequence alignment.** Sequenced reads from each GAM library were mapped to the human genome assembly GRCh38 (December 2013, hg38) with bowtie2 (v.2.3.4.3) using default settings. All non-uniquely mapped reads, reads with mapping quality <20 and PCR duplicates were excluded from further analyses.

**GAM data window calling and sample QC.** Positive genomic windows present within ultrathin nuclear slices were identified for each GAM library as previously described {Winick-Ng, 2021 #1952}. In brief, the genome was split into equal-sized windows, and the number of nucleotides sequenced in each bin was calculated for each GAM sample with bedtools. Next, we determined the percentage of orphan windows (that is, positive windows that were flanked by two adjacent negative windows) for every percentile of the nucleotide coverage distribution. The number of nucleotides that corresponds to the percentile with the lowest percentage of orphan windows in each sample was used as an optimal coverage threshold for window identification in each sample. Windows were called positive if the number of nucleotides sequenced in each bin was greater than the determined optimal threshold.

The sample quality was assessed by the percentage of orphan windows in each sample, total genomic coverage in percent of positive windows, the number of uniquely mapped reads to the mouse genome and the correlations from cross-well contamination for every sample. Each sample was considered to be of good quality if it had </=40% orphan windows, </=60% of total genome coverage, >50,000 uniquely mapped reads and a cross-well contamination score determined per collection plate of <0.4 (Jaccard index).

**GAM data curation.** To exclude genomic windows which were under- or oversampled in the GAM collection, we computed a GAM specific parameter, the window detection efficiency (WDF, {Beagrie, 2017 #1351} as previously described {Irastorza-Azcarate, 2024 #1953}. To detect genomic bins with outlying detection frequency, a smoothing algorithm was applied to the WDF values per chromosome in stretches of eleven equally sized genomic windows. Next, normalized delta (ND) was calculated for each window, according to: ND = (raw_Signal - smoothed_Signal)/smoothed Signal. If the ND was larger than a fold change of 5, the window was removed from the final dataset. Next, the four adjacent windows (2 upstream and 2 downstream) were also removed, to ensure good quality of sampling in the final GAM data used for further analyses.

Genomic bins with an average mappability score below 0.2 were also removed. Genome mappability for the hg38 human genome assembly was computed using GEM-Tools suite {Marco-Sola, 2012 #1954} setting read length to 75 nucleotides. The mean mappability score was computed for each genomic bin with bigWigAverageOverBed utility from Encode.

**GAM data normalization.** GAM contact matrices for all pairs of windows genome-wide were generated as previously described, to produce pair-wise co-segregation maps and pointwise mutual information (NPMI) maps which consider window detectability {Winick-Ng, 2021

#1952}. For visualization of the contact matrices, scale bars are adjusted in each genomic region displayed to a range between 0 and the 99th percentile of NPMI values in the region.

**GAM Insulation scores.** The insulation scores were calculated from NPMI-normalized pairwise chromatin contact matrices, as previously described {Crane, 2015 #1172} with minor modifications adjusted for GAM input data by keeping both positive and negative values in the matrix {Winick-Ng, 2021 #1952}.

**Identification of compartments A and B in GAM data.** Compartments were calculated using co-segregation matrices, as previously described {Irastorza-Azcarate, 2024 #1953}. In brief, each chromosome was represented as a matrix of observed interactions O(i,j) between locus i and locus j. The expected interactions E(i,j) matrix was calculated, where each pair of genomic windows is the mean number of contacts with the same distance between i and j. A matrix of observed over expected values O/E(i,j) was produced by dividing O by E. A correlation matrix C(i,j) was calculated between column i and column j of the O/E matrix. PCA was performed for the first three components on matrix C before extracting the component with the highest correlation to GC content. Loci with eigenvector values with the same sign were designated as A compartments, while those with the opposite sign were identified as B compartments. For chromosomes 3 and 22, we manually picked PC1 and PC2, respectively, as the PC that correlated most with GC content did not display a typical AB compartmentalization pattern and good correlation with transcription levels.

**4DN accession number:** 4DNESVAMUDHA (replicates id 4DNBS46FF9D9, 4DNBSB45FJC1)
**Software:** bowtie2 (v.2.3.4.3), samtools (v.1.14), bedtools (v.2.30), fastq_screen (v.0.14.0), GEM-tools(v.1.315)

**Data processing - Hi-C, Micro-C, ChIA-PET, PLAC-Seq and SPRITE**
Cooler files for Hi-C, Micro-C, ChIA-PET, PLAC-Seq and SPRITE were downloaded from DCIC Data Portal. The link for Hi-C, MIcro-C and ChIA-PET files can be found here:
https://data.4dnucleome.org/browse/?type=ExperimentSetReplicate&experimentset_type=replicate&experiments_in_set.biosample.biosource.organism.name=human&experiments_in_set.biosample.biosource_summary=HFFc6+%28Tier+1%29&experiments_in_set.biosample.biosource_summary=H1-hESC+%28Tier+1%29&experiments_in_set.experiment_categorizer.combined=Target%3A+RNA+Pol+II&experiments_in_set.experiment_categorizer.combined=Target%3A+CTCF+protein&experiments_in_set.experiment_categorizer.combined=Enzyme%3A+DpnII&experimen

ts_in_set.experiment_categorizer.combined=Enzyme%3A+MNase&experiments_in_set.experiment_type.display_title=in+situ+Hi-C&experiments_in_set.experiment_type.display_title=DNA+SPRITE&experiments_in_set.experiment_type.display_title=Micro-C&experiments_in_set.experiment_type.display_title=PLAC-seq&experiments_in_set.experiment_type.display_title=in+situ+ChIA-PET. Link for PLAC-Seq and SPRITE files can be found here: https://data.4dnucleome.org/browse/?type=ExperimentSetReplicate&experimentset_type=replicate&experiments_in_set.biosample.biosource.organism.name=human&experiments_in_set.biosample.biosource_summary=H1-hESC+%28Tier+1%29&experiments_in_set.biosample.biosource_summary=HFFc6+%28Tier+1%29&experiments_in_set.experiment_type.display_title=DNA+SPRITE&experiments_in_set.experiment_type.display_title=PLAC-seq. Contact matrices were normalized using the iterative correction procedure from Imakaev et al. 2012 {Imakaev, 2012 #1013}.
Interaction heatmaps were created using Python. The color map is “YlOrRd” and the color scales are created taking the 10$^{th}$ and 90$^{th}$ percentile of the interaction frequencies of individual datasets.
GAM H1-hESC was also downloaded form 4DN Data Portal and can be found here: https://data.4dnucleome.org/search/?q=GAM+H1-hESC&type=Item . No additional processing was applied to GAM data.

**Hicrep correlations**

HiCRep is used to do distance corrected correlations of the multiple methods {Yardimci, 2019 #1896}. Correlation is calculated in two steps. First, interaction maps are stratified by genomic distances and the correlation coefficients are calculated for each distance separately. Second, the reproducibility is determined by a novel stratum-adjusted correlation coefficient statistic (SCC) by aggregating stratum-specific correlation coefficients using a weighted average. Chromosome specific correlation was performed for pairwise protocols and averaged the correlations across those chromosomes. Averaged pairwise correlations of chr1-22 and chr X between Hi-C, Micro-C, ChIA-PET, PLAC-Seq and SPRITE. Averaged correlation of chr 1-22 for GAM and other methods. 50kb binned interaction matrices are used to calculate Hicrep correlations.

**Compartment Analysis**

Compartments were assessed for Hi-C, Micro-C, ChIA-PET, PLAC-Seq and SPRITE using eigenvector decomposition on observed-over-expected contact maps at 100kb resolution using the cooltools package derived scripts {Open2C, 2024 #1955}. Eigenvector that has the strongest correlation with gene density is selected, then A and B compartments were assigned based on the gene density profiles such that A compartment has high gene density and B compartment has low gene density profile. A and B compartment assignments of GAM were provided by the data producers.

Spearman correlation was used to correlate the eigenvectors of different experiments performed with various protocols and cell states. Saddle plots were generated as follows (described in {Nora, 2017 #1381}: the interaction matrix of an experiment was sorted based on the eigenvector values from lowest to highest (B to A). Sorted maps were then normalized for their expected interaction frequencies; the upper left corner of the interaction matrix represents the strongest B-B interactions, lower right represents strongest A-A interactions, upper right and lower left are B-A and A-B respectively. To quantify saddle plots we took the strongest 20% of BB and strongest 20% of AA interactions and normalized them by the sum of AB and BA (top(AA)/(AB+BA) and top(BB)/(BA+AB)). Saddle quantifications were used to create the scatter plots. The list of parameters that are used for the saddle plot are; --strength, --vmin 0.5, --vmax 2 , --regions hg38_chromsizes.bed, --qrange 0.02 0.98, --contact-type cis.

**Preferential Interactions**

Bigwig or bedgraph files for LMNB1 DamID, TSA-Seq and Repli-Seq were downloaded from DCIC Data Portal. The link for those files can be found here : https://data.4dnucleome.org/browse/?dataset_label=E%2FL+repliseq+on+H1-hESC+cells+%282017-08-17%29&dataset_label=E%2FL+repliseq+on+HFFc6+cells+%282017-08-17%29&dataset_label=E%2FL+repliseq+on+HFFc6+cells+-+Gold+Standard++%282018-02-06%29&dataset_label=TSA-seq+MKI67IP+in+H1+cells&dataset_label=TSA-seq+MKI67IP+in+HFFc6+cells&dataset_label=TSA-seq+POL1RE+in+H1+cells&dataset_label=TSA-seq+POL1RE+in+HFFc6+cells&dataset_label=TSA-seq+v2+SON+in+H1&dataset_label=TSA-seq+v2+SON+in+HFFc6&dataset_label=DamID-seq+on+H1-hESC+cells+%282017-12-14%29&dataset_label=DamID-seq+on+HFFc6+cells+%282017-12-14%29&dataset_label=ChIA-PET+in+H1-hESC&dataset_label=ChIA-PET+in+HFFc6&experiments_in_set.biosample.biosource.organism.name=human&experiments_in_set.experiment_categorizer.combined=Target%3A+LMNB1+protein&experiments_in_set.experiment_categorizer.combined=Target%3A+CTCF+protein&experiments_in_set.ex

periment_categorizer.combined=Target%3A+NIFK+protein&experiments_in_set.experiment_categorizer.combined=Target%3A+POLR1E+protein&experiments_in_set.experiment_categorizer.combined=Target%3A+SON+protein&experiments_in_set.experiment_categorizer.combined=Fraction%3A+early+fraction+of+2+fractions&experiments_in_set.experiment_categorizer.combined=Fraction%3A+late+fraction+of+2+fractions&experiments_in_set.experiment_categorizer.combined=Target%3A+RNA+Pol+II&experiments_in_set.experiment_categorizer.combined=No+value&experimentset_type=replicate&type=ExperimentSetReplicate

Heatmaps that integrate 3D methods with genome activity plots were generated as follows: First, the data was binned into 50kb bins for aforementioned assays and sorted from the highest to the lowest value. Additional filters were applied for Early/Late replication ratio. For Early/Late replication timing data; removed bins with no values and the bins with value of 0. Also removed the outlier bins that have values > 98th quantile and kept the min value for the first bin as 0.

Second, the interaction matrices (Hi-C, Micro-C, ChIA-PET, PLAC-Seq, SPRITE and GAM) are sorted based on the 1D tracks generated from the aforementioned assays from the highest to the lowest.

Next, sorted matrices were then normalized for their expected interaction frequencies; the upper left corner of the interaction matrix represents the strongest signal for non-preferential interactions, lower right represents strongest preferential interactions. To quantify these plots we took the strongest 20% of the preferential interactions. Saddle plot parameters are listed below for this quantifications: --strength, --vmin 0.5, --vmax 2 , --regions hg38_chromsizes.bed, --qrange 0.02 0.98, --range min(Sorted 1D data) max(Sorted 1D data) --contact-type cis. For GAM, --strength, --vmin 0.1, --vmax 0.4 , --regions hg38_chromsizes.bed, --qrange 0.02 0.98, --range min(Sorted 1D data) max(Sorted 1D data) --contact-type cis.

**Insulation Score**

For Hi-C, Micro-C, ChIA-PET, PLAC-Seq and SPRITE we calculated diamond insulation scores using cooltools ( https://github.com/open2c/cooltools/blob/master/cooltools/cli/diamond_insulation.py ) as implemented from Crane et al {Crane, 2015 #1172}. We defined the insulation and boundary strengths of each 25 kb bin by detecting the local minima of 25 kb binned data with a 100kb window size. We used cooltools's *diamond-insulation* function with these parameters: " --ignore-diags 2. Insulation scores of GAM were provided by data producers. We separated weak and strong log2 insulation scores using the mean insulation score of all protocols (i.e.,: weak insulation scores < mean < strong  insulation scores). We piled up strong insulation scores to compare the average insulation score strengths of methods.

**Identification of chromatin loops in different platforms**

We employed different strategies for detecting chromatin loops in different platforms. For Hi-C and Micro-C, we combined results from HiCCUPS {Rao, 2014 #1176} and Peakachu {Salameh, 2020 #1897}. To identify chromatin loops using HiCCUPS, we ran "cooltools dots" (v0.5.1 {Open2C, 2024 #1955}) at 5kb and 10kb resolutions with default parameters. Peakachu is a machine-learning based framework that learns contact patterns of pre-defined chromatin loops from a genome-wide contact map and applies trained models to predict loops on other maps generated by the same/similar experimental protocol. Here, we first trained Peakachu models on GM12878 Hi-C data at 5 kb and 10 kb resolutions, using a high-confidence loop set detected by at least two platforms among Hi-C, CTCF ChIA-PET, Pol2 ChIA-PET, CTCF HiChIP, H3K27ac HiChIP, SMC1A HiChIP, H3K4me3 PLAC-Seq, and TrAC-loop. These models were then used to predict chromatin loops on Hi-C and Micro-C maps of H1-hESC and HFFc6 cell lines at corresponding resolutions. The probability cutoffs were manually adjusted to balance sensitivity and specificity based on visual inspection.

For ChIA-PET, we combined loop predictions from ChiaSig {Paulsen, 2014 #1956} and Peakachu. For each ChIA-PET dataset, we conducted multiple runs of ChiaSig with varying parameter settings, specifically adjusting the "-M", "-C", and "-c" parameters while keeping other parameters constant ("-m 8000 -S 4 -s 6 -A 0.01 -a 0.1 -n 1000"). The "-M" value was selected from 1000000, 2000000, and 4000000, while both the "-C" and "-c" values were set to either 2 or 3. Only chromatin loops consistently identified across all parameter settings were retained, while others were discarded. As ChiaSig heavily relies on one-dimensional (1D) peak annotation for loop detection, chromatin interactions outside peak regions are not identified as loops. To capture loops with similar contact patterns to those detected by ChiaSig but outside peak regions, we again utilized Peakachu to learn the patterns. For each ChIA-PET dataset, we trained 23 Peakachu models using interactions detected by ChiaSig, with each model trained on data from different combinations of 22 chromosomes. During prediction, loops on each chromosome were predicted using the model trained on the other 22 chromosomes. The probability cutoffs were determined to ensure that Peakachu-predicted loops covered 90% of ChiaSig-detected interactions. Training and prediction were conducted separately at 2 kb and 5 kb resolutions, and the final loop predictions for each

ChIA-PET dataset were obtained by combining ChiaSig-predicted interactions and Peakachu predictions.

For PLAC-Seq, we identified chromatin loops at 10 kb resolution using MAPS {Juric, 2019 #1957} with default parameters.

When calculating the union loops from different platforms, methods, and resolutions, two chromatin loops (i,j) and (i',j') were considered overlapped if and only if |i-i'|<min(0.2|i-j|,15 kb) and |j-j'|<min(0.2|i-j|,15 kb). If two loops are overlapped, only the one predicted at a higher resolution with a more precise location was retained.

**Consensus chromatin-state annotations for H1-hESC and HFFc6 cells**

We computed epigenomic annotations using ChromHMM (v1.23) on 14 observed and 2 imputed ChIP-Seq datasets for 8 marks (H3K36me3, H3K4me1, H3K27ac, H3K9ac, H3K3me3, H3K4me2, H3K27me3, and CTCF) in both H1-hESC and HFFc6 cells. All the ChIP-Seq datasets were obtained from the WashU Epigenome Browser (https://epigenome.wustl.edu/epimap/data/) in bigwig format, and the coordinates were transformed from hg19 to hg38 using CrossMap (v0.5.2, http://crossmap.sourceforge.net/). To prepare the data for ChromHMM, we divided the whole genome into 200 bp bins and calculated the average signals within each bin. For the observed data, values were binarized with a -log10P value cutoff of 2. For the imputed data (H3K9ac and H3K4me2 in HFFc6), we downloaded both the imputed and observed data in H1-hESC for the same marks. Then, for each mark, we set the binarization cutoff for the imputed data to match the quantile in the observed data corresponding to the -log10P>2 cutoff, enabling comparison with the observed data.

Finally, we ran the “ChromHMM LearnModel” command on the binarized data to segment both the H1-hESC and HFFc6 genomes into 12 states. The name of each state was manually annotated based on prior knowledge about each mark. The “12_Heterochrom” state was excluded from further analysis, as it did not contain signals of any marks (Supplemental Figure 2a).

**Enrichment analysis of chromatin states for chromatin loops and loop anchors**

To characterize the chromatin states of loop anchors detected by specific combinations of chromatin interaction methods, we calculated fold-enrichment scores by comparing the overlap with each ChromHMM state between the observed loop anchors and 100 randomly generated control sets. Specifically, for each chromatin state, we iterated through the loop anchor list and counted the number of anchors overlapping at least one region with that state. We then randomly shuffled the anchors in the genome to generate 1,000 control sets and repeated the same procedure for each control. For each control, we kept the size distribution and the number of random regions on each chromosome the same as the observed loop anchors, and the intervals of each region did not overlap with any gaps in the hg38 reference genome. Finally, the fold-enrichment score was calculated by dividing the number of anchors with a specific chromatin state by the average number of random loci with the same state.

We employed a similar method to characterize chromatin states for a specific cluster of chromatin loops. Briefly, for each pair of chromatin states, we iterated through the loop list and counted the number of loops with one anchor overlapping regions marked by one chromatin state and the other anchor overlapping regions marked by the other chromatin state. Again, we generated 1,000 random control sets for chromatin loops. Each random loop set maintained the same genomic distance distribution between loop anchors and the same number of random loops on each chromosome, ensuring that the interval between the two ends of each loop did not overlap any gaps in hg38. Finally, the fold-enrichment score was calculated by dividing the number of loops between a specific pair of chromatin states by the average number of random loops between the same states.

**Enrichment analysis of transcription factors for different loop clusters**

To explore whether different loop clusters exhibit differential binding of various transcription factors (TFs) at their anchors, we downloaded the ENCODE ChIP-Seq peak files for 62 TFs in H1-hESCs. A fold enrichment score was computed for each TF at loop anchors using a procedure analogous to the one described above. Briefly, we first identified non-redundant loop anchors from each loop cluster in H1-hESCs. For each TF, we iterated through this anchor list and counted the number of anchors overlapping at least one ChIP-Seq peak. Subsequently, we generated 1,000 random control sets by shuffling the loops and repeated the same procedure for each control set. The fold-enrichment score was then calculated by dividing the number of anchors containing ChIP-Seq peaks by the average number of random loci containing ChIP-Seq peaks for the same TF.

**UMAP projection of chromatin loops**

To construct an input feature matrix for projecting chromatin loops, we calculated the proportion of each ChromHMM state at interacting loop anchors. This resulted in a feature matrix Mij of size 124,061 for H1ESC and a feature matrix Nij of size 115,850 for HFFc6. Each row in the matrix represents one chromatin loop, with the first 11 columns representing features of one anchor and the next 11 columns representing features of the other anchor. Subsequently, we standardized (z-score normalized) both Mij and Nij to ensure comparability between different features.

Next, for each row of the normalized matrices Mij and Nij, we swapped the order of the two anchors to ensure that the highest value was always observed in the first 11 columns. Following this, we concatenated Mij and Nij row-wise to create a new combined matrix. This new matrix served as input for training the UMAP (https://github.com/lmcinnes/umap) projection function with the parameters "n_neighbors=40, min_dist=0, n_components=2, metric='correlation'".

The same UMAP projection function was utilized to project chromatin loops from different cell lines and different platforms.

**Calculation of the average contact strength for different loop clusters**

To calculate the average contact strength for each loop cluster across different experimental platforms, we utilized distance-normalized (observed/expected) contact frequencies. Specifically, for Hi-C, Micro-C, and DNA SPRITE datasets, we computed this value using interaction frequencies normalized by matrix balancing or iterative correction and eigenvector decomposition (ICE) at the 5kb resolution. In contrast, for ChIA-PET and PLAC-Seq datasets, we calculated the value using raw interaction frequencies at the same 5kb resolution. For GAM, we used the NPMI-normalized co-segregation frequencies at the 25 kb resolution.

### 2. Methods for relating chromatin loops to gene expression.

#### Datasets

| Description | Cell Type | Data Source/Download Link |
|---|---|---|
| Loop predictions combined from multiple 4DN experimental assays | H1-ESC | https://www.jianguoyun.com/p/DQkAPQkQh9qdDBik0cYFIAA |
| | HFFc6 | https://www.jianguoyun.com/p/Dd-fjT8Qh9qdDBim0cYFIAA |
| ATAC-Seq peaks | H1-ESC | 4DN (4DNFI247OOFU) |
| | HFFc6 | 4DN (4DNFIWQJFZHS) |

| | | |
|---|---|---|
| House-keeping genes in human | - | HRT Atlas v1.0 (https://housekeeping.unicamp.br/) |
| H3K27ac ChIP-Seq signals | H1-ESC | ENCODE (ENCFF986PCY) |
| | HFFc6 | 4DN (4DNFINRI6WOL) |
| ATAC-Seq signals | H1-ESC | 4DN (4DNFICPNO4M5) |
| | HFFc6 | 4DN (4DNFIZ9191QU) |
| CTCF ChIP-Seq signals | H1-ESC | ENCODE (ENCFF332TNJ) |
| | HFFc6 | ENCODE (ENCFF406SZM) |
| LMNB1 DamID-Seq signals | H1-ESC | 4DN (4DNFI6BH48Y3) |
| | HFFc6 | 4DN (4DNFI7724Y7Q) |
| RNA-Seq for 116 tissues / cell types | ovary | ENCODE (ENCFF095OFV, ENCFF940WXY, ENCFF857MSA) |
| | right renal cortex interstitium | ENCODE (ENCFF320UVF, ENCFF821JAE, ENCFF981FYW) |
| | hindlimb muscle | ENCODE (ENCFF680ZPA) |
| | heart right ventricle | ENCODE (ENCFF823DWN, ENCFF102BTQ) |
| | A549 | ENCODE (ENCFF627QMV, ENCFF369ZNM) |
| | chorionic villus | ENCODE (ENCFF274JIK,ENCFF101AAR, ENCFF529CAT,E NCFF834AEP) |
| | Panc1 | ENCODE (ENCFF890DEQ, ENCFF248YCR) |
| | luminal epithelial cell of mammary gland | ENCODE (ENCFF047QOH) |
| | SK-N-SH | ENCODE (ENCFF389TFR, ENCFF161JEA, ENCFF390KQP, ENCFF067ZMG) |
| | natural killer cell | ENCODE (ENCFF036GDL) |
| | testis | ENCODE (ENCFF850LMK, ENCFF845QSA) |
| | RWPE-1 | ENCODE (SRR8446409, SRR8446410, SRR8446411) |
| | aorta | ENCODE (ENCFF277LBD, ENCFF914RDG) |

| | |
|---|---|
| germinal matrix | ENCODE (ENCFF951YSP) |
| brain | ENCODE (ENCFF784ZTQ) |
| right lung | ENCODE (ENCFF035KGA, ENCFF535JUK, ENCFF015DMG, ENCFF083YGV, ENCFF487DRK) |
| endodermal cell | ENCODE (ENCFF563QGW, ENCFF237ZQX) |
| lung | ENCODE (ENCFF947WLV, ENCFF051UVH, ENCFF014OUE) |
| liver | ENCODE (ENCFF239EUU, ENCFF908GIP, ENCFF592KZK, ENCFF203UGC) |
| myoepithelial cell of mammary gland | ENCODE (ENCFF674EKN) |
| GM23248 | ENCODE (ENCFF341SCS, ENCFF775DYT) |
| endocrine pancreas | ENCODE (ENCFF174RSS, ENCFF982TBJ) |
| foreskin melanocyte | ENCODE (ENCFF441UUO, ENCFF724NAG) |
| adrenal gland | ENCODE (ENCFF217TKV, ENCFF802ADF, ENCFF555RGZ, ENCFF467PRR, ENCFF866LBS, ENCFF918DYI, ENCFF739OIE, ENCFF908UKE) |
| neural cell | ENCODE (ENCFF813LWT, ENCFF081JBX) |
| fibroblast of lung | ENCODE (ENCFF227FMH, ENCFF983VCS) |
| MCF10A | ENCODE (SRR5364109, SRR5364108, SRR5364107, SRR5364106) |
| U-87 MG | ENCODE (ENCFF164HCK, ENCFF334XLV) |
| peripheral blood mononuclear cell | ENCODE (ENCFF475DKC, ENCFF443WJD) |

| | |
|---|---|
| spleen | ENCODE (ENCFF597SJD, ENCFF693OQP, ENCFF921BKL, ENCFF545TFV, ENCFF809RAX, ENCFF398TQO) |
| hepatocyte | ENCODE (ENCFF138JDF, ENCFF797ZIB) |
| neural stem progenitor cell | ENCODE (ENCFF789VZB, ENCFF183XSM) |
| foreskin fibroblast | ENCODE (ENCFF219FYH, ENCFF964QLH) |
| MCF7 | ENCODE (ENCFF009GDJ, ENCFF885LEQ) |
| BJ | ENCODE (ENCFF800TGS, ENCFF839MWS) |
| fibroblast of breast | ENCODE (ENCFF355QNL, ENCFF281CHW) |
| smooth muscle cell | ENCODE (ENCFF003QAY, ENCFF852RUU) |
| placenta | ENCODE (ENCFF435PHN) |
| renal cortex interstitium | ENCODE (ENCFF656GXW, ENCFF885EHC, ENCFF067NHZ) |
| UCSF-4 | ENCODE (ENCFF219WTP) |
| heart left ventricle | ENCODE (ENCFF860DPP, ENCFF998SEL) |
| NCI-H460 | ENCODE (ENCFF322HJX, ENCFF011JTT, ENCFF522SUJ) |
| cardiac muscle cell | ENCODE (ENCFF761ACO, ENCFF509IXK) |
| HEK293 | ENCODE (SRR5137672, SRR5137671, SRR5137670) |
| mesodermal cell | ENCODE (ENCFF553EAV, ENCFF749RUQ) |
| adipose tissue | ENCODE (ENCFF878UHQ, ENCFF862LZV, ENCFF732LRY, ENCFF272HOG) |

| | |
|---|---|
| GM12878 | ENCODE (ENCFF200USH, ENCFF905XDJ, ENCFF644DIQ, ENCFF456WWG, ENCFF876LUX, ENCFF886FDY, ENCFF121ZOP, ENCFF599JTV, ENCFF545OJE, ENCFF853VUK, ENCFF392CRO, ENCFF102NLY, ENCFF315WZE, ENCFF477JYI, ENCFF902UYP, ENCFF550OHK) |
| CD14-positive monocyte | ENCODE (ENCFF397DFK, ENCFF299BIL, ENCFF219ECV) |
| skeletal muscle myoblast | ENCODE (ENCFF505GUJ, ENCFF354AZS) |
| skin fibroblast | ENCODE (ENCFF694YJO, ENCFF202DSC, ENCFF458PJZ, ENCFF959PRH) |
| muscle of leg | ENCODE (ENCFF125HHE, ENCFF068NMI, ENCFF393OUO, ENCFF884IWB, ENCFF316SOJ, ENCFF559DXJ, ENCFF622TLD, ENCFF114CDE, ENCFF398PFM) |
| layer of hippocampus | ENCODE (ENCFF323NAG) |
| HT1080 | ENCODE<br>ENCFF241KQK, ENCFF284XTA) |
| HFFc6 | 4DN (4DNFI5MR6C3G, 4DNFIF3H5ZCH) |
| ectodermal cell | ENCODE (ENCFF034KRQ, ENCFF419KMW, ENCFF691ZYQ, ENCFF768SPT) |
| foreskin keratinocyte | ENCODE (ENCFF680BHT, ENCFF994UBN, ENCFF892CWH, ENCFF051VYX) |
| PFSK-1 | ENCODE (ENCFF635ALW, ENCFF568GAV) |

| | |
|---|---|
| left renal cortex interstitium | ENCODE (ENCFF507HAK, ENCFF068QRU, ENCFF869JVY, ENCFF278ZFZ) |
| SK-N-DZ | ENCODE (ENCFF635RKM, ENCFF594NJL, ENCFF499DSN, ENCFF611SKK) |
| chorion | ENCODE (ENCFF550CHR, ENCFF364QYB, ENCFF672UJB, ENCFF397ZBW) |
| kidney | ENCODE (ENCFF525CMT, ENCFF804WTK, ENCFF099FXO, ENCFF229DFM, ENCFF326MQO) |
| trophoblast | ENCODE (ENCFF768QIJ, ENCFF591XIE, ENCFF270MHZ, ENCFF873XNT) |
| left kidney | ENCODE (ENCFF593SHI) |
| B cell | ENCODE (ENCFF770XDU, ENCFF231GYC, ENCFF485EUP) |
| GM23338 | ENCODE (ENCFF305XIS, ENCFF149CBS) |
| keratinocyte | ENCODE (ENCFF401JWS, ENCFF344FGV, ENCFF065UCN, ENCFF697CPR, ENCFF734GZX, ENCFF345YOV, ENCFF330VCJ, ENCFF165MYR) |
| thymus | ENCODE (ENCFF487KXD, ENCFF118YIZ, ENCFF380IZG, ENCFF237HIP, ENCFF432EQO, ENCFF123LVM, ENCFF608SHV) |
| HepG2 | ENCODE (ENCFF640ZBJ, ENCFF861GCR, ENCFF534SLQ, ENCFF945LNB, ENCFF197XZL, ENCFF874RXH, ENCFF401KRE, ENCFF004HYK) |

| | |
|---|---|
| left lung | ENCODE (ENCFF066QDJ, ENCFF919EYT, ENCFF934RBH, ENCFF643UJO, ENCFF391BNP, ENCFF406OKZ) |
| common myeloid progenitor, CD34-positive | ENCODE (ENCFF690QPA) |
| AG04450 | ENCODE (ENCFF025DRM, ENCFF350BOF) |
| stomach | ENCODE (ENCFF874WKO, ENCFF031EAX, ENCFF881ZXZ, ENCFF953SNL, ENCFF850QFV, ENCFF815DCS, ENCFF082DAH, ENCFF775TWS, ENCFF648ZHB, ENCFF050VJS) |
| pancreas | ENCODE (ENCFF625HJC, ENCFF390JAT, ENCFF971GFG) |
| renal pelvis | ENCODE (ENCFF524ZGN, ENCFF237VRQ, ENCFF013EVX) |
| T-cell | ENCODE (ENCFF158VJT) |
| muscle of trunk | ENCODE (ENCFF073LSZ, ENCFF419PGC) |
| esophagus | ENCODE (ENCFF993JBC, ENCFF566SLH) |
| 293T | ENCODE (SRR12137695, SRR12137698, SRR12137696, SRR12137697) |
| sigmoid colon | ENCODE (ENCFF474EOZ, ENCFF904HJO, ENCFF362GHJ, ENCFF395HHA) |
| spinal cord | ENCODE (ENCFF144IMD, ENCFF400RFD, ENCFF340FSS) |
| forelimb muscle | ENCODE (ENCFF927KFT) |

| | |
|---|---|
| muscle of arm | ENCODE (ENCFF404ZXJ, ENCFF993LHQ, ENCFF350ZWB, ENCFF272NAO, ENCFF726VKQ, ENCFF603FIV, ENCFF905BGI, ENCFF038FYS, ENCFF007LRI, ENCFF513PLX, ENCFF647PIL, ENCFF690YGY) |
| urinary bladder | ENCODE (ENCFF986LPX) |
| H1 | ENCODE (ENCFF059UBK, ENCFF667AIY, ENCFF741PUY, ENCFF235XMZ, ENCFF113VWX, ENCFF915AUQ, ENCFF653XHG) |
| mesenchymal stem cell | ENCODE (ENCFF693WRN, ENCFF290OQE) |
| astrocyte | ENCODE (ENCFF256APB) |
| psoas muscle | ENCODE (ENCFF543WCB, ENCFF630ZKI, ENCFF489IIG) |
| NB4 | ENCODE (SRR6006856, SRR6006857, SRR6006858) |
| fibroblast of skin of abdomen | ENCODE (ENCFF327BZW, ENCFF063EGA) |
| Jurkat clone E61 | ENCODE (ENCFF489SJY, ENCFF558JTV) |
| IMR90 | ENCODE (ENCFF244VME, ENCFF118OFK) |
| mammary epithelial cell | ENCODE (ENCFF380GBC, ENCFF370XTW) |
| SK-MEL-5 | ENCODE (ENCFF620LZN, ENCFF845RTT, ENCFF070UJX, ENCFF448BML) |
| mesendoderm | ENCODE (ENCFF466QUZ, ENCFF044YLS) |
| HUVEC | ENCODE (ENCFF917CHS, ENCFF804WUK, ENCFF650CVW, ENCFF238WEU) |

| | |
|---|---|
| small intestine | ENCODE (ENCFF411RSF, ENCFF769FAA, ENCFF309DZJ, ENCFF845OKW, ENCFF647PTK, ENCFF686JQP ,ENCFF537WNL, ENCFF080HMB, ENCFF256ELA, ENCFF970SOK) |
| heart | ENCODE (ENCFF227RBV, ENCFF089NNQ, ENCFF496OSH) |
| muscle of back | ENCODE (ENCFF706VUD, ENCFF382FLL, ENCFF313PHN, ENCFF302HTI, ENCFF282SUS, ENCFF424LYQ, ENCFF115ORA, ENCFF195KMJ, ENCFF185TFE, ENCFF860PYR) |
| neurosphere | ENCODE (ENCFF153QRP, ENCFF707AMK, ENCFF993ZTH) |
| K562 | ENCODE (ENCFF490IGF, ENCFF022QGS, ENCFF768TKT, ENCFF172GIN, ENCFF026BMH, ENCFF868MFR, ENCFF937GNL, ENCFF047WAI, ENCFF427EWZ, ENCFF342LXD, ENCFF472EUD, ENCFF185UMS, ENCFF461HPX, ENCFF156DDL) |
| CD8-positive, alpha-beta T cell | ENCODE (ENCFF372OMD, ENCFF088DIY) |
| cerebellum | ENCODE (ENCFF777IQQ) |
| mammary stem cell | ENCODE (ENCFF692TAL) |
| amnion | ENCODE (ENCFF144PZJ, ENCFF443KUS, ENCFF416MQP) |
| placental basal plate | ENCODE (ENCFF345ADJ, ENCFF457JWP, ENCFF434YJO, ENCFF026SBP) |

| | | |
|---|---|---|
| | HeLa-S3 | ENCODE (ENCFF846THO, ENCFF796REI, ENCFF010RHX, ENCFF206NFZ, ENCFF344HRY, ENCFF514TNR, ENCFF922LIV, ENCFF101GPJ) |
| | BE2C | ENCODE (ENCFF396XRF, ENCFF238HUZ) |
| | HUES64 | ENCODE (ENCFF339WEH, ENCFF682DUY) |
| | left renal pelvis | ENCODE (ENCFF126KII, ENCFF602NKQ, ENCFF720JRI, ENCFF774PTF) |
| | mole | ENCODE (ENCFF726KPQ) |
| | right renal pelvis | ENCODE (ENCFF033EGX, ENCFF887YTL, ENCFF494FTI, ENCFF896OFQ) |
| | trophoblast cell | ENCODE (ENCFF342LYI, ENCFF760HDK) |
| | Purkinje cell | ENCODE (ENCFF683XBG, ENCFF890PPZ) |
| | CD4-positive, alpha-beta T cell | ENCODE (ENCFF760LWW, ENCFF794NBU) |
| | large intestine | ENCODE (ENCFF615SUS, ENCFF688MYF, ENCFF554XMC, ENCFF359YQU, ENCFF972VRS, ENCFF458LNT, ENCFF858BYE) |
| | right cardiac atrium | ENCODE (ENCFF635SOC) |
| CTCF ChIA-PET for 32 cell lines or primary cells | CD4-positive, alpha-beta T cell | ENCODE (ENCSR345UIQ) |
| | activated CD8-positive, alpha-beta T cell (treated with anti-CD3 and anti-CD28 coated beads for 36 hours) | ENCODE (ENCSR408MRB) |
| | IMR-90 | ENCODE (ENCSR076TTY) |
| | OCI-LY7 | ENCODE (ENCSR401JWQ) |
| | A673 | ENCODE (ENCSR549TMF) |
| | CD4-positive, alpha-beta memory T cell | ENCODE (ENCSR106INW) |
| | CD8-positive, alpha-beta T cell | ENCODE (ENCSR180GEY) |

| | |
|---|---|
| HUVEC | ENCODE (ENCSR404FPI) |
| activated CD8-positive, alpha-beta memory T cell (treated with anti-CD3 and anti-CD28 coated beads for 36 hours) | ENCODE (ENCSR697ZHG) |
| activated CD8-positive, alpha-beta memory T cell (treated with 10 ng/mL Interleukin-2 for 5 days, anti-CD3 and anti-CD28 coated beads for 7 days) | ENCODE (ENCSR378IUJ) |
| activated CD8-positive, alpha-beta T cell (treated with 10 ng/mL Interleukin-2 for 5 days, anti-CD3 and anti-CD28 coated beads for 7 days) | ENCODE (ENCSR531UMR) |
| naive thymus-derived CD4-positive, alpha-beta T cell | ENCODE (ENCSR120LMS) |
| K562 | ENCODE (ENCSR597AKG) |
| activated CD4-positive, alpha-beta memory T cell (treated with 10 ng/mL Interleukin-2 for 5 days, anti-CD3 and anti-CD28 coated beads for 7 days) | ENCODE (ENCSR291RCO) |
| activated CD4-positive, alpha-beta T cell (treated with 10 ng/mL Interleukin-2 for 5 days, anti-CD3 and anti-CD28 coated beads for 7 days) | ENCODE (ENCSR962FFY) |
| activated T-cell (treated with 50 U/mL Interleukin-2 for 72 hours, anti-CD3 and anti-CD28 coated beads for 72 hours) | ENCODE (ENCSR411UEL, ENCSR038GON) |
| CD8-positive, alpha-beta memory T cell | ENCODE (ENCSR187PXW) |
| T-cell | ENCODE (ENCSR545NUL, ENCSR592BWZ) |
| Caco-2 | ENCODE (ENCSR185PEE) |

| | | |
|---|---|---|
| | WTC11 | ENCODE (ENCSR016VPZ) |
| | A549 | ENCODE (ENCSR911ZMB) |
| | GM12878 | ENCODE (ENCSR184YZV) |
| | activated CD4-positive, alpha-beta memory T cell (treated with anti-CD3 and anti-CD28 coated beads for 36 hours) | ENCODE (ENCSR154MAJ) |
| | MCF10A | ENCODE (ENCSR403ZYJ) |
| | MCF-7 | ENCODE (ENCSR200VHL) |
| | activated B cell (treated with 0.5 μM CpG ODN for 24 hours) | ENCODE (ENCSR494NNF) |
| | activated naive CD4-positive, alpha-beta T cell (treated with anti-CD3 and anti-CD28 coated beads for 36 hours) | ENCODE (ENCSR731OFF) |
| | HCT116 | ENCODE (ENCSR278IZK) |
| | activated CD4-positive, alpha-beta T cell (treated with anti-CD3 and anti-CD28 coated beads for 36 hours) | ENCODE (ENCSR093CTT) |
| | HepG2 | ENCODE (ENCSR411IVB) |
| | Panc1 | ENCODE (ENCSR145PYF) |
| | B-cell | ENCODE (ENCSR536ZNI) |
| RNAPII ChIA-PET for 32 cell lines or primary cells | CD4-positive, alpha-beta T cell | ENCODE (ENCSR448ZLA) |
| | activated CD8-positive, alpha-beta T cell (treated with anti-CD3 and anti-CD28 coated beads for 36 hours) | ENCODE (ENCSR982KEM) |
| | IMR-90 | ENCODE (ENCSR966RPQ) |
| | OCI-LY7 | ENCODE (ENCSR882BUM) |
| | A673 | ENCODE (ENCSR623KNI) |
| | CD4-positive, alpha-beta memory T cell | ENCODE (ENCSR569TBN) |
| | CD8-positive, alpha-beta T cell | ENCODE (ENCSR185VQH) |
| | HUVEC | ENCODE (ENCSR080OMN) |

| | |
|---|---|
| activated CD8-positive, alpha-beta memory T cell (treated with anti-CD3 and anti-CD28 coated beads for 36 hours) | ENCODE (ENCSR041UPG) |
| activated CD8-positive, alpha-beta memory T cell (treated with 10 ng/mL Interleukin-2 for 5 days, anti-CD3 and anti-CD28 coated beads for 7 days) | ENCODE (ENCSR114KEO) |
| activated CD8-positive, alpha-beta T cell (treated with 10 ng/mL Interleukin-2 for 5 days, anti-CD3 and anti-CD28 coated beads for 7 days) | ENCODE (ENCSR217TFN) |
| naive thymus-derived CD4-positive, alpha-beta T cell | ENCODE (ENCSR763OCG) |
| K562 | ENCODE (ENCSR880DSH) |
| activated CD4-positive, alpha-beta memory T cell (treated with 10 ng/mL Interleukin-2 for 5 days, anti-CD3 and anti-CD28 coated beads for 7 days) | ENCODE (ENCSR159PXF) |
| activated CD4-positive, alpha-beta T cell (treated with 10 ng/mL Interleukin-2 for 5 days, anti-CD3 and anti-CD28 coated beads for 7 days) | ENCODE (ENCSR733XLQ) |
| activated T-cell (treated with 50 U/mL Interleukin-2 for 72 hours, anti-CD3 and anti-CD28 coated beads for 72 hours) | ENCODE (ENCSR165FXG, ENCSR891IMI, ENCSR538SBO) |
| CD8-positive, alpha-beta memory T cell | ENCODE (ENCSR149TQU) |
| T-cell | ENCODE (ENCSR722NQM, ENCSR743YTL) |
| Caco-2 | ENCODE (ENCSR713NCY) |
| WTC11 | ENCODE (ENCSR972JTN) |

| | | |
|---|---|---|
| | A549 | ENCODE (ENCSR138NSW) |
| | GM12878 | ENCODE (ENCSR905HWW) |
| | activated CD4-positive, alpha-beta memory T cell (treated with anti-CD3 and anti-CD28 coated beads for 36 hours) | ENCODE (ENCSR877ZRR) |
| | MCF10A | ENCODE (ENCSR499JGQ) |
| | MCF-7 | ENCODE (ENCSR059HDE) |
| | activated B cell (treated with 0.5 μM CpG ODN for 24 hours) | ENCODE (ENCSR096ZPB) |
| | activated naive CD4-positive, alpha-beta T cell (treated with anti-CD3 and anti-CD28 coated beads for 36 hours) | ENCODE (ENCSR402NAQ) |
| | HCT116 | ENCODE (ENCSR035PVZ) |
| | activated CD4-positive, alpha-beta T cell (treated with anti-CD3 and anti-CD28 coated beads for 36 hours) | ENCODE (ENCSR314TNQ) |
| | HepG2 | ENCODE (ENCSR789ZIJ, ENCSR857MYZ) |
| | Panc1 | ENCODE (ENCSR447IUA) |
| | B-cell | ENCODE (ENCSR172WWJ) |
| DNase-Seq peaks for 32 cell lines or primary cells | CD4-positive, alpha-beta T cell | ENCODE (ENCFF138ZRF, ENCFF396CHT, ENCFF348SDZ, ENCFF147ZBC) |
| | activated CD8-positive, alpha-beta T cell (treated with anti-CD3 and anti-CD28 coated beads for 36 hours) | ENCODE (ENCFF834QBM) |
| | IMR-90 | ENCODE (ENCFF800DVI, ENCFF525PWS) |
| | OCI-LY7 | ENCODE (ENCFF196DIZ, ENCFF903CVK) |
| | A673 | ENCODE (ENCFF607YBN, ENCFF355NVJ, ENCFF606OTK) |

| | |
|---|---|
| CD4-positive, alpha-beta memory T cell | ENCODE (ENCFF157MKK, ENCFF967NLQ, ENCFF386UFI, ENCFF010BIN) |
| CD8-positive, alpha-beta T cell | ENCODE (ENCFF070WZM, ENCFF792FLJ, ENCFF326MXR, ENCFF017FZE) |
| HUVEC | ENCODE (ENCFF833UNB, ENCFF136MYL, ENCFF406AWN) |
| activated CD8-positive, alpha-beta memory T cell (treated with anti-CD3 and anti-CD28 coated beads for 36 hours) | ENCODE (ENCFF401WTQ, ENCFF848EBC, ENCFF713XCT, ENCFF810ZJY) |
| activated CD8-positive, alpha-beta memory T cell (treated with 10 ng/mL Interleukin-2 for 5 days, anti-CD3 and anti-CD28 coated beads for 7 days) | ENCODE (ENCFF792PYU) |
| activated CD8-positive, alpha-beta T cell (treated with 10 ng/mL Interleukin-2 for 5 days, anti-CD3 and anti-CD28 coated beads for 7 days) | ENCODE (ENCFF596UGI) |
| naive thymus-derived CD4-positive, alpha-beta T cell | ENCODE (ENCFF885PJD, ENCFF937FLI, ENCFF901UGO, ENCFF951XVJ) |
| K562 | ENCODE (ENCFF274YGF, ENCFF264UFX, ENCFF185XRG, ENCFF807ICZ) |
| activated CD4-positive, alpha-beta memory T cell (treated with 10 ng/mL Interleukin-2 for 5 days, anti-CD3 and anti-CD28 coated beads for 7 days) | ENCODE (ENCFF552TYH) |
| activated CD4-positive, alpha-beta T cell (treated with 10 ng/mL Interleukin-2 for 5 days, anti-CD3 and anti- | ENCODE (ENCFF765OGD) |

| | |
|---|---|
| CD28 coated beads for 7 days) | |
| activated T-cell (treated with 50 U/mL Interleukin-2 for 72 hours, anti-CD3 and anti-CD28 coated beads for 72 hours) | ENCODE (ENCFF725EBY, ENCFF168GAN) |
| CD8-positive, alpha-beta memory T cell | ENCODE (ENCFF931AKV, ENCFF375PHK, ENCFF041GGN, ENCFF224KBE) |
| T-cell | ENCODE (ENCFF873FYV, ENCFF729DOV, ENCFF933QZD, ENCFF839QLN) |
| Caco-2 | ENCODE (ENCFF810WJX, ENCFF637OJO, ENCFF948AMM, ENCFF579UXQ) |
| WTC11 | ENCODE (ENCFF668BJR, ENCFF854DSG) |
| A549 | ENCODE (ENCFF128ZVL, ENCFF410KIB, ENCFF302JWZ) |
| GM12878 | ENCODE (ENCFF338SAU, ENCFF759OLD) |
| activated CD4-positive, alpha-beta memory T cell (treated with anti-CD3 and anti-CD28 coated beads for 36 hours) | ENCODE (ENCFF067SGU) |
| MCF10A | ENCODE (ENCFF667FTX) |
| MCF-7 | ENCODE (ENCFF107HQA, ENCFF886OJN, ENCFF835KCG, ENCFF536CIK) |
| activated B cell (treated with 0.5 μM CpG ODN for 24 hours) | ENCODE (ENCFF489NNB) |
| activated naive CD4-positive, alpha-beta T cell (treated with anti-CD3 and anti-CD28 coated beads for 36 hours) | ENCODE (ENCFF261QWU) |

| | |
|---|---|
| HCT116 | ENCODE (ENCFF356GTJ, ENCFF240LRP) |
| activated CD4-positive, alpha-beta T cell (treated with anti-CD3 and anti-CD28 coated beads for 36 hours) | ENCODE (ENCFF107XDE) |
| HepG2 | ENCODE (ENCFF748QCZ, ENCFF897NME, ENCFF453AEP) |
| Panc1 | ENCODE (ENCFF415BYA, ENCFF842AOT) |
| B-cell | ENCODE (ENCFF262LZH, ENCFF245RKH, ENCFF526MQV, ENCFF248ACA) |

**Annotation of enhancer regions in different cell types**

To define candidate enhancer regions in each cell type, we first downloaded the total set of human cis-regulatory elements (cCREs) from the ENCODE data portal website using the following link https://screen.encodeproject.org/. We then extracted all regions marked as ELS (enhancer-like signatures) from the downloaded file. Finally, enhancer regions in each cell type were defined as a subset of these regions that overlap with ATAC-Seq or DNase-Seq peaks in corresponding cells, based on data availability for those cells.

**Gene expression breadth analysis**

To explore the gene expression profiles of a specific gene set across a diverse range of cell type or tissues, we collected RNA-Seq datasets for 116 human cell types or tissues (from ENCODE, see table of datasets for this section above). The transcripts per million (TPM) values were used to measure gene transcription levels. To normalize the RNA-Seq data, we first applied a logarithm transformation to the original TPM values using the formula log2(TPM+1) for each sample, and then quantile-normalized the transformed TPM values across all samples.

In each sample, genes with a normalized TPM value greater than 3 were considered expressed in the corresponding sample, and the gene expression breadth is defined as the number of samples in which a gene is expressed.

### 3. Method for SPIN states identification and analysis

#### Data availability

| Cell Type | Experiment Type | Data Type | Data Source/Download Link | Note1 |
|---|---|---|---|---|
| H1-ESC | SON TSA-seq | Signal (BigWig) | 4DN (4DNESUTK6QWG, 4DNESC3D6NGQ) | |
| | LaminB TSA-seq | Signal (BigWig) | 4DN (4DNESHZ8WKRX, 4DNESGGXKI1H) | |
| | Nucleous TSA-seq | Signal (BigWig) | 4DN (4DNESO6HFSAD, 4DNES6PANOF4) | |
| | LaminB DamID | Signal (BigWig) | 4DN (4DNESOFQR5FS, 4DNESXKBPZKQ) | |
| | Hi-C | Contact matrix | 4DN (4DNESX75DD7R, 4DNES2M5JIGV) | |
| | H2AFZ ChIP-Seq | Signal (BigWig) | ENCODE (ENCFF758YFI) | |
| | H3K27ac ChIP-Seq | Signal (BigWig) | ENCODE (ENCFF771GNB) | |
| | H3K27me3 ChIP-Seq | Signal (BigWig) | ENCODE (ENCFF277UCT) | |
| | H3K36me3 ChIP-Seq | Signal (BigWig) | ENCODE (ENCFF687LJF) | |
| | H3K4me1 ChIP-Seq | Signal (BigWig) | ENCODE (ENCFF335ZGP) | |
| | H3K4me2 ChIP-Seq | Signal (BigWig) | ENCODE (ENCFF501AUN) | |
| | H3K4me3 ChIP-Seq | Signal (BigWig) | ENCODE (ENCFF698DKQ) | |
| | H3K79me2 ChIP-Seq | Signal (BigWig) | ENCODE (ENCFF640WRD) | |
| | H3K9ac ChIP-Seq | Signal (BigWig) | ENCODE (ENCFF890MIB) | |
| | H3K9me3 ChIP-Seq | Signal (BigWig) | ENCODE (ENCFF358AWN) | |
| | H4K20me1 ChIP-Seq | Signal (BigWig) | ENCODE (ENCFF453OCL) | |
| | 16-fraction Repli-seq G1 | Alignment (Bam) | 4DN (4DNES2W2DXM6) | The smoothed and normalized RT score can be downloaded from GSE137764 |

| | | | | |
|---|---|---|---|---|
| | 16-fraction Repli-seq S1 | Alignment (Bam) | 4DN (4DNES8GLHOD6) | |
| | 16-fraction Repli-seq S2 | Alignment (Bam) | 4DN (4DNESWWMQ1IC) | |
| | 16-fraction Repli-seq S3 | Alignment (Bam) | 4DN (4DNES2LV4C4Q) | |
| | 16-fraction Repli-seq S4 | Alignment (Bam) | 4DN (4DNES2RXY8IB) | |
| | 16-fraction Repli-seq S5 | Alignment (Bam) | 4DN (4DNESVX7IH76) | |
| | 16-fraction Repli-seq S6 | Alignment (Bam) | 4DN (4DNESLOM1MB7) | |
| | 16-fraction Repli-seq S7 | Alignment (Bam) | 4DN (4DNESR9FVS6K) | |
| | 16-fraction Repli-seq S8 | Alignment (Bam) | 4DN (4DNESFMEOEHW) | |
| | 16-fraction Repli-seq S9 | Alignment (Bam) | 4DN (4DNESBZIJJ1E) | |
| | 16-fraction Repli-seq S10 | Alignment (Bam) | 4DN (4DNESDZOL2VU) | |
| | 16-fraction Repli-seq S11 | Alignment (Bam) | 4DN (4DNESJJKOIH4) | |
| | 16-fraction Repli-seq S12 | Alignment (Bam) | 4DN (4DNES4I9OXS7) | |
| | 16-fraction Repli-seq S13 | Alignment (Bam) | 4DN (4DNES6G2SSN2) | |
| | 16-fraction Repli-seq S14 | Alignment (Bam) | 4DN (4DNESGY7RRTV) | |
| | 16-fraction Repli-seq S15 | Alignment (Bam) | 4DN (4DNESYIVVMOC) | |
| | 16-fraction Repli-seq S16 | Alignment (Bam) | 4DN (4DNESIK2HK47) | |
| | iMARGI | Read pairs | 4DN (4DNESOBRUQ12, 4DNES8B3R3P8, 4DNESGRI8A8N, 4DNESNOJ7HY7) | Only interchromosomal repetitive element (RE)-containing chromatin-associated RNAs (caRNAs) read pairs are used |
| HFFc6 | SON TSA-seq | Signal (BigWig) | 4DN (4DNESB5I8TGR, 4DNES85R9TIB) | |
| | LaminB TSA-seq | Signal (BigWig) | 4DN (4DNES16C6XVY, 4DNESMF4T7QQ) | |
| | Nucleous TSA-seq | Signal (BigWig) | 4DN (4DNESGAR9ZBW, 4DNES2RN8ZJ1) | |

| | | | |
|---|---|---|---|
| | LaminB DamID | Signal (BigWig) | 4DN (4DNESOOCOBBA, 4DNESXZ4FW4T) |
| | Hi-C | Contact matrix | 4DN (4DNESNMAAN97, 4DNES2R6PUEK) |
| | Imputed H2AFZ ChIP-Seq | Signal (BigWig) | 4DN (4DNFIB79WDUY) |
| | Imputed H3K27ac ChIP-Seq | Signal (BigWig) | 4DN (4DNFIBNZ478I) |
| | Imputed H3K27me3 ChIP-Seq | Signal (BigWig) | 4DN (4DNFIMJHO898) |
| | Imputed H3K36me3 ChIP-Seq | Signal (BigWig) | 4DN (4DNFIDU8WT76) |
| | Imputed H3K4me1 ChIP-Seq | Signal (BigWig) | 4DN (4DNFIBC1VSF6) |
| | Imputed H3K4me2 ChIP-Seq | Signal (BigWig) | 4DN (4DNFIFB7SHA1) |
| | Imputed H3K4me3 ChIP-Seq | Signal (BigWig) | 4DN (4DNFIKRPF9QP) |
| | Imputed H3K79me2 ChIP-Seq | Signal (BigWig) | 4DN (4DNFIBS37Z1K) |
| | Imputed H3K9ac ChIP-Seq | Signal (BigWig) | 4DN (4DNFINBZ9NNY) |
| | Imputed H3K9me3 ChIP-Seq | Signal (BigWig) | 4DN (4DNFILJFE7WW) |
| | Imputed H4K20me1 ChIP-Seq | Signal (BigWig) | 4DN (4DNFI13RAB7H) |
| | CUT&RUN CTCF | Raw (fastq) | 4DN (4DNES1RQBHPK) |
| | CUT&RUN H2A.Z | Raw (fastq) | 4DN (4DNESHPUFWTR) |
| | CUT&RUN H3K27ac | Raw (fastq) | 4DN (4DNESIMWCLF8) |

| | | | | |
|---|---|---|---|---|
| | CUT&RUN H3K27me3 | Raw (fastq) | 4DN (4DNES8TY5P5P) | |
| | CUT&RUN H3K4me1 | Raw (fastq) | 4DN (4DNESRFWR5SV) | |
| | CUT&RUN H3K4me2 | Raw (fastq) | 4DN (4DNESEQW5QAI) | |
| | CUT&RUN H3K4me3 | Raw (fastq) | 4DN (4DNESPE6J9FU) | |
| | iMARGI | Read pairs | 4DN (4DNES9Y1GHK4) | Only interchromosomal repetitive element (RE)-containing chromatin-associated RNAs (caRNAs) read pairs are used |

**Data acquisition and processing**

We obtained TSA-seq, Lamin-B-DamID, and Hi-C data for H1-hESCs and HFFc6 from the 4DN data portal (http://data.4dnucleom.rog). The data generation and processing pipeline for TSA-seq data is described in {Chen, 2018 #1443}{Zhang, 2021 #1892}. The data generation and processing pipeline for DamID data is described in {Leemans, 2019 #1932}. For the SPIN states inference, we used Hi-C data generated by the formaldehyde (FA) _ disuccinimidyl glutarate (DSG) Hi-C protocol (1% FA followed by incubation with 3 mM DSG) using restriction enzyme DnpII {Akgol Oksuz, 2021 #1887}. We binned TSA-seq, Lamin-B-DamID and Hi-C mapped reads at 25 kb resolution. We then identified significant interactions from the normalized Hi-C data in each cell type previously described {Wang, 2021 #1783}.

**Identifying SPIN states for large-scale genome compartmentalization**

In this work, we used a modified SPIN method to perform joint modeling across multiple cell types. To ensure TSA-seq and DamID scores across different cell types are comparable, we first applied a data normalization method to transform data into a Gaussian or more-Gaussian-like distribution. To do that we identified genomic bins that are spatially stable by calculating the Pearson correlation of interchromosomal Hi-C interactions for each non-overlapping 25 kb genomic bin. Bins were then ranked based on the average Pearson correlation coefficient, and the top 25% were selected as spatially conserved regions. We then obtained TSA-seq or DamID scores for these bins in all cell types and standardized each data track by fitting a power-transformation function. We used the Yeo-Johnson transformation function with the default parameters from the Python scikit-learn package. Next, we modified the framework of SPIN by jointly modeling the probability across multiple cell types. The hidden Markov random field (HMRF) model is defined on an undirected graph $G^c = (V, E^c)$ for each cell type, where in our case $V$ represents non-overlapping 25kb genomic bins and $E^c$ represents the cell-type specific edges (i.e., significant Hi-C interactions) in cell type $c$. For each $i \in V$, $O_i^c \in \mathbb{R}^d$ is a vector with dimension $d$ indicating the observed TSA-seq and DamID signal of this bin in cell type $c$. Each node $i\ inC$ also has a hidden state $H_i^c$ for each cell type, representing its underlying spatial environment relative to different nuclear landmarks that we want to estimate. In this work, we assume that the set of hidden states are shared across cell types. Edges $(i^c, j^c) \in E^c$ in the graph are cell type-specific and there are no edges that are connecting nodes from different cell types. Therefore, the hidden state $H_i^c$ is only dependent on cell-type specific observation $O_i^c$ and the neighbors of node $i$ ($N^c(i) = \{j | j \in V, (i,j) \in E^c\}$) in cell type $c$. The overall objective is to estimate the hidden states $H_i^c$ for all nodes in all cell types that maximize the following joint probability as shown below:

$$P(\overrightarrow{H}, \overrightarrow{O}) \propto \frac{1}{Z} \prod_{c \in 1..4} \left( \prod_{i \in V} P_V(O_i^c | H_i^c) \prod_{(i^c, j^c) \in E^c} P_{E^c}(H_i^c, H_j^c)] \right)$$

To estimate the number of SPIN states, we applied the same approaches as we used in the previous version of the SPIN method {Wang, 2021 #1783}. We used both the Elbow method based on K-means clustering and AIC/BIC scores to search for the optimal number of SPIN states. Both AIC and BIC scores decrease as the number of states increases. We found that the slope of the curve drops close to zero as the number of states exceeds 9. So, we chose 9.

**Processing other epigenomic data**

We downloaded or processed other epigenomic data and compared SPIN states with these datasets. For ChIP-seq datasets, we downloaded the processed p-value tracks from the ENCODE website for H1-hESC and Avocado {Schreiber, 2020 #1904}{Schreiber, 2021 #1905} imputed p-value tracks from the 4DN data portal. Multi-fraction Repli-seq data were collected from {Zhao, 2020 #1908}. For CUT&RUN data, we downloaded raw sequencing reads from the 4DN data portal and processed them using a similar procedure according to the standard ENCODE ChIP-seq pipeline. First, we mapped reads to hg38 reference genome using Bowtie2 (version 2.2.9) with the default parameters. We then used MACS3 to generate p-value tracks as well as peaks for CUT&RUN data. The enrichment score of epigenomic data on SPIN states is determined by the log2 ratio between the average observed signals on each SPIN states over genome-wide expectation.

**SPIN states enriched caRNA sequence features.**

Processing of iMARGI data was performed with iMARGI-Docker {Wu, 2019 #1910}. To quantify enrichment of repetitive element (RE)-containing chromatin-associated RNAs (caRNAs) with specific chromatin states, we computed an enrichment score (log2 observed/expected interaction frequencies) for each RE caRNA class and SPIN states. Observed frequencies were derived from the number of iMARGI read pairs with RNA ends mapped to RE class of interest and DNA ends mapped to SPIN states. Expected frequencies computed as the total number of iMARGI read pairs multiplied by the product of the marginal probabilities of RE class abundance (proportion of all caRNAs mapping to each RE class, irrespective of DNA mapping locations) and SPIN state abundance (proportion of DNA reads mapping to each SPIN state). Only inter-chromosomal iMARGI pairs were analyzed to mitigate potential biases from nascent RNA transcripts interacting with proximal genomic regions.

**Nascent transcription measured by iMARGI**

iMARGI RNA ends coverage are derived from RNA, DNA interactions represented in bedpe files generated by iMARGI docker {Wu, 2019 #1910}. The RNA end abundance bigwig file is generated by calculating the pileup reads coverage (R, coverage function) on the genome using RNA ends only in a strand specific manner.

The iMARGI datasets used for this paper are as follows:

| Cell line | DCIC accession |
|---|---|
| H1 | 4DNESNOJ7HY7 |
| HFFc6 | 4DNES9Y1GHK4 |
| K562 | 4DNESIKCVASO |

The bigwig tracks of iMARGI's RNA reads are as follows:

| Cell line | Treatmen t | Strand | DCIC accession |
|---|---|---|---|
| H1 | Control | Plus | 4DNFIOYVWEYZ |
| H1 | Control | Minus | 4DNFI2LXIREI |

4. **Methods for Integrated Genome Structure Modeling**

**Datasets**

The 4DN accession codes to the input data used in simulating and analyzing the genome structures are given in the table below.

| | **Hi-ESC** | **HFFc6** |
|---|---|---|
| **Ensemble Hi-C** | 4DNESX75DD7R {Akgol Oksuz, 2021 #1887} | 4DNESNMAAN97 {Akgol Oksuz, 2021 #1887} |
| **Lamina DamID** | 4DNESXKBPZKQ | 4DNESXZ4FW4T |
| **SPRITE** | 4DNESASBN1JK {Bhat, 2023 #1958} | 4DNESJYGTI8S {Bhat, 2023 #1958} |
| **TSA-seq** | 4DNFI625PP2A {Zhang, 2021 #1892} | 4DNFI6FTPH5V {Zhang, 2021 #1892} |
| **RNA-seq** | 4DNES3IOYG74<br>GSE75748 {Chu, 2016 #1959} | 4DNESFH3EHTU<br>GSE75748 {Chu, 2016 #1959} |
| **Histone ChIP-seq** | ENCFF986PCY {Roadmap Epigenomics Consortium, 2015 #1286}<br>ENCFF088MXE {Roadmap Epigenomics Consortium, 2015 #1286} | ENCFF426TLD {ENCODE-Project-Consortium, 2012 #1015}<br>ENCFF792IOR {ENCODE-Project-Consortium, 2012 #1015} |

| | | |
|---|---|---|
| | ENCFF084JKQ {Roadmap Epigenomics Consortium, 2015 #1286}<br>ENCFF183MHJ {Roadmap Epigenomics Consortium, 2015 #1286}<br>ENCFF860NVB {Roadmap Epigenomics Consortium, 2015 #1286}<br>ENCFF401PZS {Roadmap Epigenomics Consortium, 2015 #1286}<br>ENCFF156JZY {Roadmap Epigenomics Consortium, 2015 #1286}<br>ENCFF065VIF {Roadmap Epigenomics Consortium, 2015 #1286}<br>ENCFF445UVT {Roadmap Epigenomics Consortium, 2015 #1286}<br>ENCFF488THD {Roadmap Epigenomics Consortium, 2015 #1286}<br>ENCFF780FNS {Roadmap Epigenomics Consortium, 2015 #1286} | ENCFF994SSG {ENCODE-Project-Consortium, 2012 #1015}<br>ENCFF690KUY {ENCODE-Project-Consortium, 2012 #1015}<br>ENCFF995LLA {ENCODE-Project-Consortium, 2012 #1015}<br>ENCFF070SWD {ENCODE-Project-Consortium, 2012 #1015} |

Data preprocessing was performed as described in Boninsegna et al. {Boninsegna, 2022 #1914}, with exception of parameter $f_{maxation} = 16$ for Hi-C preprocessing.

**Genome representation**

The genome is represented at 200 kilobase pair resolution as described in {Boninsegna, 2022 #1914}{Yildirim, 2023 #1916}, resulting in $N = 29838$ chromatin regions, modeled as hard spheres of radius $r_0 = 118nm$. The HFFc6 nucleus is modeled as an ellipsoid of semiaxes $(a, b, c) = (7840, 6480, 2450) nm$, while the the H1ESC envelope is represented by a sphere of radius $R = 5{,}000 nm$.

We define the population of single cell genome structures as a set of $S$ diploid genome structures $X = \{X_1, \ldots, X_S\}$; A genome structure $X_s$ is a set of 3-dimensional vectors $X_s = \{\vec{x}_{is}: \vec{x}_{is} \in R^3, i = 1,2, \ldots, N\}$ representing the center coordinates of each chromatin domain within the structure $s$:, $N$ being the total number of chromatin domains in the genome and $\vec{x}_{is} = (x_{is}, y_{is}, z_{is},) \in R^3$ indicates the coordinates of a 200k base pair genomic region $i$ in structure $s$. We use the notation $I = (i, i')$ to indicate the genomic region, where $i$ and $i'$ represent the two alleles of genomic region $I$.

**Data-driven simulation of genome structures**

Genome structure populations were generated with IGM following procedures described in {Boninsegna, 2022 #1914}. The goal is to determine a population of 1,000 diploid 3D genome structures $X$ statistically consistent with all input data from different available genomics experiments. Given a collection of input data $D$ from different data sources, $D = \{D_k | k = 1, \ldots, 3\}$ (here, ensemble Hi-C, lamina DamID and SPRITE, see data availability table above), we aim to estimate the structure population $X$ such that the likelihood $P(\{D_k\}|X)$ is maximized. To represent missing information at single cell and homologous chromosome level, we introduce data indicator tensors $D^* = \{D_k^* | k = 1, \ldots, K = 3\}$, which augment missing information about allelic copies in single cells. Thus, the latent variables are a detailed expansion of the diploid and single-structure representation.

To determine a population of 3D genome structures consistent with all experimental data, we formulate a hard Expectation-Maximization (EM) problem, where we jointly optimize all genome structure coordinates and all latent variables. Given $\{D_k\}$, we aim to estimate the structure population $X$ and latent indicator variables $\{D_k^*\}$ such that the likelihood $P(X)$ is maximized. We thus aim to find the optimal structures and the optimal latent variables which satisfy: $\widehat{\boldsymbol{X}}, \widehat{\mathfrak{D}^*} = \arg\max_{\mathbf{X}, \mathcal{D}^*} P(\mathfrak{D}, \mathfrak{D}^* | \boldsymbol{X}) = \arg\max_{\mathbf{X}, \mathcal{D}^*} \prod_k P(\mathcal{D}_k | \mathcal{D}_k^*, \boldsymbol{X}) \cdot \prod_k P(\mathcal{D}_k^* | \boldsymbol{X})$ , This is a high dimensional, hard Expectation Maximization problem and it is solved iteratively by implementing a series of optimization strategies for scalable and effective model estimation. Any iteration first optimizes the latent variables, by using the input data $\{D_k\}$ and the coordinates of all genomic regions $X^{(t)}$ from the previous iteration step, i.e., $\mathfrak{D}^{*(t+1)} = \arg max_{\mathfrak{D}^*} P(\mathfrak{D}|\mathfrak{D}^*, \boldsymbol{X}^t) P(\mathfrak{D}^*|\boldsymbol{X}^t)$. Then, coordinates of the genomic regions are optimized, based on the data deconvolution $D^{*(t)}$, i.e. $\boldsymbol{X}^{t+1} =$

$argmax_X\, P(\mathfrak{D} |\, \mathfrak{D}^{*(t)}, \boldsymbol{X}) P(\, \mathfrak{D}^{*(t)} | \boldsymbol{X})$, and additional constraints such as the volume confinement effect by the nuclear envelope, chromosomal chain connectivity and excluded volume. The process is iterated until convergence is reached. Overall, the data deconvolution process ensures that the structure population expresses the single-cell variability of genome organization, while also aggregately reproducing the ensemble behavior (e.g, the ensemble contact probabilities). Details on the probabilistic formulation underlying the optimization process and how that is designed and implemented for the different data sources can be found in Boninsegna et al. {Boninsegna, 2022 #1914}, and accompanying Supporting Information.

**Structural features**

The population of 1,000 single cell 3D genome structures was used to calculate a host of structural features $f$ that characterize the folding of each genomic region. All features and their cell-to-cell variability are calculated as described in {Boninsegna, 2022 #1914}{Yildirim, 2023 #1916}, unless otherwise noted.

Variability If applicable, cell-to-cell variability $\delta f_I$ of structural feature $f$ for a chromatin region $I$ (from chromosome $c$) is defined as: $\delta f_I = \frac{\sigma[f]_I}{\underline{\sigma}[f]_c}$, $\sigma[f]_I$ indicating the standard deviation of the feature value across the population and $\underline{\sigma}[f]_c$ being the mean standard deviation of the feature values of all regions within the same chromosome. $\delta f_I > 0$ $(< 0)$ indicates high (low) variability of that feature at locus $I$.

- **RAD: Normalized average radial position and $\delta RAD$ as its cell-to-cell variability** is calculated as the normalized radial distance of a locus $I$ to the nuclear center averaged over all structures in the population:

$$RAD_I = \frac{1}{S}\sum_{s}\frac{1}{2}\sum_{i\in I} r_{is}$$

$r^2{}_{is} = (\frac{x_{is}}{a})^2 + (\frac{y_{is}}{b})^2 + (\frac{z_{is}}{c})^2$ being the squared radial distance of locus $i$ in structure $s$, and $(a, b, c)$ being the nuclear semi-axes. The cell-to-cell variability of the radial position is defined $\delta RAD_I$.

**Local folding properties** These features encode local properties of the chromatin fiber and chromatin-chromatin interactions.

- **Local chromatin fiber compaction (Rg)** indicates the chromatin local compactness. If $R_g[i,s]$ indicates the radius of gyration of a 1Mb chromatin segment centered at the locus of interest in structure $s$, then

$$Rg[I] = \frac{1}{S}\sum_{S}\frac{1}{2}\sum_{i\in I} R_g[i,s]$$

The compaction variability is denoted with $\delta Rg$.

- **Interchromosomal contact probability (ICP)** indicates the average fraction of trans interactions out of all contacts formed by a genomic region $I$,

$$ICP[I] = \frac{1}{S}\sum_{S}\frac{1}{2}\sum_{i\in I}\frac{n^s{}_{i,trans}}{n^s{}_{i,trans} + n^s{}_{i,cis}}$$

$n^s{}_{i,trans}(n^s{}_{i,cis})$ being the number of trans (cis) contacts formed by region $i$ in structure $s$.

- **Interior localization frequency (ILF)** indicates the fraction of structures in which a locus $I$ (either copy) occupies an inner position,

$$ILF[I] = \frac{1}{S}\sum_{S}\theta(RAD_i \leq \frac{1}{2} \ or \ RAD_{i'} \leq \frac{1}{2})$$

$\theta$ being the Heaviside function

- **Median trans AB ratio (transAB)** For each chromatin region $i$ we define its “trans” A $n^t{}_{is,A}$ and B $n^t{}_{is,B}$ neighborhoods as $n^t{}_{is,A}(n^t{}_{is,B}) = \#\{j: chr[i] \neq chr[j] \wedge |x_{is} - x_{js}| \leq 500\, nm \ \wedge j \in A(B)\}$; $j \in A/B$ indicates locus $j$ is assigned to compartment A/B. The median trans AB ratio for that region across the population is computed by pooling the values from all homologues and structures,

$$transAB_I = median[\{\frac{n^t{}_{is,A}}{n^t{}_{is,B}}\}_{i\in I,s}]$$

The values are then rescaled so that $0 \leq transAB_I \leq 1$.

**Prediction of nuclear body's locations using Markov clustering**

A chromatin interaction network (CIN) is calculated for nuclear body associated chromatin regions as described in {Yildirim, 2023 #1916}. Speckle associated chromatin regions are defined as the top 10% 200 kb regions with highest SON-TSA seq signal, nucleolus associated chromatin regions are 200-kb regions overlap with nucleolus organizing regions identified in {Németh, 2010 #1019}. Spatial partitions of nuclear bodies are further calculated via the Markov Clustering Algorithm (MCL). Specifically, MCL clustering is performed for each nuclear body's CIN with mcl tool in the MCL-edge software {Enright, 2002 #1078}. Speckle locations are defined as the geometric center of speckle partitions identified by MCL in speckle CINs. nucleolus locations are identified following the same protocol in nucleolus CINs. Only spatial partitions with size larger than three chromatin regions are considered for downstream analysis.

<u>Structure features defining the location of genomic regions with respect to nuclear bodies and compartments:</u>

**SpD**, **NuD**, **LaD** define the population averaged distance of a genomic region to the nearest nuclear speckle, nucleolus or the nuclear envelope, respectively, while **SAF**, **NAF**, **LAF** quantify the frequency with which a genomic region is in association with a speckle, nucleus or the nuclear envelope in the population of cells. Approximate locations of nuclear speckles and nucleoli are predicted in each single cell structure following a procedure described in {Yildirim, 2023 #1916}. Specifically, we identified locations of nuclear bodies in single cells by the geometric centers of highly connected subgraphs determined from a chromatin Interaction network, where each node represents a chromatin region with high probability to be associated with a specific nuclear body and edges if their distances is smaller than an interaction cutoff. Details of the procedure are described in {Yildirim, 2023 #1916}.

- **Average distance to the lamina (LAD) and its cell-to-cell variability (**$\delta LAD$**)** is the (normalized) radial distance of a locus $I$ to the nuclear lamina averaged averaged over the cell population:

$$LAD_I = \frac{1}{S}\sum_{s}\frac{1}{2}\sum_{i \in I}(1 - r_{is})$$

The cell-to-cell variability is defined by $\delta LAD_I$

- **Average distances to closest speckle (SpD) and nucleolus (NuD) and their cell-to-cell variabilities (**$\delta SpD, \delta NuD$**)**

$$SpD_I = \frac{1}{S}\sum_{s}\frac{1}{2}\sum_{i\in I} d_{is}{}^{Sp} \;, \qquad NuD_I = \frac{1}{S}\sum_{s}\frac{1}{2}\sum_{i\in I} d_{is}{}^{Nu}$$

where $d_{is}{}^{Sp}$ ($d_{is}{}^{Nu}$) is the distance of genomic region $i$ to the predicted closest speckle (or nucleolus) in structure $s$. The related variability across the population is $\delta SpD_I(\delta NuD_I)$.

- **Association frequencies with nuclear bodies (SAF, LAF, NAF)** The speckle association frequency is defined as:

$$SAF_I = \frac{1}{S}\sum_{s}\frac{1}{2}\sum_{i\in I} \theta(d^{Sp}{}_{is} \le d_{Sp})$$

where $d_{Sp} = 500nm$ and $\theta$ is the Heaviside distribution. Analogous formulas are valid for association frequencies of genomic regions with the lamina (LAF) and nucleoli (NAF):

$$LAF_I = \frac{1}{S}\sum_{s}\frac{1}{2}\sum_{i\in I} \theta(d^{Lam}{}_{is} \le d_{Lam})\,, \qquad NAF_I = \frac{1}{S}\sum_{s}\frac{1}{2}\sum_{i\in I} \theta(d^{Nu}{}_{is} \le d_{Nu})$$

With $d_{Lam} = 0.2RAD$ and $d_{Nu} = 1000\,nm$.

**Calculation of enrichment scores for expression/genes/SPIN groups for structure features.** All structure feature values are min-max normalized to scale the feature value to a 0-1 range. For structure features RAD, SpD and NuD the normalized value is subtracted by 1 to signify lower value for closer proximity to the respective nuclear bodies.

To calculate the enrichment fold of a structure feature in a selected group (either selected based on gene expression, genes, SPIN, etc.), we calculated the log2 ratio of the feature mean within this group over the average mean value calculated from 100 random permutations of chromatin regions in the genome.

**Dimension reduction of structural features with t-Distributed Stochastic Neighbor Embedding (tSNE).**

Structure features RAD, LAF, TransAB, RG, SpD, ICP are normalized by Z-score and combined in feature vector for each genomic region. The tSNE algorithm in python *scikit-learn* library is then applied to the structure feature vector of all chromatin regions with the following parameters: verbosity level of 1, perplexity of 80, maximum of 300 iterations for the optimization, and random state seed of 123 to ensure reproducibility. The first two components are selected for 2D visualization.

**The categorization of gene into different expression groups**

Expected gene counts were estimated by the RSEM package from the alignment bam file and are then normalized by using the NOIseq R package into RPKM values. For genes with non-zero RPKM value, the lower threshold for expression is defined as the 25 percentile (Q1) of the RPKM values; genes with expression level above 0 but below the lower threshold are defined as lowly expressed genes. The upper threshold for expression is defined as the 75 percentile (Q3) of the RPKM values; genes with expression level above the upper threshold are defined as highly expressed genes.

**Calculated enhancer features**

**E:** The number of active enhancers within a 200 kb window where the gene TSS is located.
**E/G:** The number of active enhancers within a 200 kb window normalized by the total number of transcription starts sites of genes.
**$E^N$_Inter/G:** The number of active enhancers from other chromosomes that are located within a spatial distance of 350 nm, normalized by the number of transcription starts sites of genes within the local 200 kb window.
**$E^N$_Intra/G:** The number of active enhancers within the same chromosome that are located within a spatial distance of 350 nm, normalized by the number of transcription starts sites of genes within the local 200 kb window.
**$E^N$_Intra(<2Mb)/G:** The number of active enhancers within a 2Mb sequence distance on the same chromosome that are located within a spatial distance of 350 nm, normalized by the number of transcription starts sites of genes within the local 200 kb window.

### 5. Methods for single cell 3D genome analysis

#### Cell culture

The modified WTC-11 (GM25236) hiPS cell line with GFP tagged AAVS1 locus (clone 6 and clone 28) was cultured following the 4DN approved SOP (https://data.4dnucleome.org/protocols/d5889062-ec16-4246-9606-8d51f6b02dfa/) with two minor differences: 1) For Penicillin/Streptomycin, catalog #15140-122 (Gibco) was used with a final concentration of 1% (v/v); 2) The density of seeding cells into 6-well plate was 50-100K.

**Generating scHi-C data**

The scHi-C libraries were prepared using methods previously described with slight modifications {Flyamer, 2017 #1434} In brief, 1-3 million crosslinked WTC-11 cells were first lysed and permeabilized by 0.5% SDS. Then the cells were incubated overnight at 37 °C with 300 U MboI followed by proximity ligation with T4 ligase at room temperature with slow rotation for 4 hours. Then the nuclei were stained with Hoechst and the single 2N nuclei were sorted by FACS into wells of 96-well plate. After overnight reverse crosslinking at 65 °C, the 3C-ligated DNA in each cell was amplified using GenomiPhi v2 DNA amplification kit (GE Healthcare) for 4.5 hours. After purification with AMPure XP magnetic beads and quantification, 10ng WGA product was used to construct a library with Tn5. The detailed experimental procedures can be found in 4DN portal (https://data.4dnucleome.org/protocols/3286b08d-d1d6-4853-a201-7dd08400d357/).

ScHi-C data can be found here:

For clone 6:

https://data.4dnucleome.org/experiment-set-replicates/4DNESJQ4RXY5/

and for clone 28:

https://data.4dnucleome.org/experiment-set-replicates/4DNESF829JOW/

**The Strings and Binders (SBS) polymer model of the studied *DPPA* locus**

To investigate at the single-molecule level the 3D folding of the *DPPA* locus (chr3: 108.3Mb-110.3Mb) in WTC-11 pluripotent stem cells, we used the Strings and Binders (SBS) polymer model {Nicodemi, 2009 #1925}{Barbieri, 2012 #1781}. In the SBS, a chromatin region is represented as a self-avoiding polymer chain of beads, along which different types of binding sites are located for diffusing cognate molecular binders that can bridge them. The specific attractions between the binders and the polymer binding sites drive the folding of the system via thermodynamic mechanisms of polymer phase separation {Conte, 2020 #1924}. The model binding domains are determined by using the PRISMR algorithm, which infers the optimal, i.e., minimal, sets of different types of polymer binding sites by taking as input only bulk Hi-C contact data {Bianco, 2018 #1945}. In our studied 2Mb wide *DPPA* locus, PRISMR returned 10 different binding domains. To derive a statistical ensemble of in-silico *DPPA* single-molecule 3D conformations, we performed massive Molecular Dynamics (MD) simulations. In the MD implementation of the model, the system of polymer beads and binders is subject to a stochastic Langevin dynamics based on classical interaction potentials of polymer physics with standard parameters {Kremer, 1990 #1962}{Conte, 2020

#1924}. We ran the SBS simulations up to $10^8$ MD time iteration steps, when stationarity is fully reached. To ensure statistical robustness, we collected up to $10^3$ independent model conformations in the steady-state. We used the free available LAMMPS software (v. 30july2016) to run MD simulations highly optimized for parallel computing {Thompson, 2022 #1963}.

### 6. Methods for the section "Relationship among A/B compartments, SPIN states, TADs/subTADs, loops, and replication timing"

#### A/B compartment detection

We called compartments in Hi-Cv2.5 data generated from H1 human ES cells (H1-hESC; {Akgol Oksuz, 2021 #1887} (https://data.4dnucleome.org/files-processed/4DNFIOUDCJRH/) via eigenvector decomposition on each 25 kb chromosomal balanced matrix. We then normalized each matrix by an expected distance dependence mean counts value with removal of rows or columns with less than 2% non- zero counts coverage. We transformed to a z-score each off-diagonal count and a Pearson correlation matrix was computed. Subsequently, we performed eigenvector decomposition on the z-scored Pearson correlation matrix using LA.eig() (linalg package in numpy), selecting the eigenvector with the largest eigenvalue. We identified inflection points demarcating boundaries of compartments by genomic coordinates with a transition in eigenvector sign. We assigned compartments to either an A or B identity by collecting intervals of same eigenvector sign orientation (positive or negative) and counting total number of unique genes per direction then reassigning those with greater gene number intersection as A and the lesser as B.

#### TAD/subTAD detection

We called TADs and subTADs as previously reported {Norton, 2018 #1620}{Zhang, 2019 #1748}{Emerson, 2022 #1935}{Chandrashekar, 2024 #1964} in 10 kb binned Hi-Cv2.5 data generated from H1-h1ESC (https://data.4dnucleome.org/files-processed/4DNFI82R42AD/)) using 3DNetMod (https://bitbucket.org/creminslab/cremins_lab_tadsubtad_calling_pipeline_11_6_2021), as previously described (https://data.4dnucleome.org/files-processed/4DNFIR94OF6S/).

**Dot detection**

We used dots indicative of loops called in Hi-C data from H1-hESC (https://data.4dnucleome.org/files- processed/4DNFIEEF14ST/) as recently described using our published methods (https://bitbucket.org/creminslab/cremins_lab_loop_calling_pipeline_11_6_2021/src/initial/) {Emerson, 2022 #1935}{Chandrashekar, 2024 #1964}. Using geometric donut-shaped, lower-left, vertical, and horizontal filters (parameters p=2 bins, w=10 bins), we compute an expected interaction frequency for every given bin-bin pair. We computed p-values for each bin-bin pair using a Poisson distribution and corrected for multiple testing using the Benjamini-Hochberg procedure. Final clusters were identified using dynamic false discovery rate (FDR) thresholding.

**SPIN-centric intersection with A/B compartments**

We stratified H1-hESC SPIN states into compartment A or B if they either co-register with a Jaccard index of greater than 1.70 or are embedded within a compartment. All other SPINs partially overlapping compartments were assigned into an 'other' category.

**TAD/subTAD-centric intersection with SPIN states**

We utilized H1-hESC dot and dotless TAD/subTADs previously described (https://data.4dnucleome.org/files-processed/4DNFIW5EIIO2/ and https://data.4dnucleome.org/files-processed/4DNFIU7GTTMW/) {Emerson, 2022 #1935}. We classified dot TAD/subTADs as those in which loops intersect the midpoint (i.e. apex of the TAD/subTAD triangle) +/- 20% the size of the domain. All remaining TAD/subTADs were assigned as dotless. We then intersected dot and dotless TAD/subTADs with SPIN states. All those with a Jaccard index of greater than 0.70 were stratified as co-registering or residing within a SPIN state. We then further stratified dot and dotless TADs co-registered/embedded within SPINs into those that were further co-registered/embedded within A or B compartments with Jaccard index of greater than 0.70. All TADs/subTADs not co-registered/embedded in SPINs nor co-registered/embedded in compartments were assigned into an 'other' category.

**RNA-seq and iMARGI**

We used RNA-seq (https://www.encodeproject.org/experiments/ENCSR537BCG/) and iMARGI (http://sysbiocomp.ucsd.edu/public/wenxingzhao/H1_new/05-09-2019_Tri_iMARGI_H1- control.RNA.minus.bw and http://sysbiocomp.ucsd.edu/public/wenxingzhao/H1_new/05-09-2019_Tri_iMARGI_H1-control.RNA.plus.bw) data generated in h1ESCs. iMARGI's RNA-end reads represent nascent transcripts (https://doi.org/10.1101/2021.06.10.447969). The RNA-end and DNA-end reads were processed with iMARGI-Docker (https://doi.org/10.1038/s41596-019-0229-4) and mapped to the human hg38 reference genome and processed as described. See data availability through the 4DN portal below.

**High-resolution 16 fraction Repli-seq**

We used 16 fraction Repli-seq data from h1ESCs (https://data.4dnucleome.org/experiment-sets/4DNESXRBILXJ/) processed as described into normalized and scaled data arrays (https://data.4dnucleome.org/files-processed/4DNFI3N8GHKR/) {Emerson, 2022 #1935}. We identified initiation zones (https://data.4dnucleome.org/files-processed/4DNFIRF7WZ3H/) as described {Emerson, 2022 #1935}. Raw counts in each fraction ($S_{i,j}$) were normalized by sequencing depth by virtue of read per million (RPM) such that $S_{norm,j,50kb\_bin} = S_j, 50_{kb\_bin} / S_{i,j}$ * 1e6. Repli-seq arrays were subsequently constructed from RPM bedgraphs to form 16 rows with each row representing an S phase fraction and each column representing a 50 kb bin. The array was smoothed by applying a Gaussian filter and scaled such that each column sums to 100.

**Data Visualizations**

We visualized h1ESCv2.5 Hi-C (https://data.4dnucleome.org/files-processed/4DNFI82R42AD/) counts at SPIN state calls or dot and dotless TADs/subTADs by adding 60% of the size of the domain or SPIN to the edges of the maps and stretching to a defined length L. Each domain or SPIN was used once in the visualization and the counts in every pixel were normalized by mean distance-dependence expected value and then averaged across all 2D matrices. We similarly visualized high-resolution 16 fraction Repli-seq, total RNA-seq, iMARGI, and compartment eigenvectors resized to the same intervals as TADs, subTADs, or SPIN states. Signal for high- resolution 16 fraction Repli-seq is the average pileup, total RNA-seq and iMARGI is the median pileup of averaged plus and minus strands. The compartment eigenvector is the mean pileup signal. We resized to defined length L with the resize() method in OpenCV image package (https://pypi.org/project/opencv-python/).

**Initiation Zone Resampling Test**

We computed the proportion of early and late IZs intersecting with Dot TAD/subTAD boundaries (https://data.4dnucleome.org/files-processed/4DNFIWNJ5RR7/), Dotless boundaries (https://data.4dnucleome.org/files-processed/4DNFIT6QE9YU/), and no boundaries. To create a null set of IZs, we computationally sampled the genome for random intervals matched by number, size, and A/B compartment distribution of real early or late IZs. We created a null distribution by sampling $1.5x10_8$ times and computing a one-tailed empirical pvalue as the area under the null distribution to the right of the real IZs. We used only null and real IZ sets in autosomal regions with sufficient counts for the statistical test by filtering unmappable telomeric/centromeric regions.

**Chromatin dataset processing**

CUT&Run datasetx have been processed by trimming adaptors using cutadapt, locally mapping the reads using bowtie2, filtering for quality, removing duplicates and ENCODE blacklisted regions (ENCFF419RSJ) using samtools and computing the coverage using deeptools. Average chromatin landscape at IZs has been computed using HOMER on a 1Mb region centered on each IZ centre and plotted using R. Chromatin profile has been plotted using the WashU browser and IGV on chr2:20,404,583-58,108,703.

**Initiation Zone integration with chromatin and transcription**

HCT116, H1ESC and mESC IZs have been grouped by replication timing by splitting the corresponding Repli-seq data into quartiles. Matching random regions have been generated by shuffling the IZs regions within their respective replication timing quartile using bedtools. Insulation score at IZs and random regions has been computed by extracting the minimum insulation score in each IZ or random region using bedtools. Accessibility at IZs and random region has been computed by extracting the total ATAC-seq signal at each IZ or random region using bedtools. H1ESC RNA-seq data have been processed by mapping reads on hg38 using Hisat2, filtering for quality using samtools and computing the coverage using bedtools. Expression at IZs and random regions has been computed by extracting the total RNA-seq coverage (adding plus and minus strands for the H1ESC RNA-seq) at each IZ or random region using bedtools. H1ESC Cut&Run chromatin marks dataset have been processed by trimming adaptors using cutadapt, locally mapping the reads using bowtie2, filtering for quality, removing duplicates and ENCODE blacklisted regions using samtools and computing the coverage using deeptools. Average chromatin marks at IZs and random region has been computed using HOMER on a 1Mb region centered on each IZ centre. Figures have been plotted using R.

**Data accessibility**

**Hi-C**

- h1ESC Hi-C 2.5 source: https://data.4dnucleome.org/files-processed/4DNFI82R42AD/
- h1ESC dot domains: https://data.4dnucleome.org/files-processed/4DNFIW5EIIO2/
- h1ESC dotless domains: https://data.4dnucleome.org/files-processed/4DNFIU7GTTMW/
- h1ESC dot boundaries: https://data.4dnucleome.org/files-processed/4DNFIWNJ5RR7/
- h1ESC dotless boundaries: https://data.4dnucleome.org/files-processed/4DNFIT6QE9YU/
- h1ESC loops: https://data.4dnucleome.org/files-processed/4DNFIEEF14ST/
- h1ESC TADs/subTADs: https://data.4dnucleome.org/files-processed/4DNFIR94OF6S/

**16 fraction Repli-seq**

- h1ESC raw fastq: https://data.4dnucleome.org/experiment-sets/4DNESXRBILXJ/
- h1ESC read depth scaled normalized array for IZ calls & visualization: https://data.4dnucleome.org/files-processed/4DNFI3N8GHKR/
- h1ESC Early, Early-mid, Late IZs on read depth normalized: https://data.4dnucleome.org/files-processed/4DNFIRF7WZ3H

**RNA**

- h1ESC RNA-seq source: https://www.encodeproject.org/experiments/ENCSR537BCG/
- h1ESC RNA-seq processed file: https://www.encodeproject.org/files/ENCFF584VXW/ (plus strand signal of unique reads) https://www.encodeproject.org/files/ENCFF307LLA/ (minus strand signal of unique reads)
- h1ESC raw files: http://sysbiocomp.ucsd.edu/public/wenxingzhao/H1_new/05-09-2019_Tri_iMARGI_H1-control.RNA.minus.bw;

http://sysbiocomp.ucsd.edu/public/wenxingzhao/H1_new/05-09-2019_Tri_iMARGI_H1- control.RNA.plus.bw

**SPIN states**

- h1ESC and HFFc6 SPIN states are in Supplemental Table 1.

**h1ESC**

- h1ESC A/B Compartments: https://data.4dnucleome.org/files-processed/4DNFIOUDCJRH/

**Contact map**

- H1ESC: https://data.4dnucleome.org/files-processed/4DNFIIMZB6Y9/

**ATAC-seq**

- HCT116: https://www.encodeproject.org/files/ENCFF624HRW/
- mESC: https://data.4dnucleome.org/files-processed/4DNFI6HY3NE7/
- H1ESC: https://data.4dnucleome.org/files-processed/4DNFICPNO4M5/

**Histones marks**

- H1ESC, H2AZ: https://data.4dnucleome.org/experiments-seq/4DNEXHDA1L74/ (reanalyzed)
- H1ESC, H3K27AC: https://data.4dnucleome.org/experiments-seq/4DNEX5EQJ2P2/ (reanalyzed)
- H1ESC, H3K4me3: https://data.4dnucleome.org/experiment-set-replicates/4DNESPE6J9FU
- H1ESC, H3K4me1: https://data.4dnucleome.org/experiments-seq/4DNEXOKGNAEQ/ (reanalyzed)
- HCT116, H2AZ: https://data.4dnucleome.org/files-processed/4DNFIPHO57I2/
- HCT116, H3K27Ac: https://data.4dnucleome.org/files-processed/4DNFIYNC2EO5/
- HCT116, H3K4me3: https://data.4dnucleome.org/files-processed/4DNFIT9DN6JI/
- mESC, H3K27Ac: https://data.4dnucleome.org/files-processed/4DNFIXE23VC7/
- mESC, H3K4me3: https://data.4dnucleome.org/files-processed/4DNFIQYJLKKH/
- mESC H2AZ: https://www.ncbi.nlm.nih.gov/geo/query/acc.cgi?acc=GSM4802378 (reanalyzed)

**Gene expression**

- HCT116: https://www.encodeproject.org/files/ENCFF766TYC/
- mESC: https://data.4dnucleome.org/files-processed/4DNFI4XVSIFH/

**IZs**

| | | | |
|---|---|---|---|
| H1ESC | IZs | 4DNFI5OBN63G | https://data.4dnucleome.org/files-processed/4DNFI5OBN63G |
| HCT116 | IZs | 4DNFIYO3H24N | https://data.4dnucleome.org/files-processed/4DNFILNNSFMD/ |
| mESC | IZs | 4DNFI53E1A38 | https://data.4dnucleome.org/files-processed/4DNFI53E1A38/ |

**2-fractions repli-seq**

- H1ESC: https://data.4dnucleome.org/experiments-seq/4DNFIWSH27RS ; https://data.4dnucleome.org/experiments-seq/4DNFIQ6F59NT
- HCT116: https://data.4dnucleome.org/files-processed/4DNFIM9S18WO/ ; https://data.4dnucleome.org/files-processed/4DNFIC4VUF86
- mESC: https://data.4dnucleome.org/files-processed/4DNFIPB7M5B6 ; https://data.4dnucleome.org/files-processed/4DNFIRHK7RZF

**Insulation score**

- H1ESC: https://data.4dnucleome.org/files-processed/4DNFIUK3UVZX
- HCT116: https://data.4dnucleome.org/files-processed/4DNFIGKFF445/
- mESC: https://data.4dnucleome.org/files-processed/4DNFI2LE1KZL/

**iMARGI datasets**

- h1ESC: https://data.4dnucleome.org/experiment-set-replicates/4DNESNOJ7HY7/

### 7 Methods for predicting Hi-C maps from sequence (Akita)

Two convolutional neural network models with the Akita architecture {Fudenberg, 2020 #1938} were trained, one on H1-hESC and one on HFFc6 Micro-C data. Micro-C data was downloaded from the 4DN data portal {Reiff, 2022 #1966} and processed as previously described {Fudenberg, 2020 #1938}. The coordinates of the deletion at the TAL1 locus with experimentally verified changes on genome folding was obtained from Hnisz *et al*. {Hnisz, 2016 #1374}. The deletion was centered and systematically extended on both sides to get the input sequence for Akita ($2^{20}$ bp).

FIMO {Grant, 2011 #1967} was used to scan the human genome (hg38) to identify the potential binding sites (p-value < 1e-5) of cell type specific transcription factors POU2F1::SOX2 (MA1962.1) and FOSL1::JUND (MA1142.1) using their annotations in the JASPAR database {Castro-Mondragon, 2022 #1968}. TAD boundaries in the Micro-C datasets at 5 kb resolution were downloaded from the 4DN data portal. The ones that were not shared between the cell types (no boundary in the other cell type within 20 kb) were identified as cell-type-specific TAD boundaries. The binding sites overlapping cell-type-specific TAD boundaries of each cell type were extracted using bedtools {Quinlan, 2010 #1969}. *In-silico* mutagenesis was performed on the resulting binding sites by replacing the motifs with random sequences to evaluate their effects on predicted genome folding. Motif logos were visualized using the contribution scores of the motifs, which were computed by DeepExplainer (DeepSHAP implementation of DeepLIFT) {Avsec, 2021 #1946}{Shrikumar, 2019 #1971}.
